# Lipid- and protein-directed photosensitizer proximity labeling captures the cholesterol interactome

**DOI:** 10.1101/2024.08.20.608660

**Authors:** Andrew P. Becker, Elijah Biletch, John Paul Kennelly, Ashley R. Julio, Miranda Villaneuva, Rohith T. Nagari, Daniel W. Turner, Nikolas R. Burton, Tomoyuki Fukuta, Liujuan Cui, Xu Xiao, Soon-Gook Hong, Julia J. Mack, Peter Tontonoz, Keriann M. Backus

## Abstract

The physical properties of cellular membranes, including fluidity and function, are influenced by protein and lipid interactions. In situ labeling chemistries, most notably proximity-labeling interactomics are well suited to characterize these dynamic and often fleeting interactions. Established methods require distinct chemistries for proteins and lipids, which limits the scope of such studies. Here we establish a singlet-oxygen-based photocatalytic proximity labeling platform (POCA) that reports intracellular interactomes for both proteins and lipids with tight spatiotemporal resolution using cell-penetrant photosensitizer reagents. Using both physiologically relevant lipoprotein-complexed probe delivery and genetic manipulation of cellular cholesterol handling machinery, cholesterol-directed POCA captured established and unprecedented cholesterol binding proteins, including protein complexes sensitive to intracellular cholesterol levels and proteins uniquely captured by lipoprotein uptake. Protein-directed POCA accurately mapped known intracellular membrane complexes, defined sterol-dependent changes to the non-vesicular cholesterol transport protein interactome, and captured state-dependent changes in the interactome of the cholesterol transport protein Aster-B. More broadly, we find that POCA is a versatile interactomics platform that is straightforward to implement, using the readily available HaloTag system, and fulfills unmet needs in intracellular singlet oxygen-based proximity labeling proteomics. Thus, we expect widespread utility for POCA across a range of interactome applications, spanning imaging to proteomics.

## Introduction

Cholesterol is required for cellular and organismal health. As a key component of cellular membranes, cholesterol functions to regulate membrane fluidity, curvature^1^, and the creation of discrete membrane structures such as lipid rafts^2^. Beyond organelle-based compartmentalization of cellular processes, cholesterol and related metabolites, such as bile acids^3^ and oxysterols^4^, also function as key signaling molecules. Further illustrating cholesterol’s importance, many human diseases are associated with aberrant cholesterol levels, including common cardiovascular diseases such as atherosclerosis^5^ as well as rare monogenic disorders, including familial hypercholesterolemia, Niemann-Pick diseases^6^ and Smith-Lemli-Opitz syndrome^7,8^.

Due to cholesterol’s hydrophobicity and essential functions in cellular physiology, animals have evolved tightly regulated processes to sense cholesterol and facilitate sterol movement between and within cells^9^. Intracellularly, expression of cholesterol biosynthesis and transport proteins are tightly controlled by the cholesterol sensing activity of the sterol response element-binding proteins (SREBPs) and SREBP cleavage-activating protein (SCAP). Extracellularly, the high- and low-density lipoproteins (HDL and LDL) are the primary cholesterol delivery mechanism between cells and organs with the body^10^, with cholesterol transport to peripheral tissues primarily via LDL and delivery to the liver for excretion largely by HDL^11^. Lipoprotein particles undergo selective uptake after binding to key plasma membrane receptors, most notably low-density lipoprotein receptor (LDLR) and scavenger receptor class B member 1 (SR-B1) for LDL and HDL, respectively. LDL binding to LDLR results in clathrin-mediated endocytosis^12,13^ followed by the release of free cholesterol in lysosomes and subsequent transport to the endoplasmic reticulum (ER) regulated by the Niemann-Pick like protein (NPC). HDL-cholesterol (HDL-C) delivery to cells is predominantly mediated through SR-B1, where both non-vesicular^14^ and endocytotic^15^ transport pathways are controlled by SR-B1. For non-vesicular selective uptake, SR-B1 forms a channel that delivers cholesterol (primarily esterified cholesterol, although the channel can also accommodate free cholesterol^16^ directly to the plasma membrane (PM)^17^ prior to transport to the ER.

While the ER is the primary cholesterol sensing organelle^18–20^, it is comparatively cholesterol poor, and most cellular cholesterol (>90%) is found in the PM^21^. Thus, defining the proteins responsible for the dynamic and bidirectional movement of cholesterol between the PM and the ER remains a central challenge for understanding mechanisms governing cholesterol homeostasis. One recently described mechanism involves the Aster family of proteins (Aster-A, -B and -C, also known as GRAMD1 proteins), a family of cholesterol binding proteins that shuttles cholesterol from the PM to the ER^22–24^.

Cholesterol binding proteins (CBPs) have classically been identified via binding studies enabled by limited proteolysis, detergent extractability assays, and radiolabeled or photoactivatable cholesterol^25–27^. More recently, mass-spectrometry-based proteomic approaches have emerged as a favored strategy for high-throughput and proteome-wide delineation of CBPs. Illustrating the power of these global interactomic studies, recent chemoproteomic efforts have used diazirine and alkyne functionalized cholesterol probes^28^ to profile CBPs in a proteome-wide manner^29^. This approach has been applied to map sterol binding sites in VDAC1^30^ and TMEM97^31^. Despite their widespread utility, diazirine probes have drawbacks, including low overall labeling efficiency and biased reactivity towards polar and acidic amino acid residues^32^ that typically are depleted in transmembrane domains^33^. Within the context of mapping CBPs, the primary delivery mechanism for cholesterol diazirine probes has been via complexation with methyl-beta-cyclodextrin (mβCD) which, while not physiological, allows for delivery of large amounts of probe required for efficient capture of interacting proteins. Recent work has demonstrated that diazirine probes can be delivered to cells when complexed with lipoproteins^34^; however, whether this delivery method is compatible with chemoproteomic applications of cholesterol probes remains to be seen.

The advent of proximity labeling proteomic (PLP) methods including the APEX^35^ and BioID^36^ platforms that harness enzymatic biotinylation and, more recently, photocatalytic PLP methods that trap interactions with reactive species activated by photocatalysts^37–44^ all effectively label proximal proteins for downstream identification. Recent work has shown that singlet oxygen (^1^O_2_)-generating photosensitizers can also be used for proximity labeling-based capture of protein interactors, with ^1^O_2_-labeled proteins trapped using nucleophilic capture reagents^43–53^. Photosensitizer-based platforms offer the potential added benefits of comparatively straightforward reagent synthesis and a highly-reactive labeling species, which enables short labeling times and spatially-restricted labeling^54–56^. By enabling reduced probe concentrations compared to diazirine probes and by defining both direct binders and the local protein environment, photosensitizer-based platforms are well suited to define cholesterol interactomes, including proteins sensitive to changes to membrane composition and fluidity. However, a key limitation of photosensitizer-based protein capture platforms is a general lack of methods compatible with intracellular labeling. This gap can be largely ascribed to the poor membrane permeability and high background labeling for most photosensitizers utilized—however, several recent reports, including the RNA-capture method Halo-seq^45,57^, as well as studies using unconjugated free photosensitizers^58,59^ provide evidence in support of the tractability of intracellular photosensitizer-based PLP.

Here we establish an intracellular ^1^O_2_-based interactomic platform called <u>p</u>hotosensitizer-dependent <u>o</u>xidation and <u>c</u>apture by <u>a</u>mine (POCA) that pairs amine-based protein capture with cell-penetrant photosensitizer reagents. POCA using a cholesterol-photosensitizer probe (termed chol-POCA) faithfully captures known and novel cholesterol interactors, with physiologically relevant lipoprotein delivery in both primary hepatocytes and immortalized cell lines. By pairing POCA with the HaloTag protein system^60^ (termed Halo-POCA), we demonstrate POCA’s versatility for both lipid and protein interactomes. Halo-POCA using HaloTag-Aster fusion proteins confirmed domain-specific oligomerization^23^ and identified the lipid raft protein flotillin-1 (FLOT1) as colocalizing with Aster. Taken together these studies showcase POCA’s capacity to delineate physiologically relevant proteins involved in cholesterol movement within cells. Illustrating the broader impact of POCA beyond cholesterol biology, we also reveal that protein-based Halo-POCA faithfully captured nearly all subunits of the nuclear pore complex (NPC). Thus, our work establishes POCA as a highly versatile platform with widespread applications in defining both the protein and lipid interactomes.

## RESULTS

### JF_570_ is a suitable Photosensitizer for intracellular singlet-oxygen-based photocatalytic proximity labeling proteomics

Our first step to establish our envisioned POCA ^1^O_2_-based protein interaction platform (**Figure 1A** and **Figure S1**) was the selection of a cell-permeable photosensitizer to enable intracellular photocatalytic proximity labeling. Inspired by recent reports utilizing bis-acetylated (Ac_2_) 4’,5’-dibromofluorescein (DBF) (Ac_2_DBF, Scheme S1), including the RNA-capture method Halo-seq^45,57^, we first investigated DBF as the photosensitizer in the POCA workflow. We synthesized two reagents, DBF-cholesterol probe S7 obtained in 5% overall yield with a 5-step longest linear sequence from 6-carboxy-Ac_2_-DBF (intermediate S1, Scheme S2), and the previously reported^45^ bis-acetylated DBF-HaloTag ligand (Ac_2_DBF-HTL, Scheme S1). Application of mβCD-complexed S7 to cells followed by light irradiation in the presence of propargylamine (PA) afforded no detectable protein labeling (**Figure S2A, B**). In contrast, when cells transiently expressing murine Aster-A fused to HaloTag (HaloTag-mAster-A) were treated with Ac_2_DBF-HTL followed by irradiation and PA capture, we observed substantial labeling as visualized by in-gel fluorescence analysis (**Figure S2C, D, E**). Labeling studies using recombinant BSA further confirmed a histidine-based labeling mechanism (**Figure S3** and **Figure S4**), as previously reported^48,49,55^. However, the high HaloTag-independent background labeling (**Figure S2E**) indicated the likelihood of poor signal-to-noise for protein-based capture. Taken together, these data indicated to us that DBF did not have suitable physicochemical properties for achieving high signal-to-noise intracellular protein proximity labeling.

**Figure 1.**
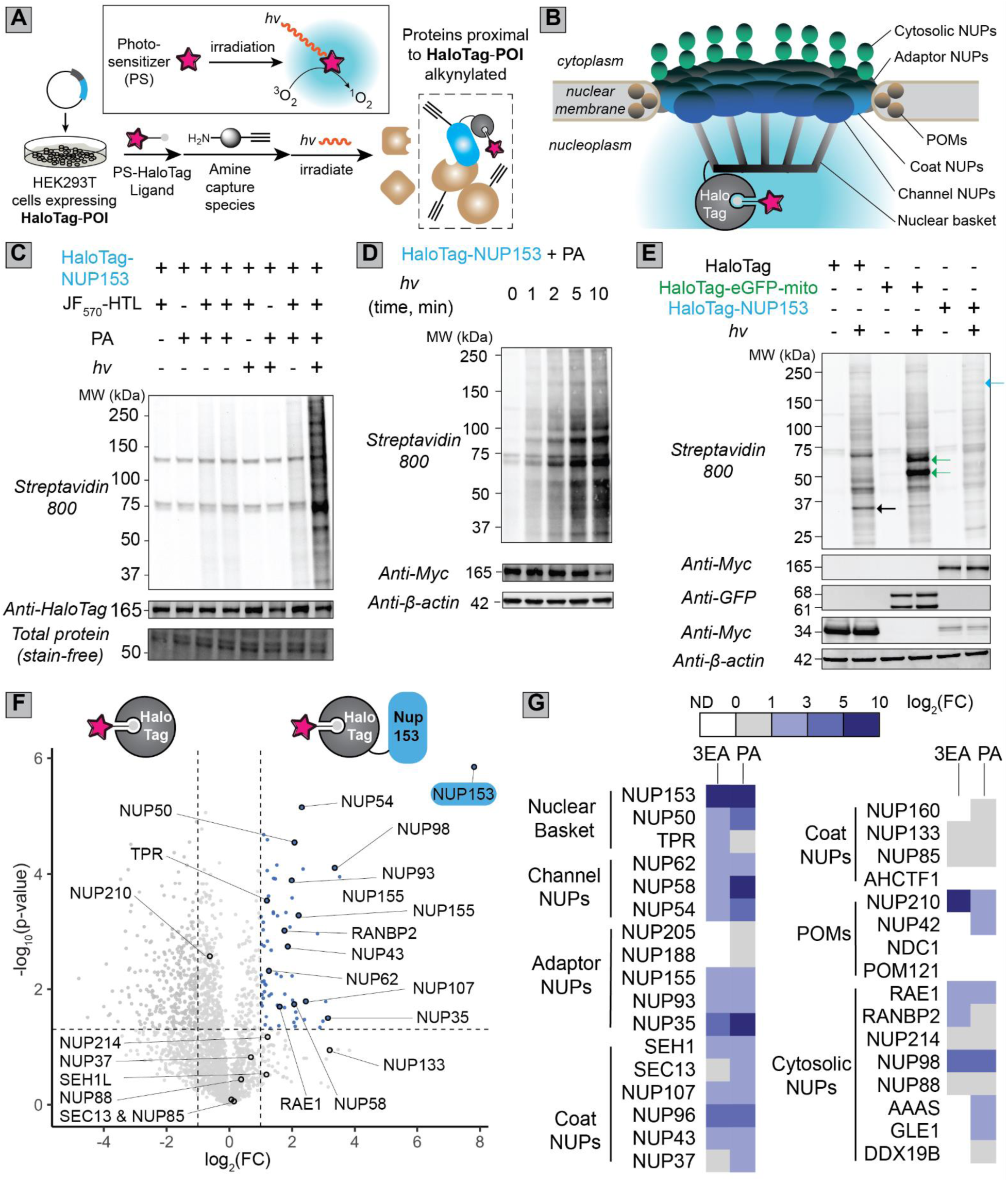
Establishing the Halo-POCA system. **(A)** General workflow for Halo-POCA in which cells overexpressing HaloTag fusion proteins are labeled with photosensitizer (PS)-functionalized HaloTag ligand, irradiated in the presence of amine trapping agent and subjected to click-chemistry enabled capture and proteomic analysis. Protein of interest (POI). **(B)** Schematic diagram of the nuclear pore complex (NPC), including the HaloTag-NUP153 bait protein used for POCA analysis. **(C-E)** Gel-based analysis of POCA labeling performed following the workflows **Figure 1** and **Figure S1A** for HEK239T cells transiently expressing the indicated HaloTag fusions treated with JF_570_-HTL (100 nM, 10 min) followed by media washouts, addition of PA (10 mM) and irradiation (15 W yellow LED, 170,000 Lux max intensity, on ice), lysis, click conjugation to biotin-azide, SDS-PAGE, and Streptavidin blot. For ‘C’, total protein was assessed by stain-free blot. ‘C’ and ‘E’ were subjected to 5 min light (*hv)* irradiation, whereas in ‘D’ irradiation time was varied as indicated. For ‘E’ colored arrows indicate biotinylated proteins corresponding to the expected MW of the color-matched bait proteins. **(F, G)** Mass spectrometry-based POCA analysis generated following the workflows in **Figure 1** and **Figure S1A** for HEK293T cells overexpressing either HaloTag-NUP153 or free HaloTag. POCA labeling was performed as described for the gel-based analysis with interactors trapped using 3-ethynylaniline (3EA, 10 mM) or propargylamine (PA, 10 mM) comparing relative POCA labeling with transiently overexpressed HaloTag-NUP153 versus free HaloTag in HEK293T cells. For ‘F’ volcano plot shows all identified proteins for EA labeling, with all nucleoporins circled and proteins significantly enriched (log_2_(fold-change)>1, *p*<0.05) with HaloTag-NUP153 highlighted in blue. For ‘G’ heatmap shows all identified nucleoporins colored based on LFQ-MBR^63^ log_2_(FC) values, *p*<0.05 for all proteins except those labeled gray. POM: pore membrane protein. For statistical significance in ‘F’ and ‘G’, variances were calculated for each sample-condition pairing and a corresponding two-tailed t-test was performed to generate p-values. n=3 per group for MS datasets. Pore membrane proteins (POMs). All MS data can be found in **Data S1**.

We hypothesized that a net neutral, bioavailable photosensitizer could simultaneously reduce the background labeling and improve cellular uptake. We turned towards the Janelia-Fluor^61^ series of dyes, which are bioavailable and have enhanced photostability, a property we envisioned would be desirable due to the intentional production of ^1^O_2_.The recently reported photosensitizer JF_57062_ fulfilled our criteria of net neutral charge and low background in cells, as demonstrated by its prior use for chromophore-assisted light inactivation of proteins, cell ablation, and photopolymerization of an electron microscopy contrast agent. Motivated by this precedent, we sourced JF_570_-HaloTag ligand (JF_570_-HTL). Gratifyingly, using Halo-POCA with a gel- and click-chemistry based photocatalytic alkynylation-based readout, we observed robust protein-dependent labeling with negligible background labeling (**Figure S5**), consistent with high specificity.

### Halo-POCA using JF_570_ enables high coverage identification of components of the nuclear pore complex

We next set out to further vet the POCA platform with the goals of establishing optimal treatment conditions and ensuring that the method faithfully captures bona fide interactors. To address both of these goals, we applied the POCA platform to the widely-studied and well-characterized nuclear pore complex (NPC)^64^, responsible for the transport of protein and RNA cargo to and from the nucleus. The constituent proteins of the NPC, known as nucleoporins (NUPs), have previously been investigated^65^ using proximity-based labeling with BioID^36^. We generated a HaloTag fusion protein for nuclear pore complex protein Nup153 (NUP153), a member of the nuclear basket, which localizes to the nuclear face of the pore (**Figure 1B**). Gratifyingly, we observed robust JF_570_-HTL-, PA-, and light-dependent labeling in gel-based analysis of protein alkynylation (**Figure 1C**). With the exception of the endogenously biotinylated 75 kDa and 125 kDa proteins, negligible streptavidin binding was observed in the absence of all reaction components. Protein labeling was observed to be dose-dependent with respect to PA concentration (**Figure S6**) and irradiation time (**Figure 1D** and **Figure S7**). No noticeable decrease in cell viability was observed after short exposures to PA (**Figure S8**). Akin to the rapid kinetics of APEX^35^, POCA labeling was observed to occur rapidly, with irradiation for merely one minute affording marked protein biotinylation (**Figure 1D** and **Figure S7**). Illustrating the generalizability of Halo-POCA, distinct labeling patterns were observed with gel-based photocatalytic alkynylation analysis for each bait protein analyzed, including HaloTag-mAster-B (**Figure S5**), free HaloTag, HaloTag-NUP153, and eGFP-HaloTag fusion (HaloTag-eGFP-mito) (**Figure 1E**) which is localized to the mitochondrial outer membrane (MOM) by a targeting sequence from *Listeria monocytogenes*^66^. Importantly, bands consistent with the molecular weights of each bait protein were also observed to be biotinylated, which provides further evidence of the proximity-dependent labeling mechanism (**Figure 1E**). Click-enabled imaging analysis with HaloTag-eGFP-mito further corroborated colocalization of biotin, eGFP, and HTL signals (**Figure S9**), with negligible background in the absence of PA (**Figure S9B**). Similar colocalization of biotin and HTL was observed when HaloTag-NUP153 was used (**Figure S10**).

Guided by the encouraging performance of Halo-POCA in gel-based and imaging analysis, we next opted to vet POCA’s capture of bona fide interactors. We find that POCA proteomic analysis identified ∼3000 unique proteins for each bait protein analyzed (**Figure S11**). We then opted to focus on the NPC as a test case, due to its well-defined composition. We compared the POCA interactomes for cells overexpressing either free HaloTag protein or HaloTag-NUP153, using label free quantitation-match between runs (LFQ-MBR) enabled by IonQuant^63^ analysis with MSBooster^67^ in the MSFragger^68^ search software (**Figure S12**). Gratifyingly a total of 228 proteins, including 20 of the 33 UniProtKB-annotated nucleoporins, were preferentially enriched (log_2_(FC) > 1, p-value < 0.05) using Halo-POCA followed by MS analysis (**Figure 1F**,**G**, **Figure S13**, and **Data S1**). All members of the nuclear basket and nuclear channel were enriched in addition to channel, adaptor, and coat NUPs of the symmetric core (**Figure 1G** and **Figure S14**), which provides further evidence in support of POCA’s capacity to map protein complexes.

The singlet oxygen generating protein miniSOG can also be used for ^1^O_2_-based PLP, and increased labeling has been reported for 3-ethynylaniline (3EA, Scheme S1) compared to PA^6949^. Therefore, we also compared the performance of 3EA to PA using our NPC test system (**Figure 1F**, **G**). For 3EA labeling, we identified 68 total proteins, including 15 total nucleoporins, that were preferentially captured by HaloTag-NUP153 (**Figure 1F**, **Figure S14**, and **Data S1**). While our PA dataset had modestly higher overall coverage, the substantial overlap for the enriched nuclear pore components supports that PA and 3EA can be used relatively interchangeably (**Figure 1G**, **Figure S13**, and **Figure S14**). In total, our PA and 3EA analysis enriched 23/33 (70%) of NUPs (**Figure 1G** and Data S1). Notably, 7 additional NUPS were identified but did not pass LFQ enrichment criteria. Only AHCTF1/ELYS, NDC1, and POM121 were not detected using Halo-POCA. Further illustrating the robust performance of the POCA platform, the coverage of NUPs obtained using a single bait protein and 5 minute labeling time with Halo-POCA is similar to the previous report using BioID, which used six different NUP-BirA fusions to probe the organization of the NPC^65^.

### POCA identifies known cholesterol binding proteins

Encouraged by the favorable performance of the Halo-POCA system, we next returned to one of our key overarching objectives, namely mapping the cholesterol-protein interactome. To establish a cholesterol-POCA (chol-POCA) system (**Figure 2A**), we synthesized both cholesterol-JF_570_ probe 1 and palmitate-JF_570_ probe 2 (**Figure 2B**), with the latter compound designed to function as a control for non-specific protein capture. Both compounds were obtained in modest yields (Scheme S3 and Scheme S4) by amide coupling of JF_570_ to either the tail end of 3*β*-hydroxy-5-cholenoic acid or palmitic acid. A polyethylene glycol-based spacer was incorporated to extend the photosensitizer away from the biomolecule by about ∼22 Å (Scheme S3 and Scheme S4). We then subjected cells to POCA analysis with mβCD-1 or mβCD-2 following the workflows shown in **Figure S15**. Gel-based analysis revealed robust compound-, light-, and amine-dependent labeling (**Figure S16A,B,C**), consistent with probe dependent protein labeling. Intriguingly, palmitate probe 2 showed increased protein labeling when compared to cholesterol probe 1 (**Figure S16D**), suggestive of reagent-dependent differences in cellular uptake and/or protein interactions.

**Figure 2:**
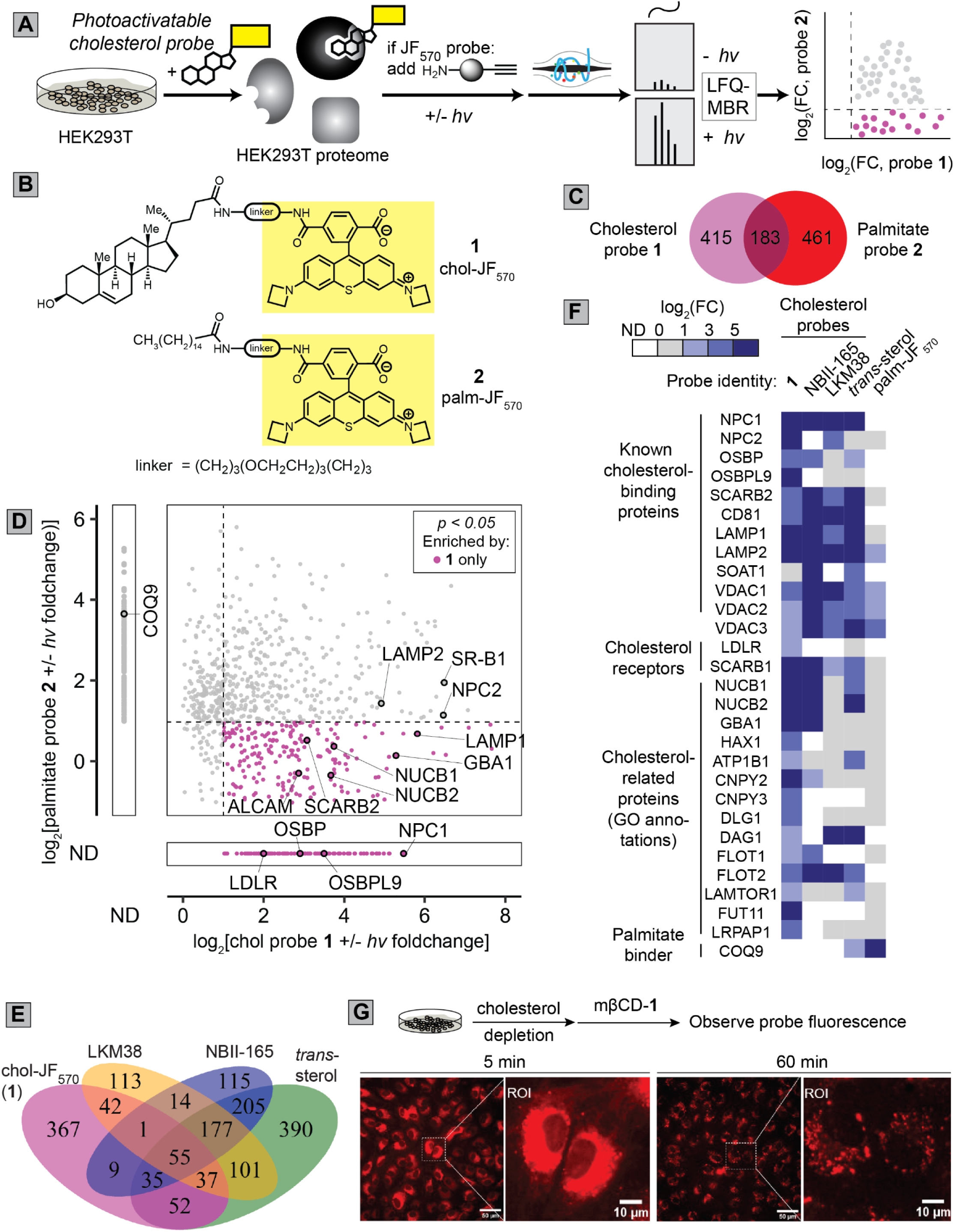
Using POCA with cholesterol identifies known cholesterol-interacting proteins. **(A)** Schematic workflow for analyzing the light-dependent enrichment of different sterol-interacting proteins in cells by photoactivatable mβCD-complexed cholesterol probes. **(B)** Structures of the photosensitizer-containing probes **1** and **2**. (**C-F**) Comparison of the protein targets captured by probe **1**, **2,** and cholesterol diazirine probes NBII-165, LMK38, and *trans-*sterol. Cells were treated with mβCD-complexed probes **1** or **2** (10 μM, 1 h) followed by PA (10 mM) and irradiation with visible light (15 W yellow LED, 170,000 Lux max intensity, on ice, 5 min), or mβCD-complexed NBII-165, LKM38, or *trans*-sterol (10 μM, 1 h) and irradiated with UV light (365 nm, 22 °C, 15 min) for diazirine probes. Following irradiation, lysed samples were subjected to click conjugation to biotin-azide, streptavidin enrichment, tryptic digest, LC-MS/MS analysis, search, and label free quantification (LFQ)^63^ using MSFragger^68^. Enrichment criteria: log_2_(FC) > 1, *p* < 0.05. **(C, D)** Comparison of the unique and shared proteins enriched by mβCD-complexed **1** and **2**, with ‘C’ showing overlap and ‘D’ comparing fold enrichment. **(E)** Overlap of proteins enriched by mβCD-complexed Cholesterol-JF_570_ probe **1** and established cholesterol-diazirine probes. **(F)** Comparison of lipid and sterol-related proteins enriched by each probe. Coloring corresponds to increased LFQ log_2_(FC) values, *p* < 0.05; grey indicates *p* > 0.5. **(G)** Confocal microscopy images of probe fluorescence in human aortic endothelial cells treated with mβCD-**1** over time. Cells were treated with mβCD (2.5 mM, 15 min) to deplete cholesterol, washed, then treated with mβCD-**1** (2 μM, 5 min) before images were taken. Scale bars = 50 μM. For ‘C’-‘F’, variances were calculated for each sample-condition pairing and a corresponding two-tailed t-test was performed to generate *p*-values. n=3 per group. LFQ-MBR: label-free quantification-match between runs; FC: fold change; ND: not detected. All MS data and lists of ‘cholesterol-binding proteins’ and ‘cholesterol-related proteins’ can be found in **Data S2**.

Guided by these findings, we extended our analysis to mass spectrometry-based proteomic capture of 1- and 2-interactors (**Figure 2A**). We identified 644 total proteins significantly enriched by palmitate probe 2 and 589 enriched by the cholesterol probe 1 with 183 (∼30%) of the enriched proteins captured by both probes (**Figure 2B** and **Data S2**). The palmitate probe 2 afforded modestly higher (∼25%) coverage when compared to the cholesterol probe 1 (**Figure S17**). We ascribe this difference in coverage to the palmitate probe’s generally increased labeling activity observed in the gel-based labeling assay (**Figure S16D**). Despite these differences in overall labeling, our head-to-head comparison of the proteins significantly enriched by each probe revealed largely distinct target profiles (**Figure 2C**,**D**). Further illustrating the POCA’s capacity to accurately delineate bona fide interactors, the cholesterol probe 1 faithfully captured a number of known cholesterol binding proteins including NPC intracellular cholesterol transporters 1 & 2 (NPC1^27^ and NPC2^70^), scavenger receptor class B member 2 (SCARB2, aka LIMP-2)^71^, oxysterol-binding protein-related protein 9 (OSBPL9)^72^, oxysterol-binding protein 1 (OSBP)^73^, lysosomal associated membrane glycoproteins 1 and 2 (LAMP1 & LAMP2)^74^, and voltage-dependent anion-selective channel proteins 1 and 2 (VDAC1^30^ and VDAC2^75^) (**Figure 2C**, **2D**, **Figure S18A**, and **Data S2**). We also observed enrichment of known lysosomal cholesterol-modifying proteins, such as Lysosomal acid glucosylceramidase (GBA1, which glucosylates cholesterol^76^) and lysosomal acid lipase/cholesteryl ester hydrolase (LIPA)^77^.

Palmitate probe 2 enriched lipid handling proteins involved in fatty acid metabolism and fatty acid oxidation (**Figure S18B**). Notable examples of proteins enriched by 2 include acyl-coenzyme A thioesterase 2, mitochondrial (ACOT2, *aka* PTE2)^78^, enoyl-CoA hydratase, mitochondrial (ECHS1)^79^, and ubiquinone biosynthesis protein COQ9, mitochondrial (COQ9)^80^, all of which bind to palmitate-like alkyl chains. Providing further evidence of the distinct target profiles of each probe, Kyoto Encyclopedia of Genes and Genomes (KEGG) pathway analysis^81^ revealed almost completely distinct pathways enriched by each POCA dataset, consistent with the two probe’s generally non-overlapping protein targets. Notably, the cholesterol dataset enriched pathways associated with the endoplasmic reticulum, lysosomal function, and protein folding diseases (**Figure S19A**), whereas the palmitate dataset captured many pathways associated with lipid metabolism (**Figure S19B**).

### Cholesterol POCA performs comparably to diazirine-based photoaffinity labeling

To further assess POCA’s performance, we next compared chol-POCA to established diazirine-based photoaffinity labeling chemistry. Prior studies using diazirine-based cholesterol photoaffinity probes have identified 265 total candidate cholesterol-interacting proteins in HeLa cells^29^ and have enabled VDAC-cholesterol interaction site mapping^30^. We obtained the previously reported *trans*-sterol and LKM38 probes and additionally synthesized cholesterol analogue NBII-165 as a closer structural analogue to 1 with substitution at carbon 25 (Scheme S5). Photoaffinity-labeling analysis with each diazirine probe by LFQ-MBR (**Figure 2A**) yielded 1346 total proteins that passed enrichment criteria, including 232 shared proteins (**Figure 2E** and **Figure S20**). For each probe, there were 442 (trans-sterol), 155 (NBII-165), and 124 (LKM38) uniquely enriched proteins (**Figure 2E** and **Figure S20**).

We then compared the proteins that were enriched by this panel of cholesterol-diazirine probes to the proteins enriched by the POCA method employing probe 1. Importantly, 55 total proteins were shared between our POCA enriched targets and the diazirine enriched targets (**Figure 2E**). Included among these shared targets are several cholesterol-interacting proteins. Notable cholesterol binding proteins shared across all probes include NPC1, NPC2, SCARB2, OSBP, VDAC1, VDAC2, CD81 antigen (CD81), LAMP1, and LAMP2, and SR-B1 (**Figure 2F**). Gene ontology (GO) analyses revealed cholesterol binding and sterol binding as the top GO Molecular Function results (**Figure S21A, B**). Lysosomal and ER membrane were the top GO Cellular Component results (**Figure S21C**). Notably, 367 proteins were uniquely enriched by chol-JF_570_ when compared with the diazirine probes. Given that the POCA labeling chemistry should capture proximal proteins in addition to direct interactors, we opted to further parse these candidate cholesterol binders. Within this group of uniquely enriched (**Figure 2F** and **Data S2**) proteins examples with documented roles in cholesterol biology including NADPH:adrenodoxin oxidoreductase, mitochondrial (FDXR)^82^, ERLIN2^83^, and OSBPL9^72^. Pointing towards the capacity of POCA to capture a larger interaction sphere compared to diazirine probes, we also found that 1 uniquely enriched several proteins known to form complexes with cholesterol binding proteins, including Alpha-2-macroglobulin receptor-associated protein (LRPAP1, or RAP), a chaperone for LDLR-associated proteins^84^, and low-density lipoprotein receptor (LDLR) (**Figure 2F**). Taken together, these data provide evidence that chol-POCA captures bona fide cholesterol binding proteins, with coverage comparable to established diazirine methods and with the likely added benefit of capturing protein complexes in addition to direct binders.

Most unesterified cholesterol localizes to the plasma membrane, endocytic recycling compartment, and trans-Golgi network^9,85^. Therefore, we hypothesized that these subcellular compartments should be significantly represented in our chol-POCA datasets, consistent with cholesterol-mediated protein labeling. Using a curated dataset of previously reported^86–90^ protein localization annotations, we first compared the subcellular distribution of enriched proteins to all proteins detected. We observed a higher proportion of ER, Golgi, PM, and surface proteins enriched by cholesterol probe **1** (**Figure S22A, B**), consistent with accumulation of cholesterol probe **1** to these compartments. Analogous analysis of enriched proteins captured by the palmitate probe **2** revealed low coverage of Golgi and ER proteins and, instead, a striking increase in the proportion of mitochondrial-annotated proteins enriched (24%, **Figure S22C**). Gratifyingly, the subcellular distribution of proteins enriched by **1** generally matched that observed for similar analysis of the diazirine probe datasets (**Figure S23**). Providing further evidence of the subcellular distribution of probe **1** as well as POCA’s compatibility with both imaging and proteomics, time dependent imaging analysis of mβCD-**1** revealed rapid probe uptake within 5 minutes (**Figure S24** and **Video S1**) localizing to the plasma membrane and ER, with increased accumulation in lysosomal puncta at longer timepoints (Figure 2G, and **Figure S25**), consistent with KEGG analysis of proteins enriched by mβCD-**1** after a 1 h incubation (**Figure S19A**).

Given JF_570_’s zwitterionic nature and the established accumulation of cationic dyes in the mitochondria^91^, the capture of mitochondrial proteins was not entirely unexpected. Therefore, we next opted to investigate whether specific mitochondrial protein interactors could be observed for each probe. Notably, mitochondrial cholesterol comprises only ∼2-4% of the total cellular cholesterol pool^92^, and both palmitate and cholesterol impact mitochondrial physiology, including respiration, polarization, and fragmentation^93–95^. Analysis of the subcellular annotation of proteins enriched by the two probes (**Figure S22A**) revealed a stark difference in the proportion of mitochondrial proteins enriched by each probe (11% for **1** versus 24% for **2**, **Figure S22B**,**C**). Comparison of the palmitate **2** and cholesterol **1** mitochondrial interactors revealed a sizable proportion 61% (48 in total) of shared binders, suggestive of some degree of non-specificity, together with 31 unique cholesterol binders and 17 unique palmitate binders. Providing further evidence of capture of authentic mitochondrial cholesterol binders, cholesterol **1** uniquely identified syntaxin-17 (STX17), which is a key mitochondrial cholesterol transport protein^96^. More broadly, when compared to the diazirine datasets, JF_570_-cholesterol only showed a slight increase in the capture of mitochondrial proteins (11% versus 9% respectively, **Figure S23**), which provides additional evidence that JF_570_-driven probe accumulation in the mitochondria is likely not the primary driver of chol-POCA protein capture.

### Competitive POCA proteomics further delineates cholesterol-interacting proteins

When compared with diazirine probes, the benefits of POCA’s catalytic labeling mechanism include the aforementioned trapping of protein complexes and increased overall reactivity, which may enable improved identification of tough-to-detect proteins. However, POCA’s larger labeling radius also comes at a potential cost, namely that direct versus indirect interactors are not readily delineated by such light-dependent proteomics. Competition studies, in which pre-treatment with an untagged molecule blocks labeling by the photoprobe, are the gold standard for pinpointing high confidence protein-small molecule interaction events, as has been illustrated previously for many exemplary diazirine probes^29,97,98^, together with more nascent interaction mapping technologies^99^. Therefore, we next incorporated competitive labeling into our POCA workflow (**Figure 3A**).

**Figure 3:**
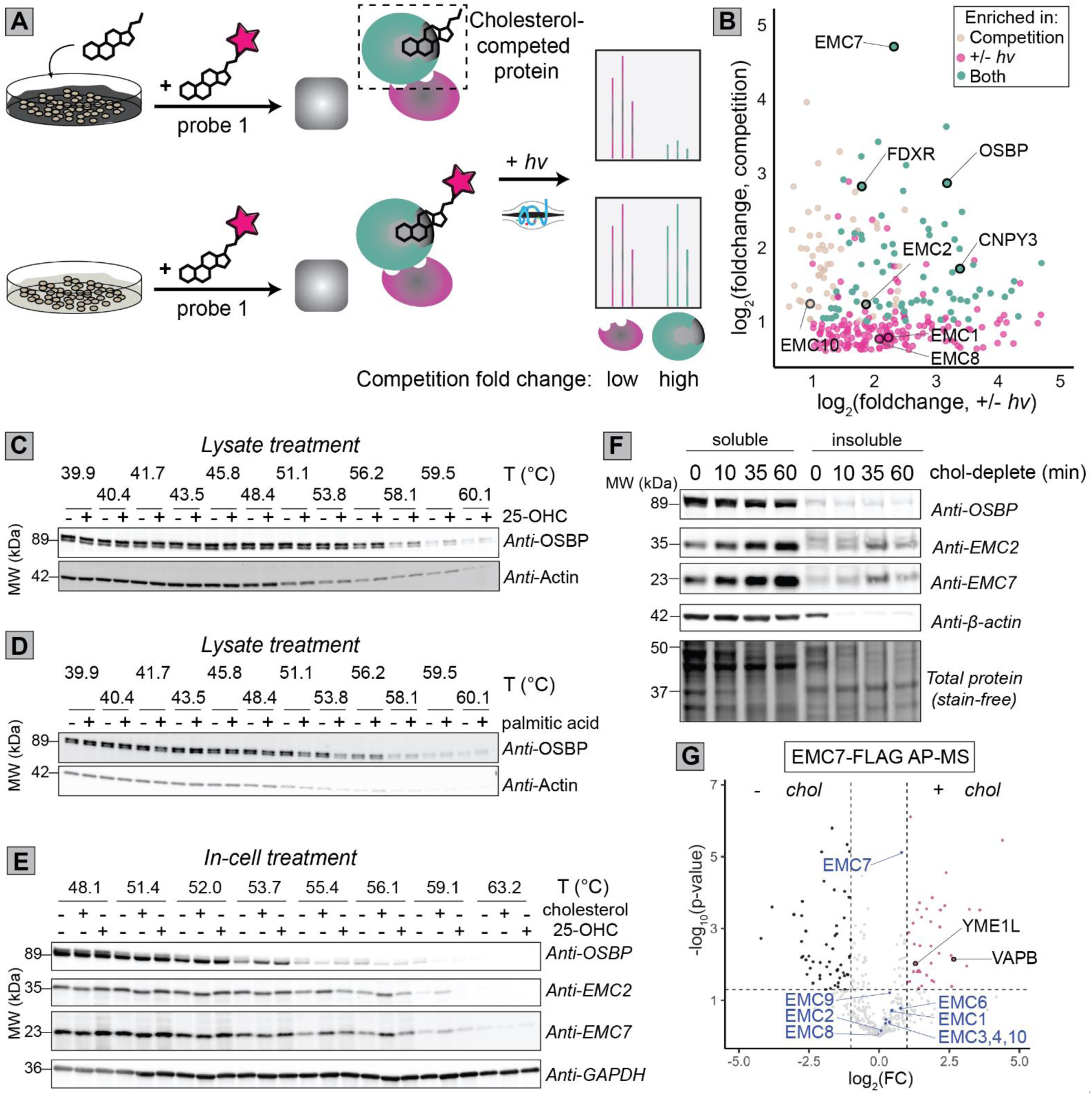
Competition of probe 1 by cholesterol identifies the ER-membrane complex (EMC) as a cholesterol-interacting complex. (**A**) Workflow for cell-based competitive chol-POCA where cells are pre-treated with either vehicle (1 h) or mβCD-complexed cholesterol (100 μM, 30 min) followed by mβCD-**1** (10 μM, 30 min) and subsequent POCA proteomic analysis. (**B**) Comparison of significant (*p*<0.05) protein LFQ log_2_(FC) values for light-dependent enrichment by mβCD-**1** and off-competed labeling by excess cholesterol. (**C, D**) Lysate-based CETSA analysis of HEK293T proteome treated with (C) 25-OHC (10 μM) or (D) palmitic acid (10 μM) and heated (3 min) at the indicated temperatures, followed by centrifugation, and immunoblot analysis n=1. (**E**) Cell-based CETSA analysis of HEK293T cells treated with either mβCD-complexed cholesterol (150 μM, 30 min) or 25-OHC (10 μM, 30 min) and cells heated (3 min) at the indicated temperatures followed by lysis, centrifugation and immunoblot. Blot shown is representative of n=3 independent experiments, see **Figure S28** for all replicates. (**F**) Acute cholesterol depletion with mβCD leads to increased amounts of soluble EMC subunits. HEK293T cells were treated with mβCD (2.5 mM) for the indicated time, harvested, and lysed in the presence of 0.4% NP-40. (**G**) Affinity-purification mass spectrometry analysis identifies VAPB and YME1L as cholesterol-dependent interactors of the EMC. HEK293T cells transiently expressing EMC7-FLAG had cholesterol extracted via treatment with mβCD (2.5 mM, 30 min) or cholesterol loaded by the addition of mβCD-cholesterol (150 10 μM, 30 min) before cells were lysed in the presence of 0.4% NP-40, pull-down with anti-FLAG resin, washed, and bound proteins were digested with trypsin before LC-MS/MS analysis and quantification^63^ of protein intensities. EMC subunits are blue-colored data points, and all fall in the ‘not significant’ portion of the graph. For measures of statistical significance used in parts ‘B’ and ‘G’, variances were calculated for each sample-condition pairing and a corresponding two-tailed t-test was performed to generate p-values. Competition experiments (+/-cholesterol) were performed with 4 replicates (2 biological, 2 technical); labeling experiments (+/-*hv*) used 6 replicates; AP-MS experiments used 6 replicates. 25-OHC: 25-hydroxycholesterol; chol: cholesterol; EMC: ER membrane complex; AP-MS: affinity purification-mass spectrometry. All MS data can be found in **Data S3**.

We subjected cells pre-treated with either vehicle or mβCD-complexed cholesterol to the chol-POCA workflow (**Figure 3A** and **Figure S26**). Gratifyingly, we observed robust cholesterol-mediated blockade of POCA capture for 81 proteins, including the canonical cholesterol-binding protein OSBP and the mitochondrial lipid-handling protein FDXR (**Figure 3B** and **Figure S26**). Intriguingly, we observed that EMC2, EMC7, and EMC10, subunits of the ER membrane complex (EMC), were sensitive to cholesterol competition (**Figure 3B**). Further highlighting the sensitivity of the EMC complex to cholesterol, three additional subunits (EMC1,2, & 8) were also observed to be captured by both probe **1** and several of the diazirine probes in a light dependent manner (**Figure 3B**, **Data S1**). As an ER transmembrane domain insertase^100^, the EMC has an established^101^ role in regulating cholesterol metabolism by functioning to insert the sterol-O-acyltransferase 1 (SOAT1) and squalene synthase (SQS or FDFT1) into the ER membrane. However, to our knowledge whether and how the EMC complex directly responds to intracellular cholesterol levels is unknown.

To test the hypothesis that subunits of the EMC directly engage cholesterol, we turned to the cellular thermal shift assay (CETSA)^102^, which is a widely employed method for detection of protein interactions based on binding-induced changes to protein stability. Recent efforts to define the oxysterol binding proteome illustrate the utility of thermal stability measurements in pinpointing sterol interactors^103^. To establish our cholesterol-directed CETSA assay, we first confirmed that cholesterol-specific stability changes could be detected for a known sterol interactor, namely the oxysterol binding protein OSBP^73^, with mβCD-cholesterol, or soluble 25-hydroxycholesterol (25-OHC). Gratifyingly, and consistent with the prior report of stabilization of oxysterol binding proteins with 25-OHC^103^, we observed stabilization of OSBP by both 25-OHC and cholesterol in lysates (**Figure 3C**). No such appreciable thermal shift was observed for lysates treated with palmitic acid (**Figure 3D**), providing evidence of the specificity of the CETSA assay. Consistent with EMC complex cholesterol interactions, cell-based CETSA revealed both cholesterol and 25-OHC stabilization of EMC7 and EMC2 (**Figure 3E**, and **Figure S27**). Intriguingly, lysate-based CETSA showed no substantial stabilization, suggesting a cell-specific effect (**Figure S28)**. Providing further evidence of the EMC complex dynamically responding to changing cholesterol levels, we observe that acute (mβCD) cholesterol depletion caused a rapid increase in net EMC7 abundance for both cell lines analyze by immunoblot, with a smaller increase observed for EMC2 and no change for OSBP (**Figure 3F** and **Figure S29A**).

Assaying membrane proteins via CETSA often requires addition of low concentrations of detergent during cell lysis^104^. During the course of testing detergent concentrations to establish the EMC CETSA assay, we observed that EMC7 cholesterol-induced stabilization was only detectable with 0.4%, but not the lower concentration 0.2% NP-40 detergent extraction. We postulated that the detergent-dependent differences in cholesterol-EMC stabilization might reflect changes in the membrane-association behavior of EMC7. To test this hypothesis, we compared the amounts of EMC7 extracted by 0.2% versus 0.4% NP-40 after addition of exogenous sterols. We find that *in cellulo* treatment with mβCD-cholesterol but not 25-OHC decreases the extractability of EMC7 at lower detergent concentrations. This decrease occurred within 10 minutes and was specific to EMC7, with no change in abundance observed for OSBP (**Figure S29B**). These data provide further evidence in support of EMC7’s cholesterol sensing activities.

Based on these data, we hypothesized that the EMC7 interactome should be responsive to cellular cholesterol content. To test this hypothesis, we subjected epitope(FLAG)-tagged EMC7 to affinity-purification mass spectrometry (AP-MS) analysis (**Figure S30A**,**B**). Confirming the robustness of our assay, we find that EMC7 robustly enriches most subunits of the EMC complex (**Figure S30B**). In cells loaded with cholesterol, we observed enhanced enrichment of the vesicle-associated membrane protein-associated protein B (VAPB) (Figure 3G, and **Figure S30C,D**), which dynamically organizes mito-ER contact sites in response to nutrient availability^105^, suggesting a cholesterol-dependent role for the EMC in VAPB-mediated tethering. We also observed enhanced binding to the ATP-dependent zinc metalloprotease YME1L1 (YME1L), a protease that degrades mitochondrial proteins such as lipid transfer proteins in response to metabolic needs^106^ in cholesterol loaded cells (**Figure 3G**) and cells not depleted of cholesterol (**Figure S30D**). In sum, our AP-MS data are consistent with cholesterol-dependent remodeling of the EMC7 interactome.

Looking beyond the EMC complex, one additional intriguing feature of the competitive POCA dataset was the sizable number of proteins that showed cholesterol treatment-induced enhanced, rather than competed, protein capture by probe **1** (**Figure S25**, and **Figure S31A**). For the enhanced interactor subset, a number of plasma membrane proteins with clear links to cholesterol, such as NOTCH and LDLR stood out, together with proteins associated with lipid rafts such as SLC2A1 and ATP1B3^107^ . More broadly, both plasma and cell surface proteins were enriched in this subset, as revealed by both GO and subcellular compartment analysis (**Figure S31B, C**). As addition of exogenous cholesterol is known to induce the formation of cholesterol-rich nanodomains within the plasma membrane^108^, these findings are consistent with chol-POCA detection of proteins within these cholesterol-enriched nanodomains (“lipid rafts”).

### Cholesterol-POCA is compatible with HDL delivery and identifies shared and state-dependent interactors

Given the importance of HDL-cholesterol (HDL-C) transport to the liver for systematic cholesterol homeostasis and reverse cholesterol transport^109,110^, we next opted to assess whether POCA analysis could be performed using HDL-complexed probe delivery—when compared to mβCD complexation, HDL delivery offers the advantage of better modeling physiological cholesterol uptake. We drew inspiration from recent work^28^ that showed the utility of HDL cholesterol diazirine probe complexation for imaging-based applications—however, to our knowledge cholesterol probe-containing lipoprotein particles have not yet been extended to chemoproteomic applications. Therefore, following established methods^111^, we generated cholesterol-JF_570_ (probe **1**) complexed HDL particles (HDL-**1**, **Figure 4A**). Primary murine hepatocytes were then subjected to POCA analysis with HDL-**1** (100 µg/mL, corresponding to a maximum dose of 100 nM **1** assuming 100% of cholesterol in the HDL particle is replaced by **1**), which revealed light-dependent enrichment of 492 total proteins (**Figure 4B**). Illustrating the capacity of the HDL-POCA analysis to capture cholesterol interactors, a number of light-dependent enriched proteins stood out with known cholesterol-relevance, including the HDL-binding surface protein SR-B1 and the known SR-B1-interactor Na(+)/H(+) exchange regulatory cofactor NHE-RF2 (NHERF2)^112^, perilipin-2 (PLIN2), a structural component of LDs^113^, liver carboxylesterase 1 (CES1), which hydrolyzes cholesteryl esters in LDs and cholesterol-binding lysosomal associated membrane glycoprotein 1 (LAMP1)^74^.

**Figure 4.**
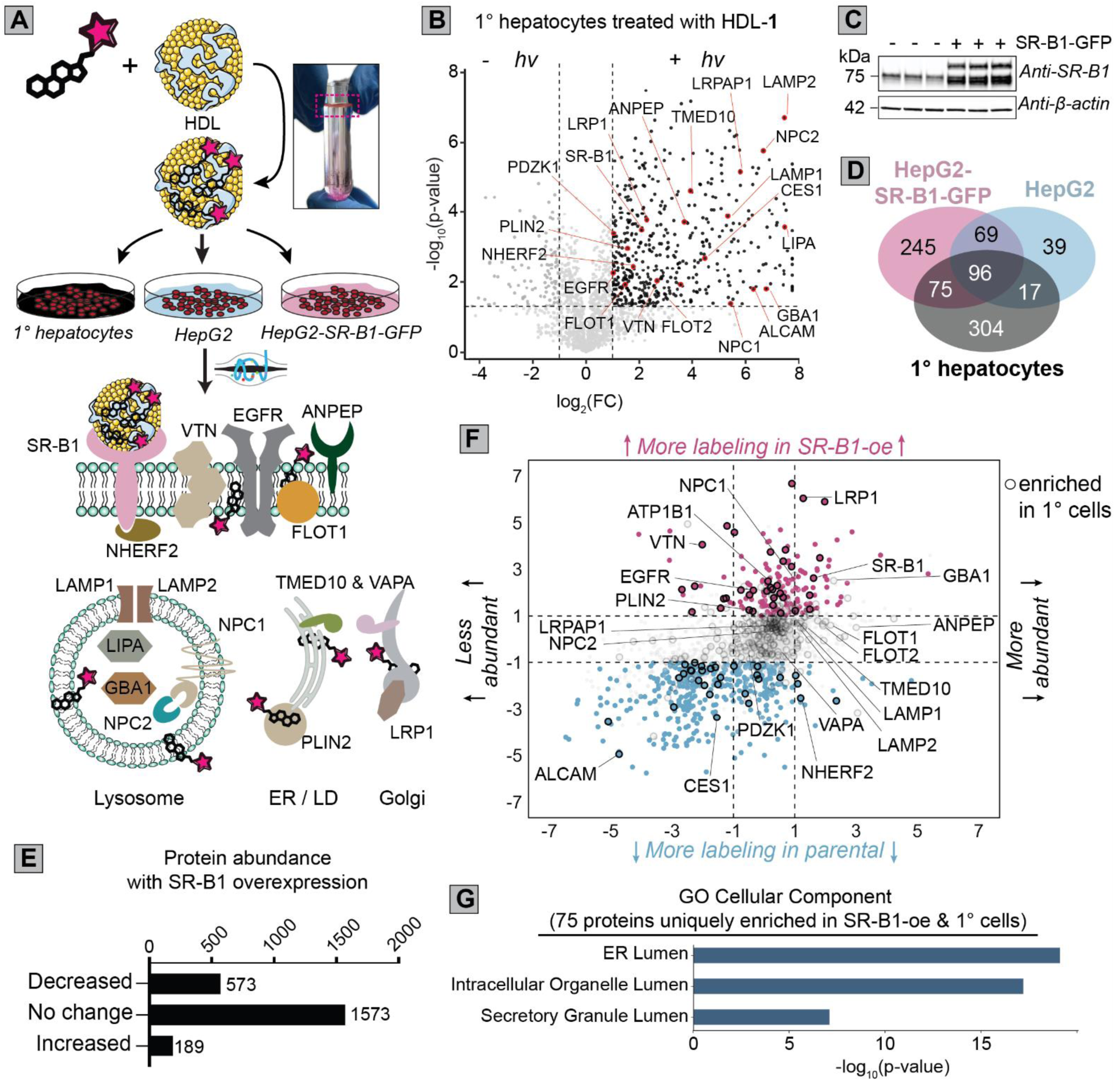
Proteomic profiling of cholesterol-associated proteins in hepatocytes using HDL-complexed POCA probe 1. **A**) Schematic workflow for the HDL complexation of probe 1 and enrichment of different cellular proteins in hepatocyte cell lines. **B**) Light-dependent POCA analysis of cholesterol depleted (simvastatin and mevalonate in LPDS) primary murine hepatocytes with HDL-complexed probe **1** (100 μg/mL)**. C)** Comparison of light-dependent HDL-enriched proteins from primary hepatocytes in ‘B’ to proteins identified for HDL-POCA labeling of HepG2 and SR-B1-overexpressing HepG2 after cholesterol depleted (Simvastatin and mevalonate in LPDS) and POCA labeling with HDL-complexed probe **1** (100 μg/mL). **D**) Immunoblot confirmation of SR-B1 overexpression in HepG2 cells. **E**) Subcellular compartments enriched for subset of proteins identified in primary hepatocytes and SR-B1-overexpressing HepG2 (75 proteins from ‘D).’ **F**) Assessment of how SR-B1 overexpression impacts cellular protein expression levels, as analyzed by LFQ analysis of bulk tryptic digests comparing HepG2 to SR-B1-overexpressing HepG2 cells. **G)** Assessing the contributions of SR-B1-induced changes to protein abundance to POCA enrichment. LFQ bulk abundance changes from ‘F’ were compared to SR-B1-induced changes to POCA enrichment from ‘D.’ Blue-decreased protein abundance, magenta-increased protein abundance. Circled points-proteins enriched by HDL-POCA in primary hepatocytes from ’B.’ Proteins labeled in parts ‘B’ and ‘G’ according to annotations with cholesterol processes. For proteins with log_2_(FC) >1 or <-1, only data points with p<0.05 are shown. For measures of statistical significance used in parts ‘B’, ‘C’, ‘E’, ‘F’, and ‘G’, variances were calculated for each sample-condition pairing and a corresponding two-tailed t-test was performed to generate p-values. n=3 per group for MS datasets. MS data can be found in **Data S4**.

Inspired by the capture of SR-B1, which is responsible for the vast majority (85%) of HDL-C ester uptake in hepatocytes^111^, we next tested whether we could combine genetic and chemoproteomic approaches to profile SR-B1-dependent cholesterol-interacting proteins. The specific steps and protein players involved in non-vesicular selective HDL-C uptake downstream of SR-B1 remain less well characterized than those involved in vesicular uptake^14^. We obtained HepG2 cells stably overexpressing SR-B1, with approximately three-fold increased expression compared to the parental cell line (**Figure 4C** and **Figure S32**). We then subjected both parental and SR-B1-overexpressing cells to POCA labeling with HDL-**1** after overnight cholesterol starvation (simvastatin and mevalonate in LPDS media) following the workflow shown in **Figure 4A**. Consistent with SR-B1 driving increased HDL particle uptake, SR-B1 overexpression was observed to afford a twofold increase in the number of proteins captured by HDL-POCA—485 significantly enriched for SR-B1-overexpressing cells versus 221 for parental HepG2 (**Figure 4D** and **Figure S33**). Further suggestive of POCA capture of proteins related to SR-B1-specific HDL-C uptake, 320 proteins (**Figure 4D** and **Figure S33**) were only enriched in the SR-B1 overexpressing HepG2 cells but not the parental cells, with 75 of these proteins also enriched in our hepatocyte dataset. This subset contained a number of SR-B1-associated proteins including SCARB2, CES1, GBA1, and low-density lipoprotein receptor-related protein 1 (LRP1)^114–116^ (**Data S4**). However, not all cholesterol-relevant proteins were enriched in the SR-B1 overexpressing cells—NPC1, FLOT1, flotillin-2 (FLOT2), SR-B1, LRPAP1, and ANPEP all showed universal enrichment in all three datasets (**Figure 4B**, **Figure S34**, **S35**, and **Data S1**), with previous literature showing that each of these proteins interact with cholesterol^27,52,84,117^.

We postulated that our SR-B1 overexpressing cell line would likely feature more widespread changes in protein abundance that could complicate delineation of bona fide SR-B1-induced changes to POCA labeling from altered bulk protein abundance. Therefore, we next opted to assess SR-B1-induced changes in protein expression, using LFQ proteomic analysis of bulk tryptic digests. Somewhat unexpectedly, we observe that SR-B1-overexpression caused decreased expression for a sizable majority (573/762) of the measured SR-B1-dependent protein abundance changes (**Figure 4E** and **Data S4)**. Many of the proteins with decreased expression were observed to be involved in cholesterol biosynthesis, including farnesyl pyrophosphate synthase (FDPS), hydroxymethylglutaryl-CoA synthase, cytoplasmic (HMGCS1), mevalonate kinase (MVK), phosphomevalonate kinase (PMVK), and sterol regulatory element-binding protein 1 (SREBF1). GO analysis (**Figure S36**, and **Figure S37**) further corroborated decreased expression of metabolic proteins as a general consequence of SR-B1 expression, consistent with decreased de novo cholesterol biosynthesis. 189 proteins were observed to show SR-B1-dependent increased expression (**Figure 4E**), including notable cholesterol-associated proteins such as SOAT1, oxysterol-binding protein-related protein 8 (OSBPL8), translocator protein (TSPO), hepatocyte nuclear factor 4-alpha (HNF4A), LAMTOR1^118^, FLOT1, FLOT2, and ANPEP^52^, which are implicated in cholesterol metabolism, cholesterol handling, or HDL binding.

With both bulk proteomic and POCA datasets in hand, we next turned to cross-dataset analysis, with the goal of delineating bona fide SR-B1-dependent interactions. All three datasets showed similar distributions in the subcellular annotations of enriched proteins (**Figure S38**). We found that 115 total proteins showed substantial SR-B1-mediated HDL-**1** interactions with no significant change in protein abundance (**Figure 4F**). Within the increased interactor subset, a number of proteins stood out as having established ties to cholesterol, including the sodium/potassium-transporting ATPase subunit beta-1 (ATP1B1), which binds HDL apoA-I ^119^, the cholesterol binding protein vitronectin (VTN)^120^, and the epidermal growth factor (EGFR) that is known to associated with lipid rafts^121^. More broadly, and illustrating the likely cholesterol-relevance of this subset, GO compartment analysis revealed strong enrichment for proteins localized to the ER lumen (**Figure 4G**), which is consistent with the ER’s established sensitivity to intracellular cholesterol levels^20^. Intriguingly, and hinting at probe **1** shuttling from the PM to the ER via the trans-Golgi network, we also noted that LRP1, a protein that is trafficked by the Golgi to the cell surface and localizes to lipid rafts^122^, showed markedly higher POCA capture relative to protein expression changes (**Figure 4F**).

### POCA-induced crosslinking captures Aster oligomerization

The Aster proteins, which are sterol transport proteins that move cholesterol from the PM to the ER^22–24,110,123^, were unexpectedly absent from all of our cholesterol POCA datasets. As we additionally did not identify any of the Aster proteins in our HepG2 bulk proteomic analyses (**Data S4**), we postulated that low protein abundance was hindering Aster detection. As the Aster proteins are known to undergo cholesterol-dependent changes to subcellular localization^24^ (**Figure S39A**) and homo-/hetero-oligomerization^23^ (**Figure S39B**), we also were particularly interested in using these proteins as a model system to test the capacity of our Halo-POCA platform to capture state-dependent changes to the Aster interactome that could additionally shed light on additional proteins involved in cholesterol transport. Inspired by these dual motivations, we generated HaloTag fusions of both the full-length protein (HaloTag-mAster-B) and a truncated construct lacking the C-terminal ER domain (ERD) required for ER anchoring and oligomerization^23^ (HaloTag-mAster-B-ΔERD, **Figure 5A**).^23^Gel-based analysis, using either the NBII-165, cholesterol-JF_570_ (**1**), or HaloTag ligand, revealed strong and specific labeling of the ΔERD construct for all three reagents (**Figure 5B**, **5C**). This finding supports both the utility of our HaloTag construct for protein-directed proximity labeling, and, more importantly, that the Aster proteins do bind to our cholesterol-JF_570_ probe. Unexpectedly, for the full-length Aster fusion we observed weaker labeling with cholesterol-JF_570_ and lower abundance of the fusion by immunoblot (**Figure 5B**). In the case of Halo-POCA using JF_570_-HTL we observed near complete disappearance of the protein detectable by immunoblot in a light and photosensitizer-dependent manner (**Figure 5C**).

**Figure 5:**
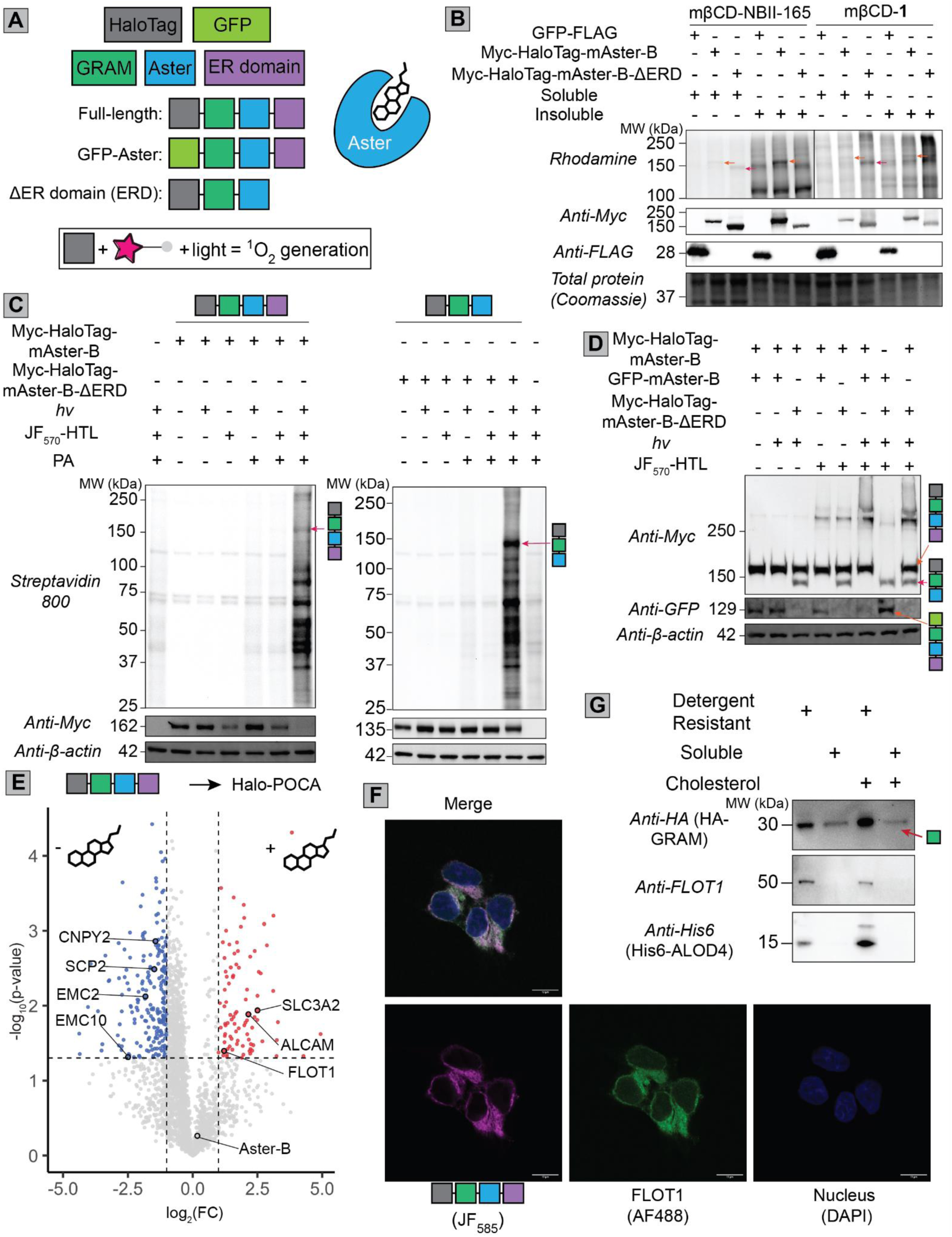
Investigation of the cholesterol transport protein Aster-B using Halo-POCA. **(A)** Shows scheme of murine (mAster) fusion proteins and highlights the cholesterol-binding Aster domain. (**B**) Cholesterol probes label full-length and ΔERD Aster constructs. HEK293T cells were transfected with the corresponding HaloTag-Aster construct, treated with mβCD-complexed probe (10 μM), labeled with the appropriate workflow, click conjugation to rhodamine-azide and visualization of the labeled proteins by fluorescent gel. (**C**) Gel-based analysis of photocatalytic protein alkynylation of the full-length and GRAM fusion proteins transiently overexpressed in HEK293T cells subjected to POCA labeling, click conjugation to biotin-azide and streptavidin blot. Arrows indicate labeling of the Aster construct. (**D**) Immunoblot detection of the indicated transiently co-expressed Aster fusion proteins after labeling with JF_570_-HTL and light irradiation in HEK293T cells. (**E**) POCA analysis of transiently overexpressed full-length HaloTag-mAster-B comparing cholesterol-depleted (1% LPDS/DMEM, overnight) versus cholesterol-loaded (addition of mβCD-cholesterol to depleted cells, 100 μM, 40 min) HEK293T cells. n=3 per group. (**F**) Imaging of HeLa cells stably expressing HaloTag-mAster-B, labeled with JF_570_-HTL, and stained for FLOT1. Additional images can be found in **Figure S44**. Scale bars = 10 μm. Each channel is a 1.0 μm thick slice. (**G**) Immunoblot assessment of protein localization to detergent resistant versus detergent soluble fractions derived from HEK293T cells overexpressing GRAM-Aster-B (for full-length protein see **Figure S46**), pulsed with either vehicle of mβCD-cholesterol (100 μM), stained with accessible cholesterol binder ALOD4, and fractionated using a linear iodixanol gradient. For measures of statistical significance used in part ‘E’, variances were calculated for each sample-condition pairing and a corresponding two-tailed t-test was performed to generate p-values. n=3 per group for MS datasets. MS data can be found in **Data S5**.

Protein crosslinking between histidine and other amino acids is a well-established consequence of ^1^O_2_ generation^124^, with the crosslinks occurring between oxidized histidine and histidine^125^, lysine^126^, and cysteine^127^ residues. Thus, we hypothesized that the POCA-induced depletion of HaloTag-mAster-B could be a consequence of such ^1^O_2_-induced protein crosslinking (**Figure S40A**), with more pronounced crosslinking expected for the full-length oligomerization competent protein and attenuated crosslinking for the ΔERD construct. To test this hypothesis, we subjected HEK293T cells co-expressing different combinations of Aster fusion proteins, including the aforementioned HaloTag-mAster-B, and HaloTag-mAster-B-ΔERD together with GFP-mAster-B (**Figure 5A**), to irradiation in the presence of JF_570_-HTL. We expected that intermolecular crosslinking between the GFP and HaloTag fusion proteins could be detected from the disappearance of monomeric protein signal together with the appearance of larger, slower migrating crosslinked protein bands. Consistent with intermolecular crosslinking, we observed JF_570_-HTL-dependent formation of high molecular weight Aster-B protein, with the expected ∼320 kDa size of crosslinked Aster fusion proteins, together with almost complete disappearance of GFP signal at the expected molecular weight of the monomer protein (**Figure 5D**). The HaloTag-mAster-B-ΔERD construct showed no appreciable crosslinking with GFP-mAster-B (**Figure 5D**). HaloTag-[ΔGRAM]-mAster-B, which retains the ER and Aster domains but not the cholesterol-binding GRAM domain, showed similar crosslinking activity to HaloTag-mAster-B (**Figure S40B**), further corroborating that the ERD is required for crosslinking. No appreciable crosslinking was observed when an inefficient photosensitizer, JF_585 61,_ was used (**Figure S40B**). These results are consistent with a model whereby domain-dependent Aster-oligomerization and localized ^1^O_2_ production are required for efficient crosslinking.

### Protein-directed POCA captures the state-dependent Aster interactome

Although initial unexpected our observation of Aster crosslinking corroborated the likelihood that the Aster HaloTag fusion protein was forming oligomers, consistent with the endogenous protein^23^. Thus, we opted to next subject the Aster-B HaloTag fusion protein to POCA analysis using the JF_570_-HTL photosensitizer. Increased cellular cholesterol levels are known to induce Aster-B translocation to ER-PM contact sites^23,24^ (**Figure S39A**). Therefore, we hypothesized that the Aster interactome should change depending on cellular cholesterol content. To test this hypothesis, we compared Aster-B interactors captured by Halo-POCA for cholesterol loaded versus cholesterol starved HEK293T cells (**Figure 5E** and **Figure S41A**), both for full-length HaloTag-mAster-B and the ΔERD constructs. Gel-based analysis of photocatalytic protein alkynylation (**Figure S41B**), with reduced irradiation intensity to minimize crosslinking, revealed construct-specific banding patterns, consistent with differences in the protein interactomes between the full-length and ΔERD constructs. Despite the ΔERD construct showing higher overall expression, similar overall coverage was observed for both constructs (**Figure S42**). Pleasingly, and consistent with full-length Aster localization to the ER, we observed a substantial increase in capture of ER proteins for full-length Aster-B 40/143 (28%) when compared to the 7/352 (2%) proteins enriched by the ΔERD construct (**Figure S43**). Exemplary ER proteins enriched include the lipid transfer protein sterol carrier protein 2 (SCP2), which functions to move cholesterol and other lipids between the ER and PM^128^, EMC2, EMC10, and CNPY3, which had all also been captured in our chol-POCA analysis (**Figure 3B**, **Data S1** and **Data S3**). Consistent with the movement of Aster-B to the PM when cells are loaded with cholesterol, we observed capture of PM proteins only in the cholesterol-loaded condition for the ΔERD construct (**Figure S44**). Looking at the full-length Aster construct, a number of PM-annotated proteins including the chol-POCA identified SLC3A2, ALCAM, and FLOT1 were enriched in the cholesterol-loaded condition. Taken together, the cholesterol-dependent shift in ER-PM localization of Aster-enriched protein provides evidence in support of POCA’s capacity to capture state-dependent interactions.

The established association between FLOT1 and lipid rafts^129^ prompted us to next ask whether Aster-B was also associated with lipid rafts. Immunofluorescence analysis revealed colocalization of Aster-B and FLOT1, which indicated that Aster-B associates with detergent-resistant membranes (DRMs), (**Figure 5F** and **Figure S45**). Furthermore, we found that both GRAM-Aster-B and Aster-B co-fractionated with FLOT1 in the detergent-resistant membrane fraction. Prior to fractionation, the cells were incubated with ALOD4, a probe for the “accessible” cholesterol pool in the plasma membrane^27^. ALOD4 was specifically enriched in the detergent resistant fraction of cells and its abundance increased with cholesterol loading. We observed a marked increase in detectable GRAM-Aster-B protein in the ALOD4- and FLOT1-positive detergent resistant fraction after cholesterol loading (**Figure 5G** and **S46**). The expected transient nature of the GRAM-PM interactions highlights the capacity of POCA to capture dynamic and potential fleeting interactions upon changes to cell state.

## Discussion

The recent emergence of a number of photocatalytic-proximity labeling proteomic platforms^37–44,130^ highlights the strong demand for such light-dependent methods that robustly report the spatial environment and interactors of proteins as well as other biomolecules. Here we establish the POCA photosensitizer-based proximity labeling platform that offers the unique advantage of generalized compatibility, using the same labeling chemistry for both intracellular small molecule and protein-interactome analyses. To build POCA, we first identified a suitable cell penetrating photosensitizer, JF_570_, which we vetted for proximity labeling applications using the HaloTag system. We find that POCA analysis using the nucleoporin NUP153 as bait captures nearly all the subunits of the nuclear pore complex. Stepping beyond these benchmarking analyses, we then demonstrate the capacity of POCA to capture the cholesterol interactome, using both a control palmitate probe as well as cholesterol diazirine probes to confirm performance and delineate specific interactors. POCA analysis using HDL delivery revealed cholesterol-induced stabilization of subunits of the EMC insertase, which functions cooperatively with SEC61 to deliver newly synthesized proteins to the ER membrane. When applied to the Aster proteins, protein-directed POCA faithfully delineated state-dependent protein oligomerization, as captured by photosensitizer-induced crosslinking, and uncovered heretofore unreported colocalization of the Aster proteins with flotillin-rich detergent resistant membranes. Thus, the POCA platform is a versatile system capable of capturing transient and stable interactions spanning subcellular compartments and types of biomolecules.

Towards discovery of protein modulators of cholesterol movement, our work revealed that the comparatively large JF_570_ modification did not abolish interactions with known cholesterol binders, including OSBP, NPC1, NPC2, LAMP1, LAMP2, OSBPL9, and Aster-B. This finding is perhaps not surprising given the widespread use of the comparatively bulky fluorescent 22-NBD-cholesterol for monitoring sterol binding and movement within cells^131^. When compared to less lipophilic molecules, cholesterol offered unique properties for interactomics. In our competition studies, we observed a sizable subset of proteins that showed increased, rather than off-competed, capture by chol-POCA in the presence of excess cholesterol. Amino acid transporter heavy chain SLC3A2 (SLC3A2/4F2) is one such protein that shows cholesterol-dependent changes to activity, forming a heterodimer with the L-Type Amino Acid Transporter 1 (LAT1) that is then stabilized through interaction with membrane cholesterol, modulating its transport activity^132^. EGFR, which showed interaction with HDL-**1** in an SR-B1-dependent manner, shows cholesterol-dependent changes to activity including resistance to inhibitors upon lipid raft localization^133^. More broadly, as many of these cholesterol-enhanced interacting proteins were found to be localized to the plasma membrane, we expect that our dataset will likely reveal new protein content associated with cholesterol-rich membrane microdomains, including key regulators of cell signaling processes.

Looking more broadly, our work also highlights the importance of delivery modality to the chemical probe under investigation. Our use of primary and immortalized hepatocytes, along with physiologically relevant HDL uptake of the probe molecule, revealed key differences in protein content labeled compared to mβCD delivery. Exemplifying these differences, while OSBP was robustly enriched by mβCD-**1**, it was not enriched in hepatocytes by HDL-**1**. In contrast, the ER-resident member TMED10 and the Golgi-resident member VAPA of the cholesterol-transporting super complex showed preferential enrichment by HDL-**1** over mβCD-**1**. While we cannot rule out differences in probe concentration between the two delivery modalities as causative for these different enrichment profiles, we expect that proteins captured by HDL-**1** more faithfully reflect physiological cholesterol transport.

Beyond the current implementation of POCA, we envision several areas for continued growth. Singlet oxygen has a comparatively large radius of labeling, which is a double-edged sword for proximity labeling, enabling capture of large protein complexes but also coming with the potential liability of non-specific bystander protein enrichment. We harnessed off-competition studies together with orthogonal genetic approaches to help delineate high and low confidence interactors. We envision that dose-dependent labeling analysis together with incorporation of additional trapping reagents that span varying reactivities into the POCA workflow, such as those used by the recently reported MultiMap method^42^ will likely enhance the resolution of the POCA platform to better define local protein neighborhoods. Complementary to advances in the trapping agents, the availability of photosensitizers spanning a range of wavelengths also offer untapped potential for sequential labeling studies using several orthogonal labeling reagents. Lastly, as our work highlights, ^1^O_2_-mediated protein crosslinking is likely a generalized feature of photocatalytic labeling platforms. Inspired by the oligomer-dependent crosslinking shown here for Aster-B, we anticipate widespread applications of the POCA methodology to more generally delineate state-dependent protein multimerization.

## Acknowledgements

We thank Luke D. Lavis and the Lavis Lab (Janelia Research Campus, HHMI) for providing the JF_570_- and JF_585_-HaloTag ligands. This study was supported by DP2 GM146246-02 (K.M.B.), Packard Fellowship (K.M.B.), 1P01HL146358 (PT, K.M.B), Leducq Transatlantic Network of Excellence 19CVD04 (P.T.), and the CNSI 2021 Noble Family Innovation Fund Seed Project Award (K.M.B, P.T). J.P. K. was supported by an American Heart Association postdoctoral fellowship (903306). A.R.J. was supported by NIGMS UCLA Chemistry Biology Interface T32GM136614. R.T.N was supported by NIGMS T32 GM008042 and an Audree Fowler Protein Science Fellowship from the UCLA Molecular Biology Institute. S.G.H was supported as a Jim Easton CDF Investigator. We thank the S10 program of the NIH Office of Research Infrastructure Programs (ORIP) grant S10OD028644 for NMR facilities funding. We thank Kelsey Martin for providing LSM880 microscope access and Sylvia Neumann for her assistance with microscopy and helpful discussions. We thank all members of the Backus and Tontonoz labs for their helpful suggestions.

## Author Contributions

A.B and K.M.B conceived of the study. A.B., JP.K., J.J.M., P.T., K.M.B contributed to the design and implementation of experiments. A.B., E.B., JP.K., A.R.J., M. V., and S.G.H. generated and analyzed data. A.B., E.B., N.B., T.F., D.T. performed chemical synthesis. A.B., JP.K., R.T.N., L.C., X.X. generated reagents and contributed to experiments. A.B., S.G.H, and X.X. contributed to visualization. A.B., E.B., and K.M.B. wrote the manuscript with assistance from all authors.

## Ethics declarations

The authors declare no known conflicts of interest.

## Supplementary information

Detailed methods of chemical synthesis, spectral characterization, and chemoproteomic sample preparation; Supplementary Figures S1-S46, Schemes S1-S5, Table S1-S5, (PDF)

Data S1: Proteomic data related to Figure 1; combined table of fold changes and p-values for all detected proteins (xlsx)

Data S2: Proteomic data related to Figure 2 (xlsx)

Data S3: Proteomic data related to Figure 3 (xlsx) Data S4 Proteomic data related to Figure 4 (xlsx) Data S5: Proteomic data related to Figure 5 (xlsx)

Video S1: Video of Cholesterol-JF_570_ uptake related to Figure 2G and Figure S24 (avi)

## Data Availability

The MS data have been deposited to the ProteomeXchange Consortium (http://proteomecentral.proteomexchange.org) via the PRIDE^134^ partner repository with the dataset identifier PXD054875.

## Code Availability

The R script used to calculate fold changes, perform two-tailed t-tests to generate p-values, compile the data, and generate plots is provided in Supplementary Data File S1.

