## Supplementary Information for "Lipid- and protein-directed photosensitizer proximity labeling captures the cholesterol interactome"

**Table of Contents**

1. Supplemental Schemes (pp. 3-6)
2. Supplemental Figures (pp. 7-49)
3. Methods (pp. 50-70)
4. Supplemental Tables (pp. 70-77)
5. NMR spectra (pp. 78-90)
6. References (pp 91-94)

### 1. Supplemental Schemes

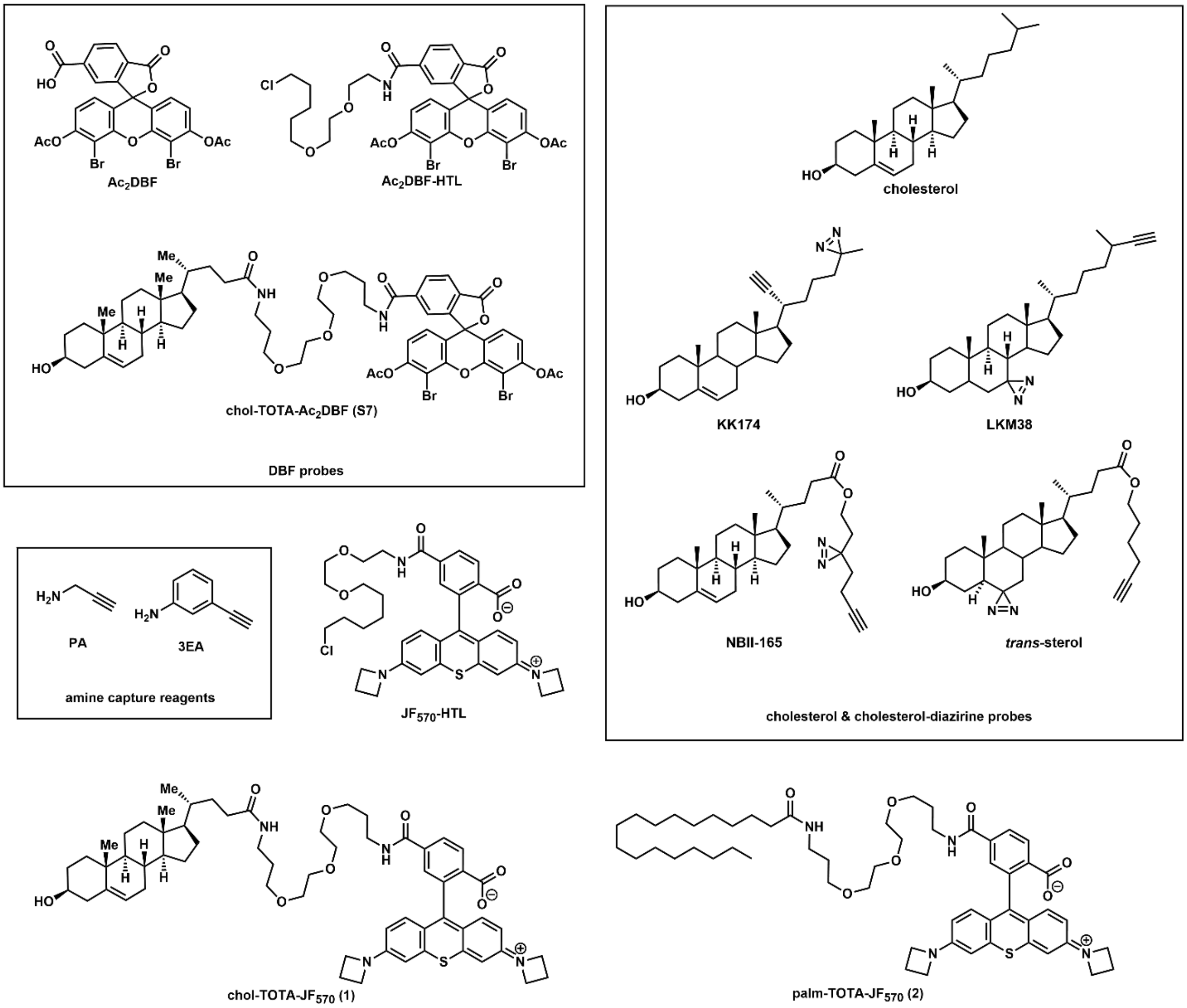


**Scheme S1:** Structures of key previously reported[1–4] and newly synthesized compounds employed in this study.

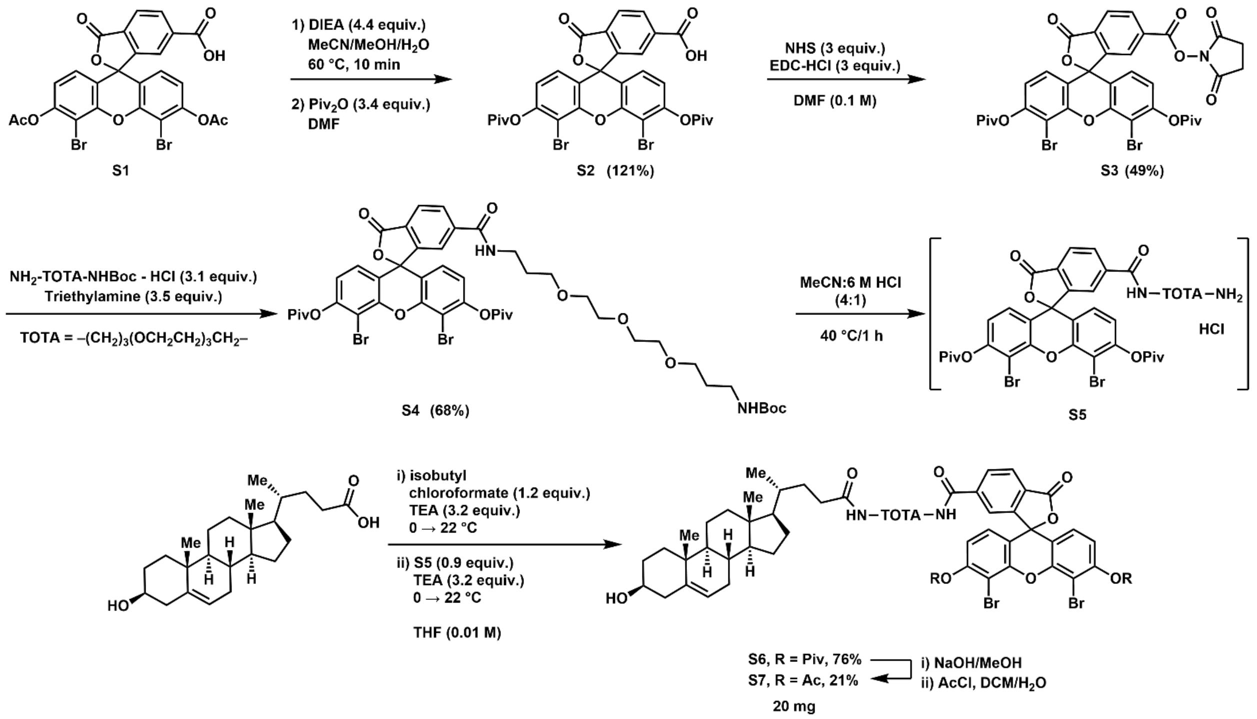


**<u>Scheme S2</u>: Synthesis of chol-TOTA-$Ac_2$DBF (S7).** Synthesis of the tail-linked cholesterol probe chol-TOTA-Ac2DBF was achieved in five steps starting from **S1**[Ref Spitale]. First, the phenol acetyl protecting groups were swapped to pivalate groups. The NHS ester was formed by treating **S2** with *N*-hydroxy succinimide and 1-Ethyl-3-(3-dimethylaminopropyl)carbodiimide hydrochloride (EDC-HCl) in DMF. Amide bond formation was achieved by addition of free base $NH_2$-TOTA-NHBoc (generated *ex situ* by combining triethylamine, TEA, and the amine hydrochloride salt) to furnish **S4**. Subsequent removal of the Boc group with HCl in mixed MeCN/$H_2O$ afforded the amine hydrochloride **S5**. The mixed anhydride of cholenic acid was formed by treatment with isobutyl chloroformate and TEA in anhydrous THF, and the amide bond was formed by addition of free base **S5** (generated *ex situ* by combining TEA and **S5**) to give **S6**. The pivaloyl protecting groups were removed by basic methanolysis, and the acetyl groups were installed using acetyl chloride (AcCl) in biphasic DCM/$H_2O$ in the presence of base to afford **S7** (chol-TOTA-$Ac_2$DBF).

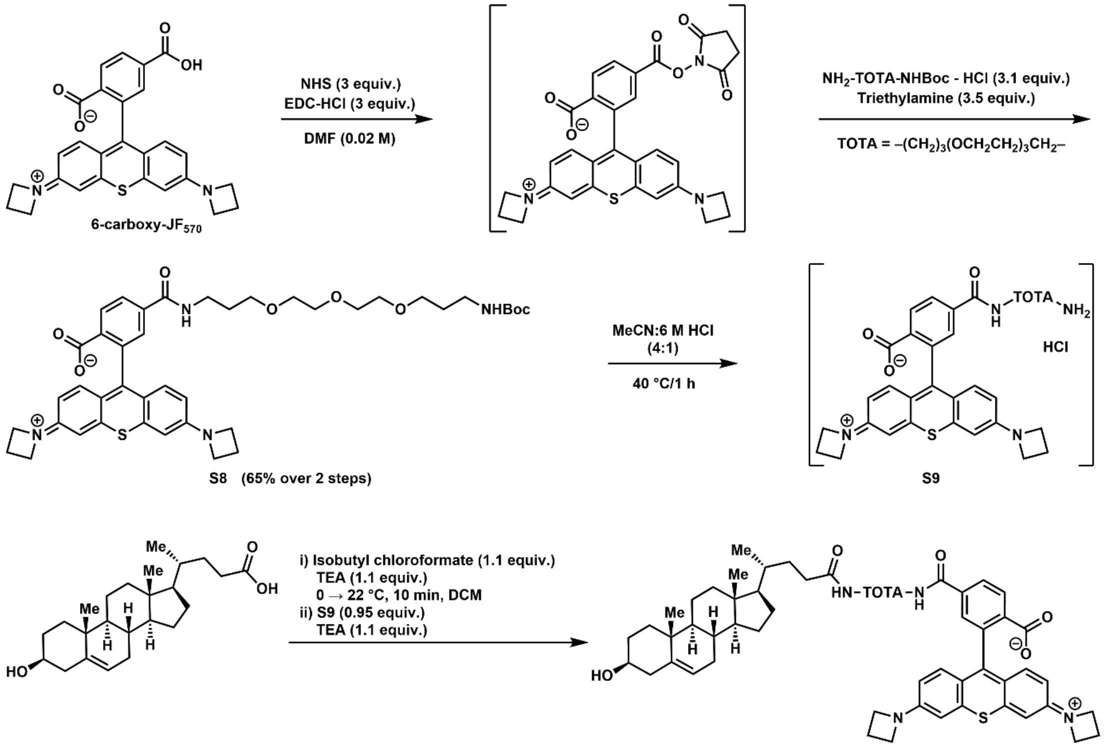


1 (18 mg, 33%)

<u>Scheme S3</u>**: Synthesis of chol-TOTA-JF$_{570}$ (1).** Synthesis of the tail-linked cholesterol probe **1** was achieved in four steps starting from **6-carboxy-JF$_{570}$**[4]. The NHS ester was formed by treating **6-carboxy-JF$_{570}$** with *N*-hydroxy succinimide and EDC-HCl in DMF. Amide bond formation was achieved by addition of free base $NH_2$-TOTA-NHBoc (generated *ex situ* by combining TEA and the amine hydrochloride salt) to furnish **S8**. Subsequent removal of the Boc group with HCl in mixed MeCN/$H_2O$ afforded the amine hydrochloride. The mixed anhydride of cholenic acid was formed by treatment with isobutyl chloroformate and TEA in anhydrous THF, and the amide bond was formed by addition of free base **S9** (generated *ex situ* by combining TEA and **S9**) to give **1** (chol-TOTA-JF$_{570}$) as a bright pink-purple solid.

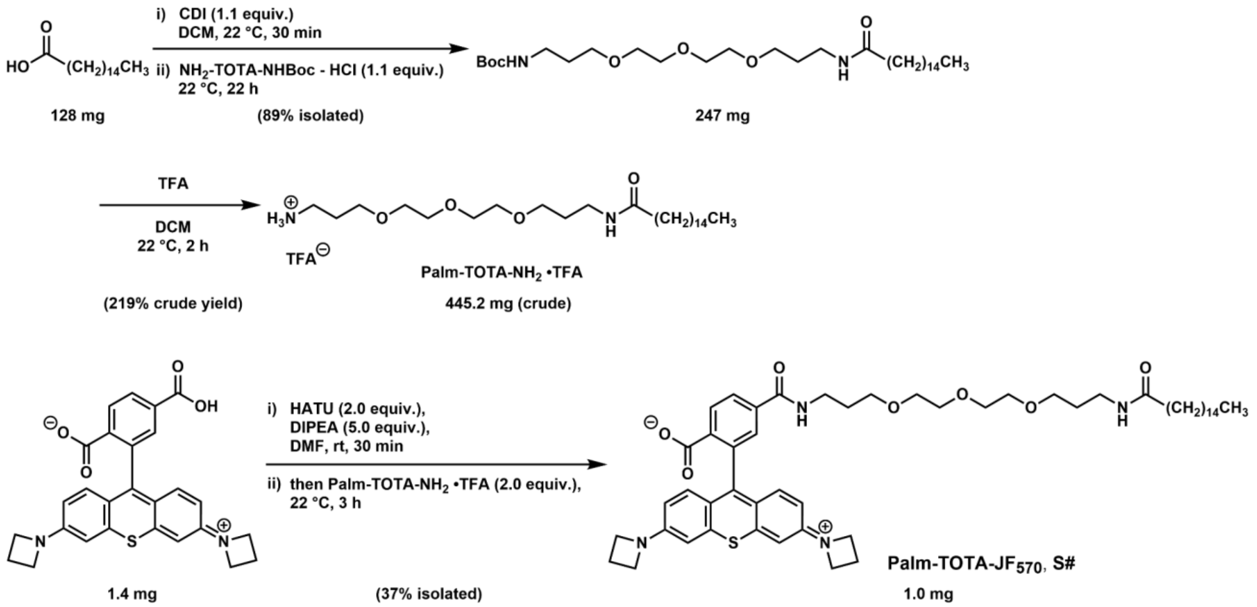


**Scheme S4: Synthesis of Palm-TOTA-JF$_{570}$ (2).** Synthesis of the palmitate-TOTA-JF$_{570}$ probe **2** was achieved in three steps starting from palmitic acid and **6-carboxy-JF$_{570}$**. The Boc-protected TOTA amide of palmitic acid was formed by combining palmitic acid and 1,1'-carbonyldiimidazole in DCM, then adding free base $NH_2$-TOTA-NHBoc (generated *ex situ* by combining TEA and the amine hydrochloride salt). Boc-deprotection using TFA in DCM furnished the crude TFA salt, which was used in the following step without purification. Amide coupling of Palm-TOTA-$NH_2$ with 6-carboxy-JF$_{570}$ was achieved using HATU coupling conditions in DMF with DIPEA as the base to afford **2** (1.0 mg, 37%) as a deep purple solid.

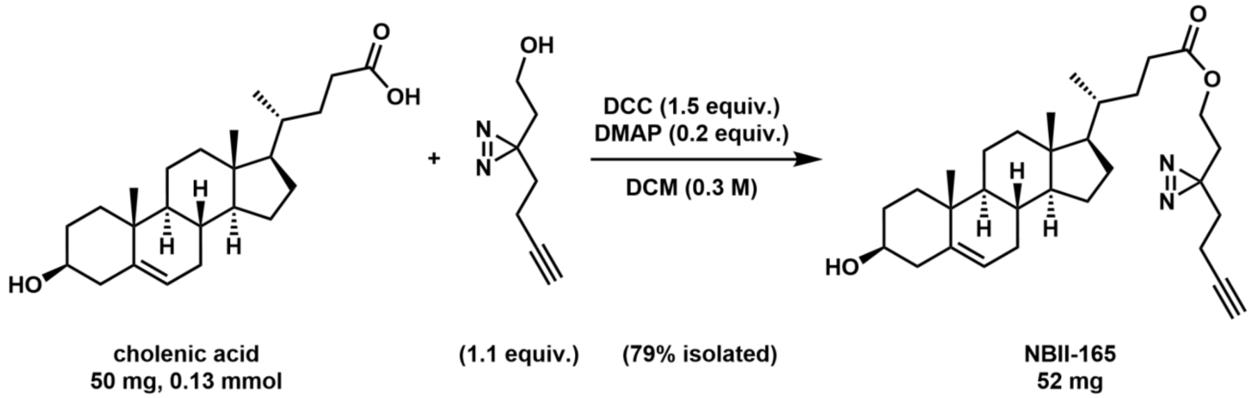


**Scheme S5: Synthesis of NBII-165.** Synthesis of the tail-linked cholesterol diazirine probe NBII-165 was achieved in one step from the previously-reported[5] alkynyl diazirine alcohol and cholenic acid. Ester formation was achieved using DCC and catalytic DMAP to furnish NBII-165 in 79% yield.

### 2. Supplemental Figures

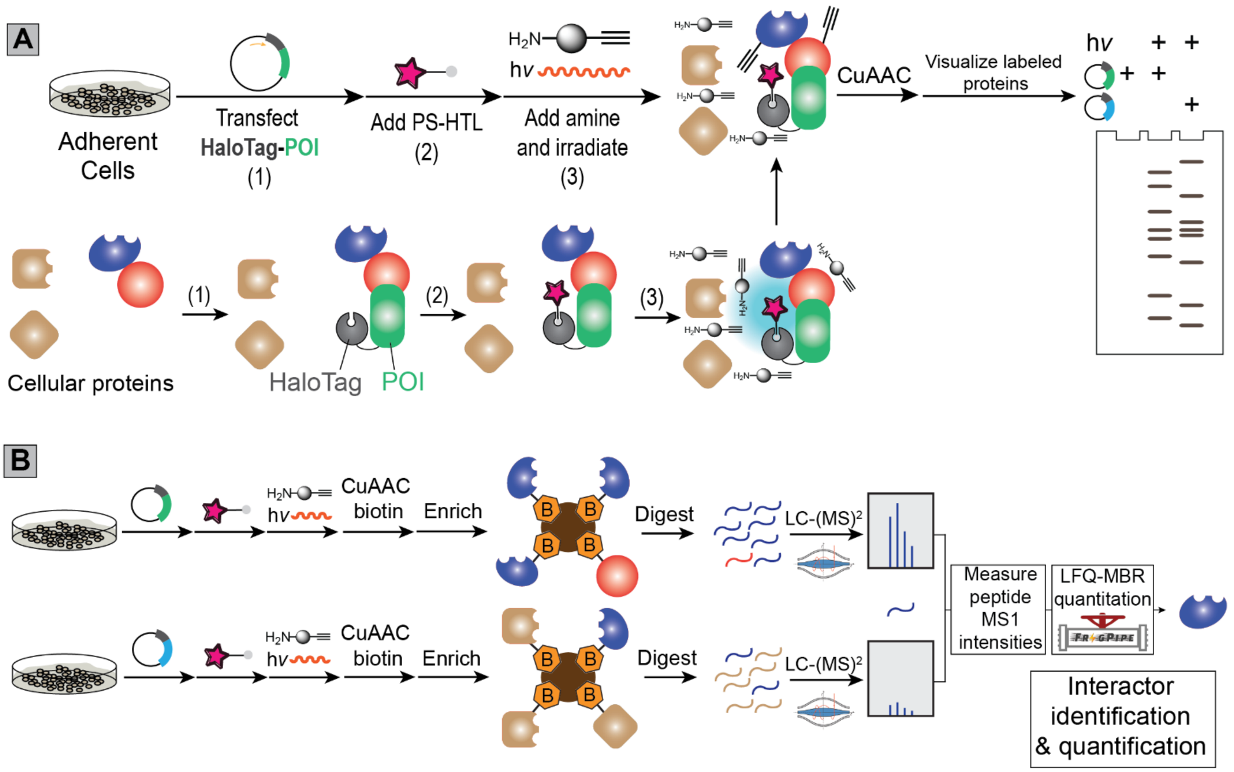


**<u>Figure S1</u>: Detailed workflows for gel-based and mass spectrometry-based protein labeling and identification via Halo-POCA proximity labeling.** For **(A)** gel-based and **(B)** mass spectrometry-based Halo-POCA, cells heterologously overexpressing a HaloTag fusion protein of interest (POI) are first incubated with a photosensitizer (PS)-HaloTag ligand (HTL). After a brief washout period, an alkyne-containing amine capture reagent, such as propargylamine (PA), is added, and the cells are irradiated with suitable wavelength light to produce singlet oxygen ($^1O_2$). The samples are then subjected to copper-catalyzed azide-alkyne cycloaddition (CuAAC) with an azido-fluorophore (e.g. tetramethylrhodamine-azide). For gel-based analysis, as shown in 'A,' labeling is then visualized by SDS-PAGE and in-gel fluorescence analysis. For proteomic analysis, as shown in 'B,' biotinylated proteins are subsequently enriched on streptavidin resin, subjected to on-resin tryptic digest, and LC-MS/MS analysis, followed by search and label free quantification (LFQ) [6] using MSFragger[6–9] (https://fragpipe.nesvilab.org/).

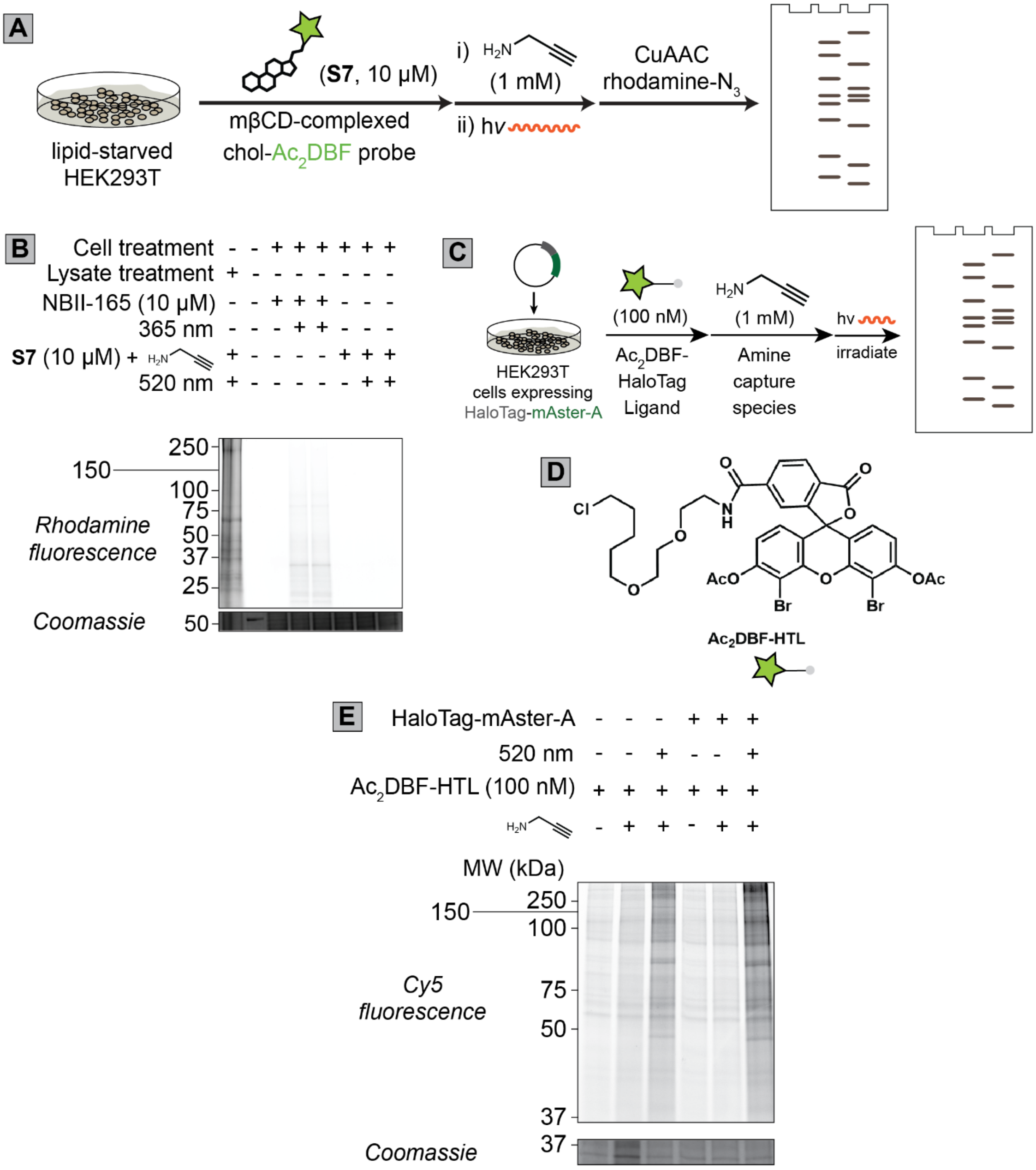


**Figure S2**: **Assessing the suitability of acetylated dibromofluorescein ($Ac_2DBF$)-functionalized cholesterol and chloroalkane reagents for cell-based POCA. (A)** Workflow used to test cell-based protein labeling for cells exposed to cholesterol-TOTA-$Ac_2DBF$ (**S7**) for chol-POCA labeling in HEK293T cells. **(B)** In-gel fluorescence comparison of HEK 293T cells or lysates subjected to labeling with either the positive control mβCD-complexed diazirine probe NBII-165 (10 μM, 10 min 365 nm irradiation for cell-based labeling) or mβCD-complexed **S7** (cholesterol-TOTA-$Ac_2DBF$, 1 μM for lysate based labeling or 10 μM probe concentration for cell-based labeling, with 5 min green LED irradiation) **(C)** Workflow followed to evaluate cell-based Halo-POCA protein labeling with $Ac_2DBF$-HaloTag Ligand ($Ac_2DBF$-HTL) and HaloTag-murine-

Aster-A (mAster-A). **(D)** The structure of $Ac_2$DBF-HTL is shown on the right. **(E)** In-gel fluorescence comparison of relative protein labeling for HEK293T cells labeled with $Ac_2$DBF-HTL (100 nM, 10 min) and subjected to the Halo-POCA workflow as shown in 'B'. For B and D, samples were subjected to CuAAC with sulfo-Cy5-azide and visualized by in-gel fluorescence after SDS-PAGE separation.

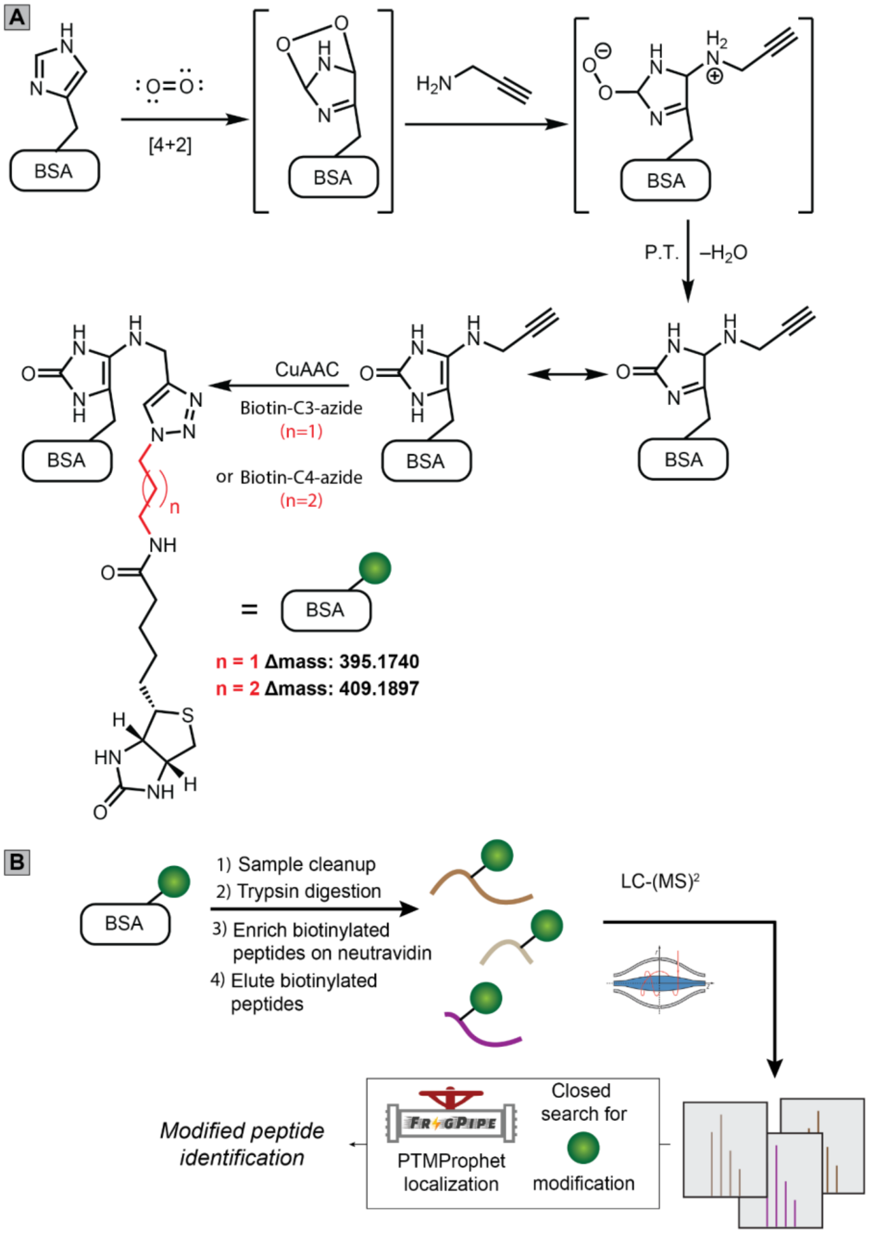


**Figure S3**: **Site of labeling analysis for bovine serum albumin (BSA) subjected to POCA labeling with either C3- or C4-biotin confirms the expected dihydroimidazolone modification. (A)** The expected mechanism[10,11] of histidine labeling by propargylamine for BSA subjected to 6-carboxy JF$_{570}$ (10 µM) and propargylamine (1 mM) followed by click conjugation to our previously reported C$_3$-biotin-azide and C$_4$-biotin-azide capture reagents[12], which were selected for this analysis to generate a +14.0157 Da mass difference between C$_3$- and C$_4$-labeled peptides—this mass shift provides additional corroborating evidence in support of the modified nature of the peptide spectrum matches (PSMs). **(B)** Biotinylated BSA was subsequently subjected to our established[13] SP3[14]-based peptide enrichment workflow followed by LC-MS/MS analysis and FragPipe search to identify biotin-modified peptides harboring the expected modifications on histidine residues, +395.1740 or +409.1897 for $C_3$-and $C_4$-biotin-azide, respectively.

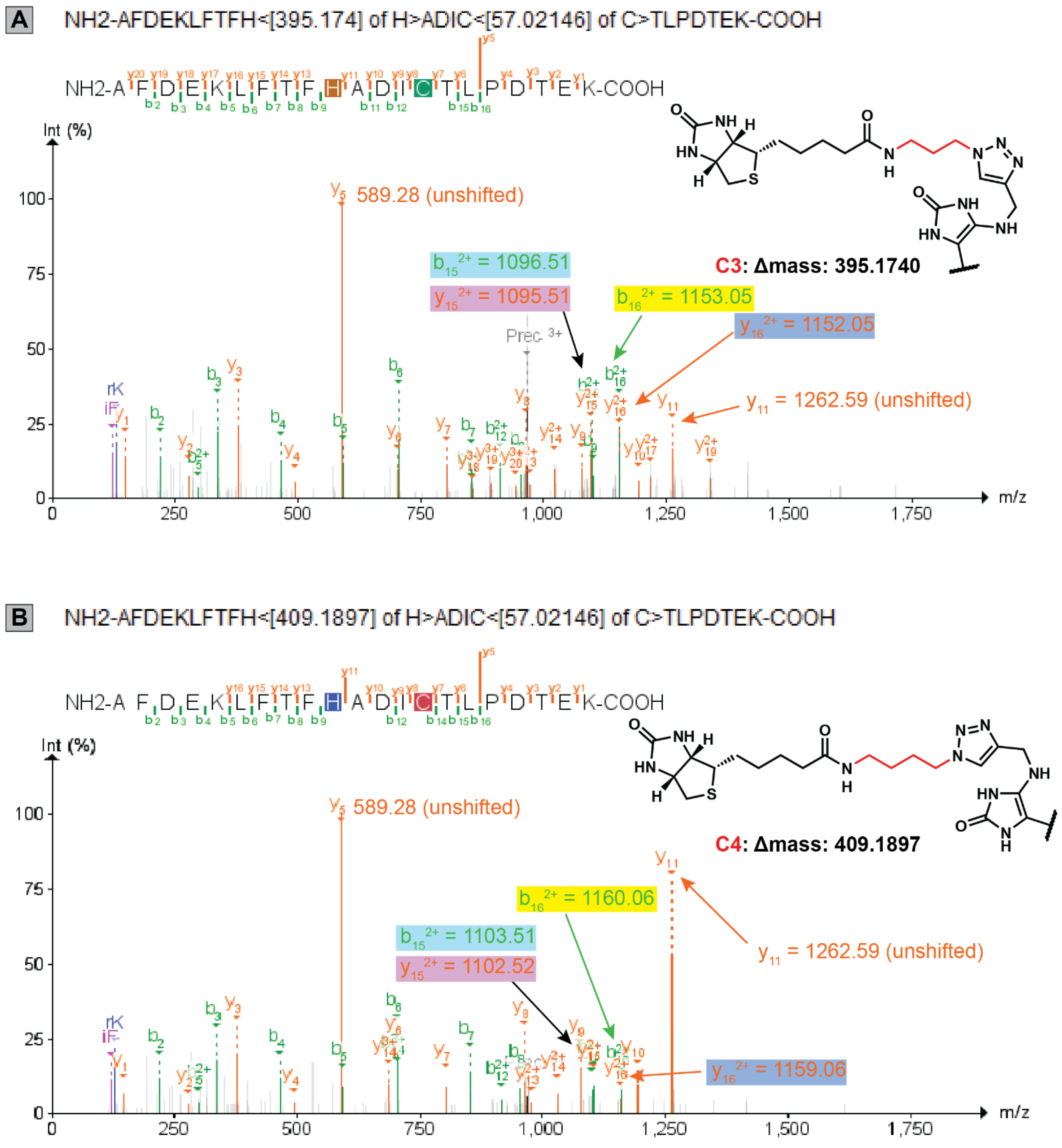


**<u>Figure S4</u>: MS/MS spectra for identified POCA modified peptides from BSA corroborates histidine modification by propargylamine.** Shown are representative MS/MS spectra for the same peptide sequence from BSA, which was assigned as biotinylated on His533 in both our $C_3$- and $C_4$-biotin datasets, generated as described in **Figure S3**. Cysteine capping with iodoacetamide afforded carbamidomethylation (+57.02146 Da) on Cys536. The modification localization sites were assigned using PTMProphet[15], and annotated spectra were generated by Fragpipe PDV viewer[16]. MS data can be found in **Data S1**.

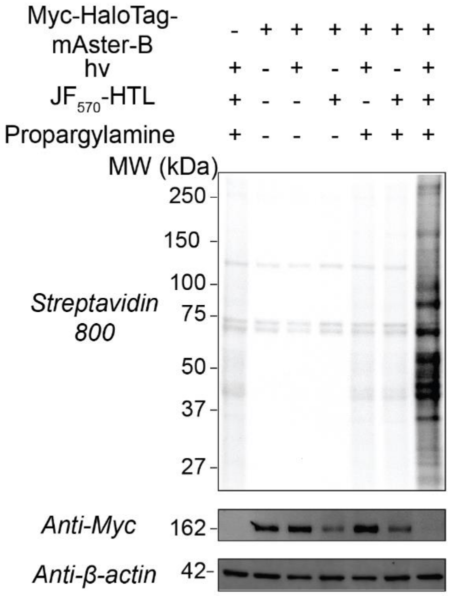


**Figure S5**. **$JF_{570}$ is a suitable photosensitizer for cell-based POCA**. HEK293T cells with or without stably overexpressed HaloTag-mAsterB were subjected to the Halo-POCA protocol. Following the workflow shown in Figure S1A, cells were first dosed with $JF_{570}$-HaloTag Ligand ($JF_{570}$-HTL, 100 nM, 10 min) followed by washout to remove excess reagent (2 x 20 min), addition of PA (10 mM) and irradiation (170,000 Lux, 5 min,15 W yellow LED). After cell lysis, CuAAC with biotin azide, and SDS-PAGE, the protein labeling was visualized by streptavidin blot. No treatment (-) conditions were processed identically with vehicle or no-light treatment replacing the indicated labeling step.

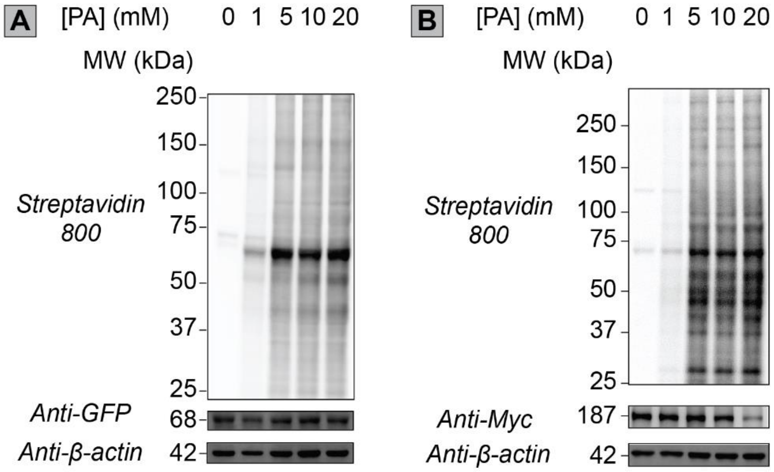

**<u>Figure S6</u>**: **Relative protein labeling by Halo-POCA is dependent on the concentration of the amine capture reagent**. Following the workflow shown in Figure 1A, HEK239T cells transiently expressing either **(A)** HaloTag-eGFP-mito (pERB254[17]) or **(B)** Myc-HaloTag-NUP153 were treated with $JF_{570}$-HTL (100 nM, 10 min) followed by media washouts, addition of PA (0–20 mM) and irradiation (15 W yellow LED, 170,000 Lux max intensity, 5 min, on ice), lysis, click conjugation to biotin-azide, SDS-PAGE, and streptavidin blot.

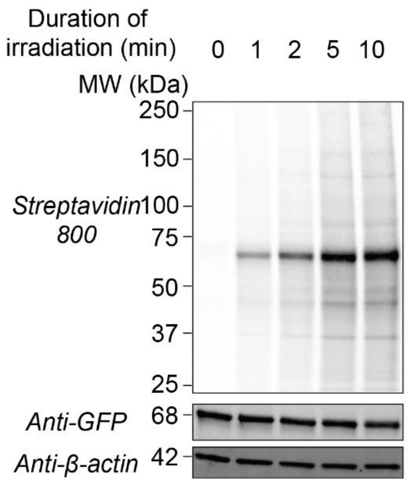


**<u>Figure S7</u>**: **Relative POCA labeling is dependent on the duration of irradiation**. Following the workflow shown in Figure S1A, HEK239T cells transiently expressing HaloTag-eGFP-mito (pERB254[17]) were subjected to gel-based Halo-POCA analysis. Cells were treated with $JF_{570}$-HTL (100 nM, 10 min) followed by media washouts, followed by PA (10 mM) and irradiation (15 W yellow LED, 170,000 Lux max intensity, 0–10 min, on ice), lysis, click conjugation to biotin-azide, SDS-PAGE, and Streptavidin blot.

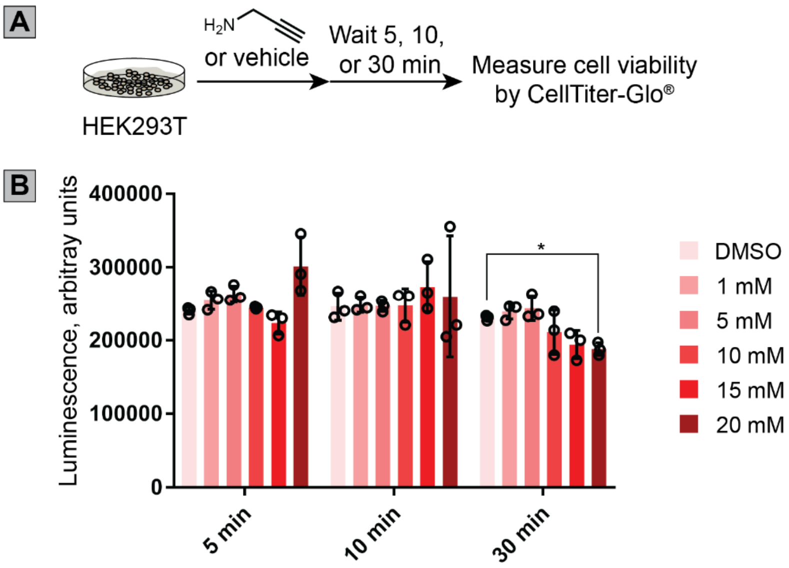

**<u>Figure S8</u>**: **Short exposure to propargylamine (PA) does not significantly affect the viability of HEK293T cells**. **(A)** Workflow to assess cell viability of HEK293T cells. **(B)** Graph of cell viability data, measured as a function of time and PA concentration relative to DMSO control. Cells in a 96-well plate-based were incubated with PA for the indicated treatment times and concentrations with 0.4% (v/v) DMSO. Shown are the mean luminescence as height and standard deviation as error bars. Each condition was performed in triplicate. Statistical significance was assessed by student's t-tests comparing PA treated samples to DMSO. Where indicated: * $p$ <0.05*.* All other comparisons to the DMSO control were not significant ($p$ > 0.05).

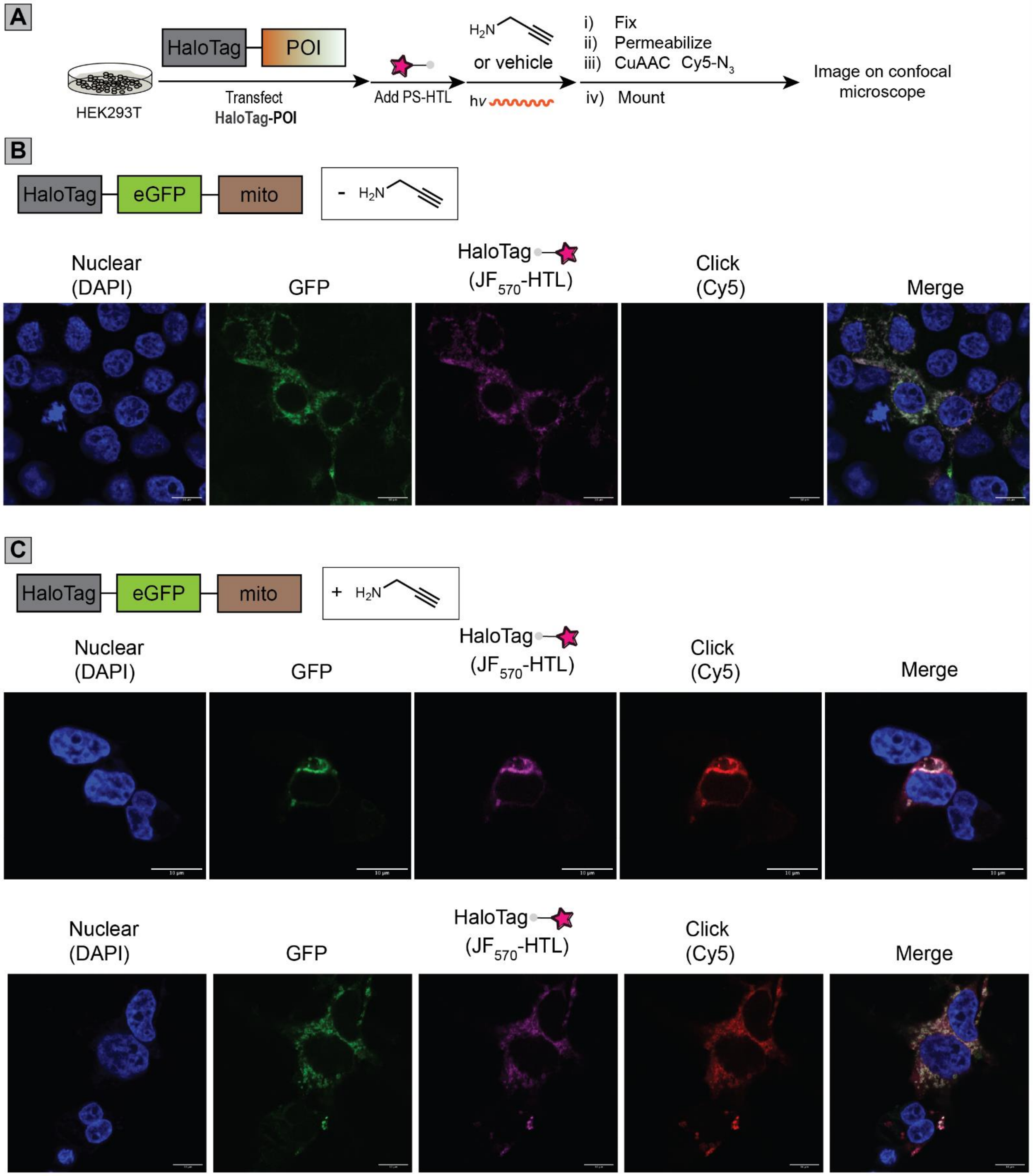


<u>Figure S9</u>: **Halo-POCA affords spatio-restricted labeling of proteins proximal to a mitochondrial-targeted HaloTag construct. (A)** Workflow for analysis of Halo-POCA labeled proteins by microscopy. **(B)** Images of individual channels or the merged channel without propargylamine present during Halo-POCA. **(C)** Images of individual channels or the merged channel when propargylamine (10 mM) is present during irradiation. HEK-293T cells transiently expressing HaloTag-eGFP-mito (pERB254[17]) subjected to the Halo-POCA procedure without (part 'B') or with (part 'C') propargylamine capture. Cells were then washed, fixed, permeabilized,

and had Cy5-azide appended via CuAAC. After washing, staining with DAPI, and mounting, cells were imaged on a Zeiss LSM880 confocal microscope. Experiment omitting propargylamine (part 'B') shows $JF_{570}$ emission does not significantly overlap with Cy5 channel. Scale bars = 10 µm.

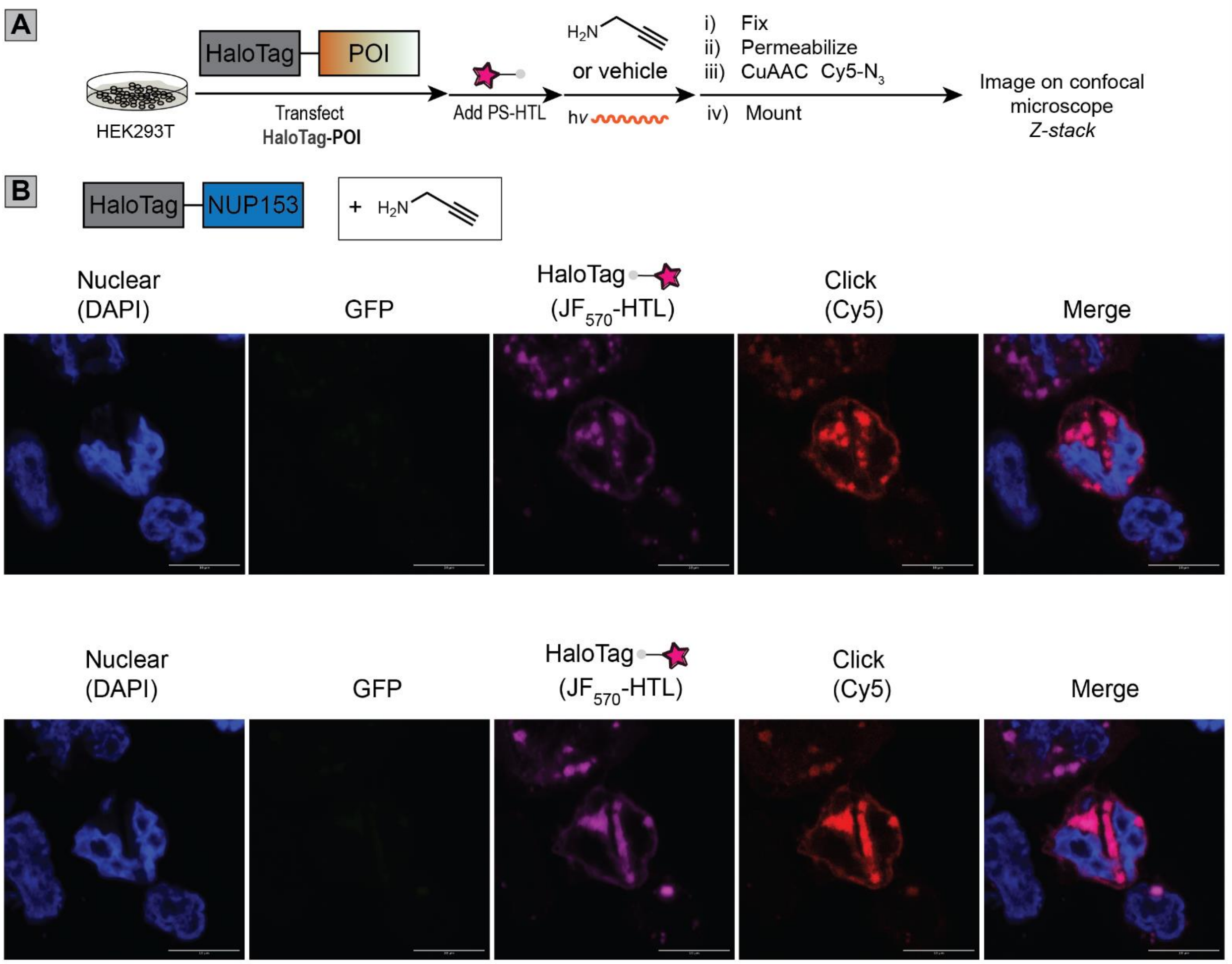


**<u>Figure S10</u>: Halo-POCA affords spatio-restricted labeling of proteins proximal to HaloTag-NUP153. (A)** Workflow for analysis of Halo-POCA labeled proteins by microscopy. **(B)** Images of individual channels or the merged channel. HEK293T cells transiently expressing HaloTag-NUP153 on poly-D-lysine-coated coverslips were taken through the Halo-POCA labeling procedure with propargylamine capture. Cells were then washed twice with PBS and fixed, permeabilized, and had Cy5-azide ($Cy5\text{-}N_3$) appended via CuAAC. After washing, staining with DAPI, and mounting, cells were imaged on a Zeiss LSM880 confocal microscope. GFP channel included for HaloTag-NUP153 experiment to indicate specificity of $JF_{570}$ signal to the magenta channel. Scale bars = 10 µm.

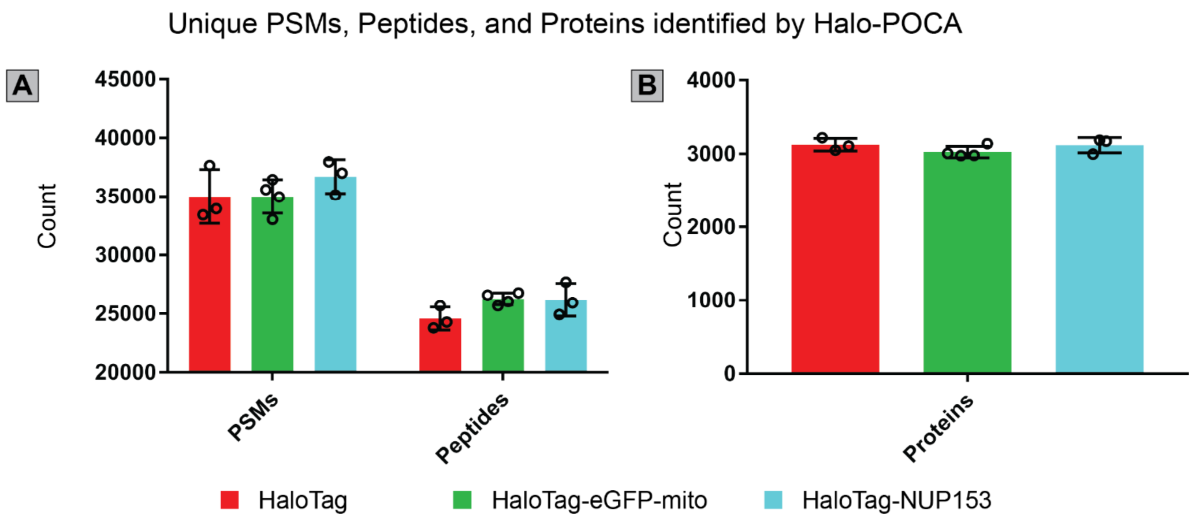


**<u>Figure S11</u>: Halo-POCA identifies similar numbers of peptide spectrum matches (PSMs), peptides, and proteins when using different HaloTag constructs. (A)** Counts of PSMs and peptides when each construct is used for Halo-POCA. **(B)** Counts of proteins when each construct is used for Halo-POCA.HEK293T cells transiently expressing either HaloTag, HaloTag-eGFP-mito (pERB254[17]), or HaloTag-NUP153 were taken through the Halo-POCA procedure. Briefly: cells were treated with $JF_{570}$-HTL (100 nM, 10 min), washed with full media, then treated with PA (10 mM) and irradiation (15 W yellow LED, 170,000 Lux max intensity, 5 min, on ice), lysis, click conjugation to biotin-azide, enrichment and MS sample preparation procedures, then the tryptic peptides were analyzed by LC-MS/MS. Closed searches were performed using FragPipe v20.0 with the *Homo sapiens* UniProtKB reviewed proteome as the database to 1% FDR. MS experiments were conducted in triplicate for each condition of +hv or –hv. All MS data can be found in **Data S1**.

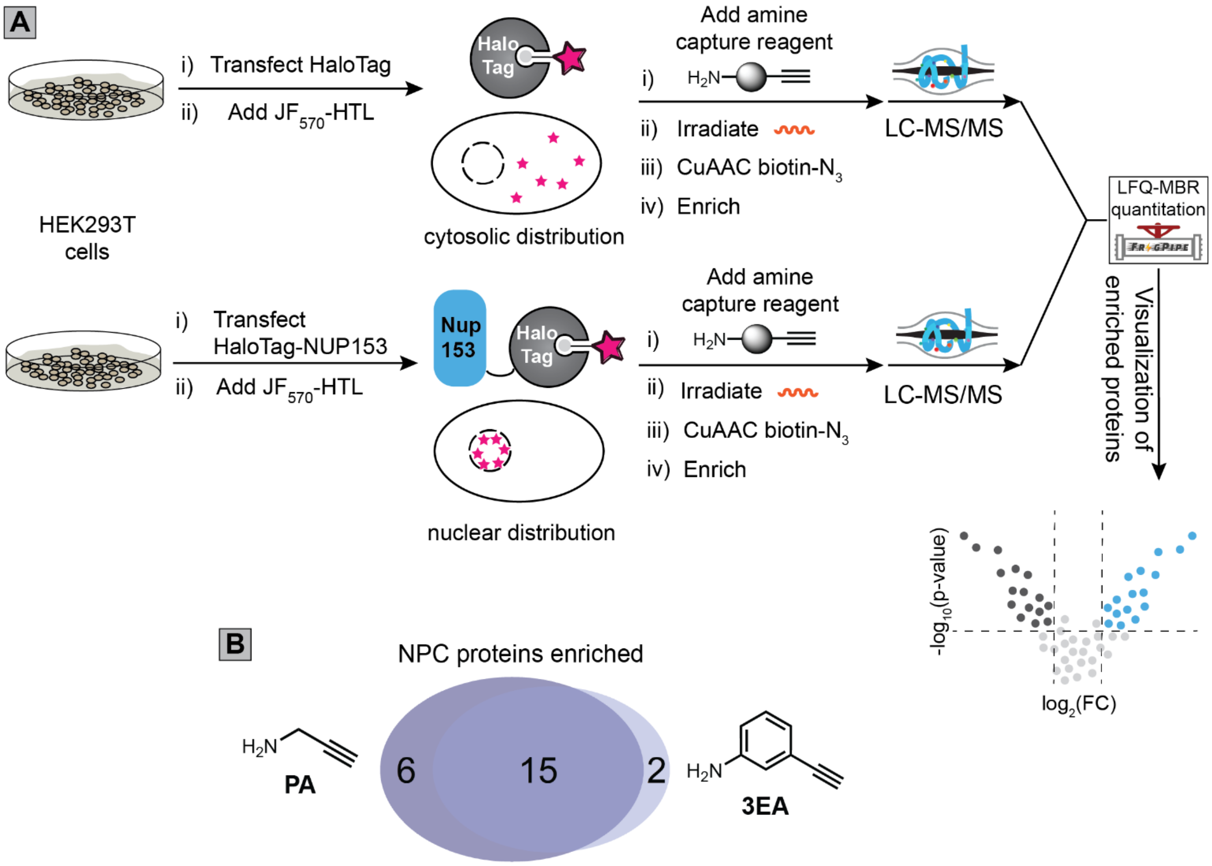


**<u>Figure S12</u>**: **Propargylamine (PA) captures a higher number of nucleoporins (NUPs) compared to 3-Ethynylaniline (3EA)**. **(A)** Halo-POCA workflow for labeling constituent proteins of the nuclear pore complex (NPC) using by comparing cells expressing either HaloTag or HaloTag-NUP153, as shown in Figure 1 of the main text. **(B)** Venn diagram of the numbers of enriched proteins when either 3-ethynylaniline (3EA) or propargylamine (PA) is used as the amine capture reagent at a concentration of 10 mM in the Halo-POCA workflow shown in part 'A'. HEK293T cells transiently expressing either HaloTag or HaloTag-NUP153 were taken through the Halo-POCA procedure. Briefly: cells were treated with $JF_{570}$-HTL (100 nM, 10 min), washed with full media, then treated with PA (10 mM) or 3EA (10 mM) and irradiation (15 W yellow LED, 170,000 Lux max intensity, 5 min, on ice), lysis, click conjugation to biotin-azide, enrichment and MS sample preparation procedures, then the tryptic peptides were analyzed by LC-MS/MS and the intensities were determined by LFQ-MBR. For measures of statistical significance used in part 'B', variances were calculated for each sample-condition pairing and a corresponding two-tailed t-test was performed to generate *p*-values. Enrichment criteria were set as raw *p*-value < 0.05 and $\log_2$(FC) > 1. MS experiments for each condition were conducted in 3 biological replicates in HEK293T cells. All MS data can be found in **Data S1**.

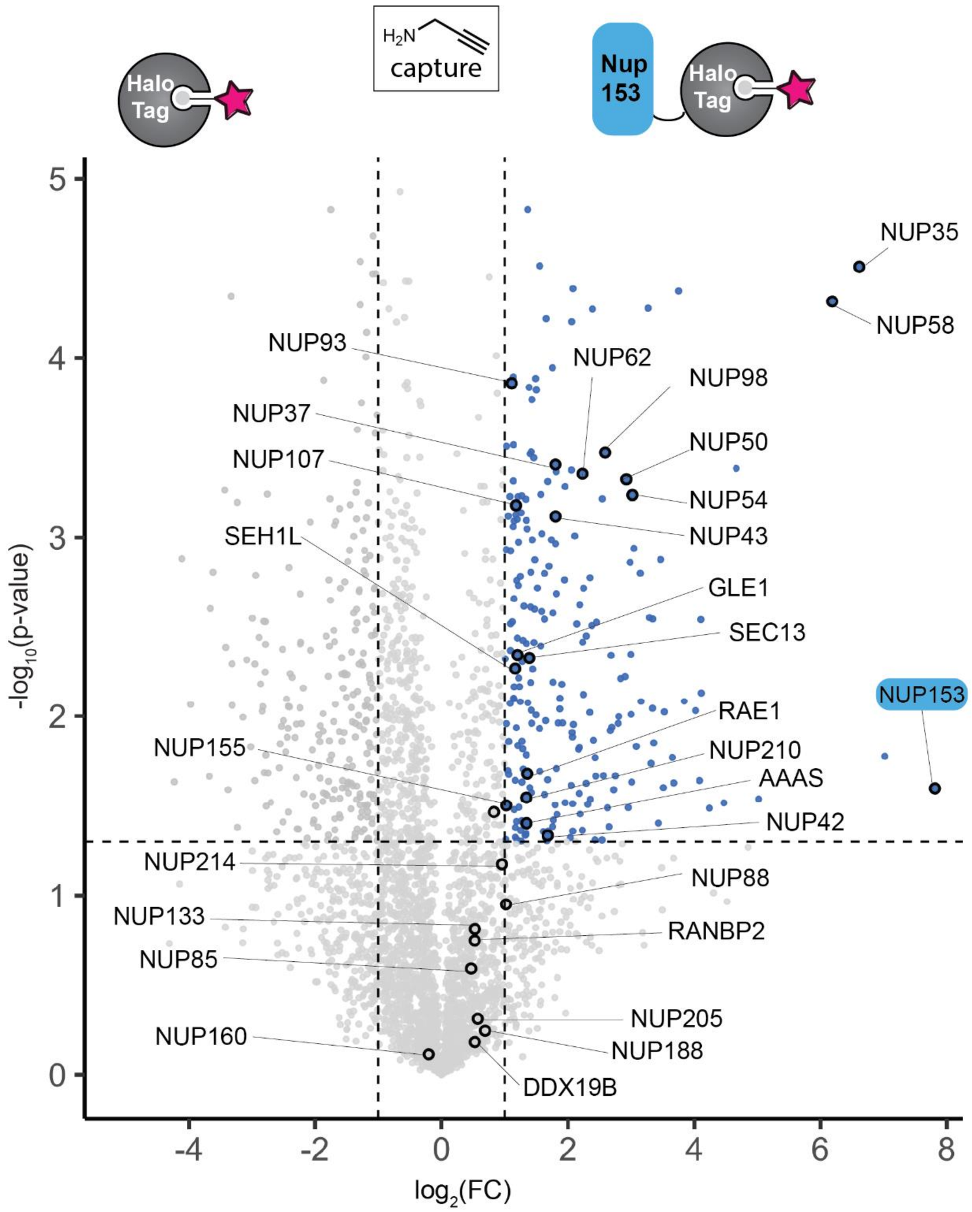


**<u>Figure S13</u>: Halo-POCA using propargylamine captures proteins of the nuclear pore complex.** Volcano plot of the proteomic data when HEK293T cells expressing either HaloTag or Myc-HaloTag-NUP153 are used for Halo-POCA labeling with propargylamine as the capture reagent. HEK293T cells transiently expressing either HaloTag or HaloTag-NUP153 were taken through the Halo-POCA procedure. Briefly: cells were treated with $JF_{570}$-HTL (100 nM, 10 min), washed with full media, then treated with PA (10 mM) and irradiation (15 W yellow LED, 170,000 Lux max intensity, 5 min, on ice), lysis, click conjugation to biotin-azide, enrichment and MS sample preparation procedures, then the tryptic peptides were analyzed by LC-MS/MS and the intensities were determined by LFQ-MBR. For measures of statistical significance, variances were calculated for each sample-condition pairing and a corresponding two-tailed t-test was performed to generate *p*-values. MS experiments were conducted in 3 replicates for each condition in HEK293T cells. All MS data can be found in **Data S1**.

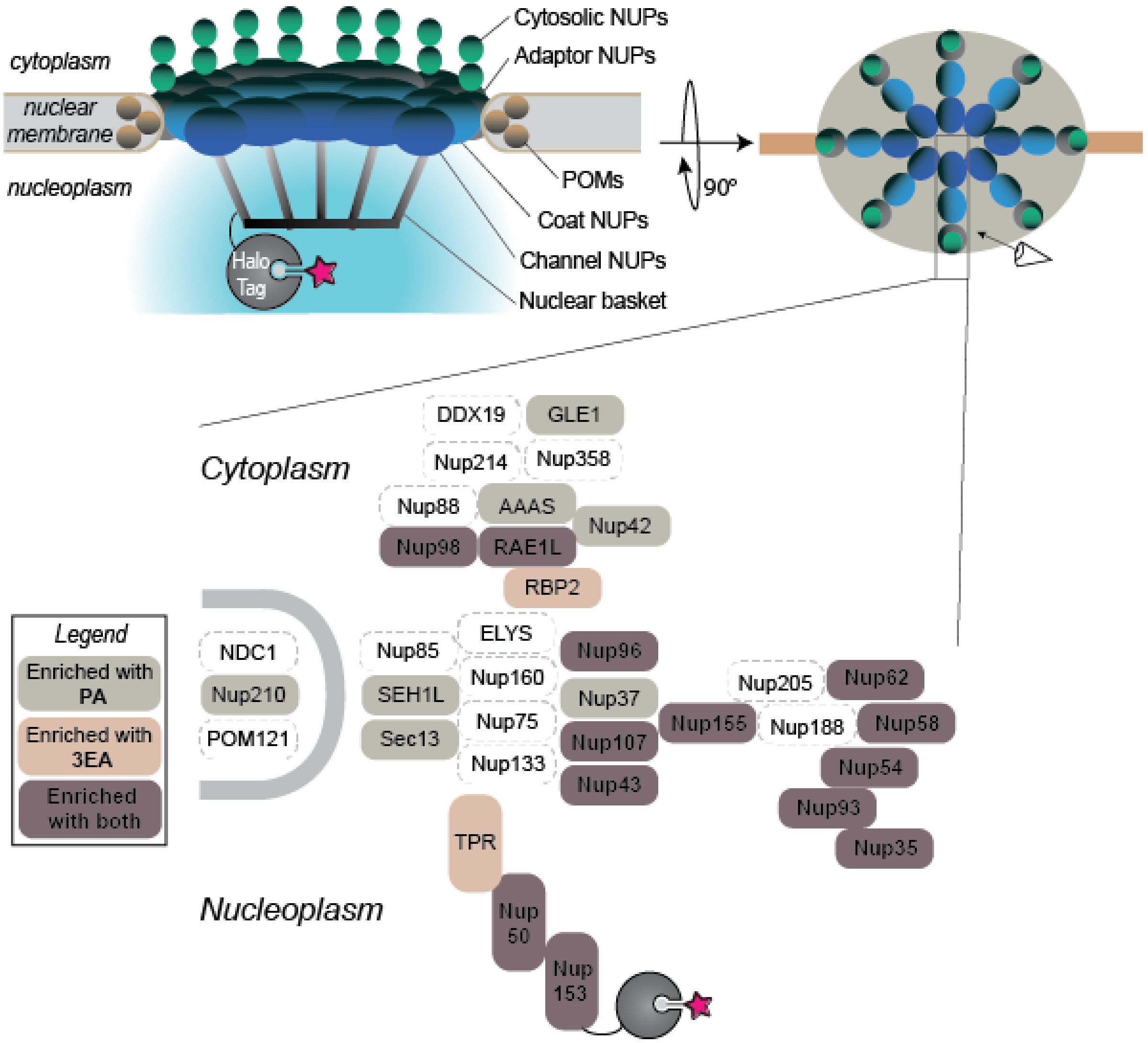


<u>**Figure S14**</u>: **The majority of enriched nucleoporins are shared between propargylamine (PA) and 3-ethynylaniline (3EA) capture using NUP153-directed Halo-POCA.** HEK293T cells transiently expressing either HaloTag or HaloTag-NUP153 were taken through the Halo-POCA procedure. Briefly: cells were treated with $JF_{570}$-HTL (100 nM, 10 min), washed with full media, then treated with PA (10 mM) or 3EA (10 mM) and irradiation (15 W yellow LED, 170,000 Lux max intensity, 5 min, on ice), lysis, click conjugation to biotin-azide, enrichment and MS sample preparation procedures, then the tryptic peptides were analyzed by LC-MS/MS and the intensities were determined by LFQ-MBR. Halo-POCA labeling with either propargylamine (PA) or 3-ethynylaniline (3EA) was compared to cells expressing the free HaloTag control. For measures of statistical significance used in part 'B', variances were calculated for each sample-condition pairing and a corresponding two-tailed t-test was performed to generate *p*-values. Enrichment criteria were set as raw *p*-value < 0.05 and $\log_2(FC) > 1$. MS experiments for each condition were conducted in 3 biological replicates in HEK293T cells. All MS data can be found in **Data S1**.

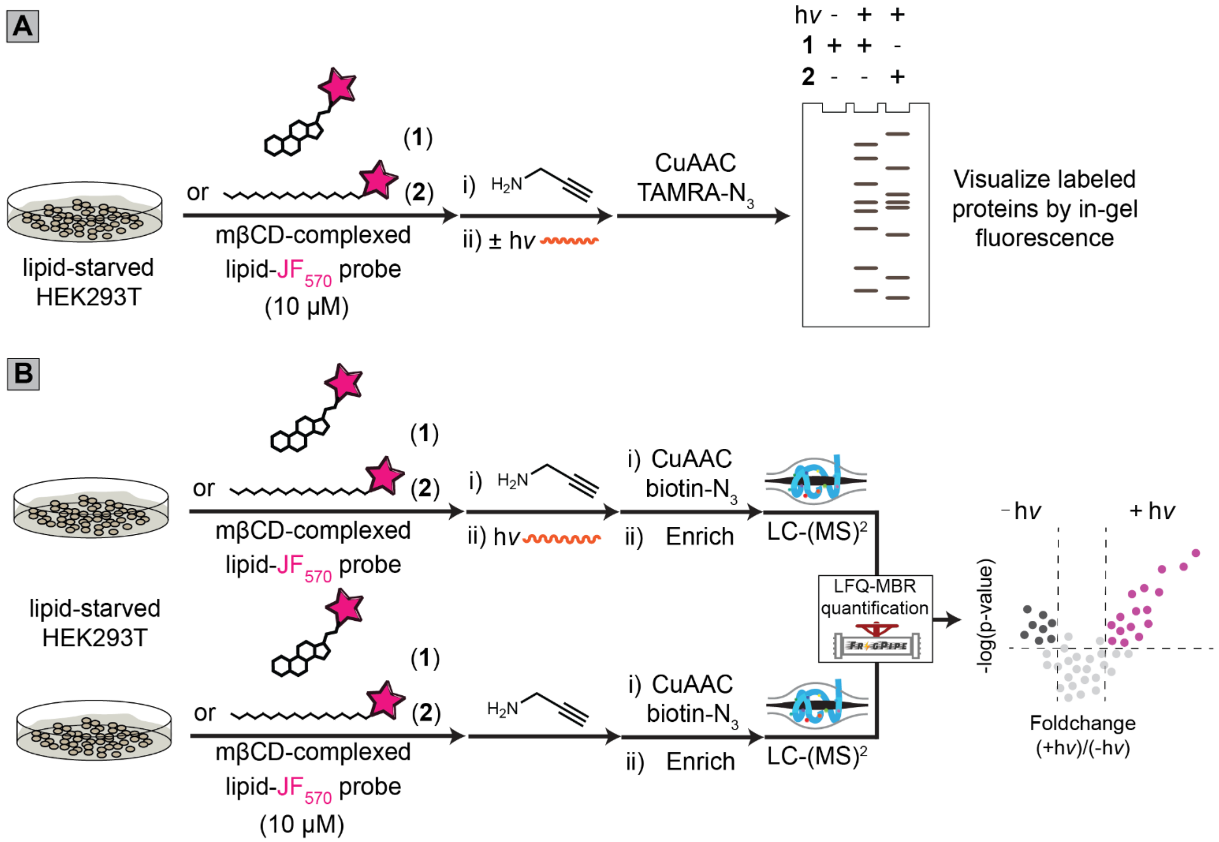


**<u>Figure S15</u>: Workflow for examining light-dependent POCA labeling of proteins *in cellulo* by small molecule probes 1 or 2.** HEK-293T cells, briefly starved of lipids by washing away complete media and incubating with serum-free media for 4 h, were treated with mβCD-cholesterol-$JF_{570}$ probe **1** or mβCD-palmitate-$JF_{570}$ probe **2** (10 µM) followed by either **(A)** gel-based assay of POCA labeling or **(B)** label-free quantification–match between runs (LFQ-MBR) proteomic analysis and quantification of labeled proteins for visualization by volcano plot.

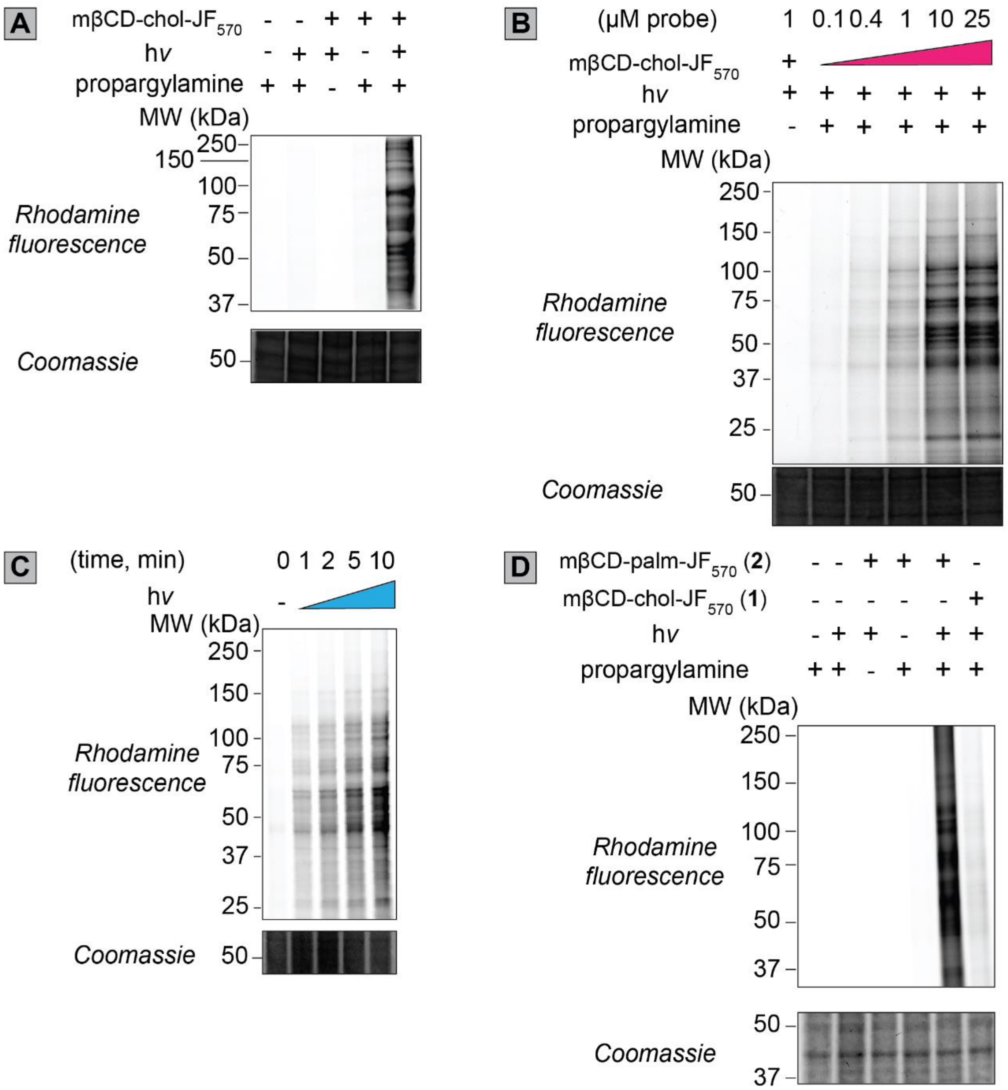


**<u>Figure S16</u>: POCA labeling with small molecule probes is dependent on the presence of probe, amine capture reagent, and irradiation. (A)** Fluorescent gel of chol-POCA when applied to HEK293T cells, with different components of the method removed. Only efficient labeling is observed when all three components (photosensitizer-containing probe, light, amine capture reagent) are present. Reaction conditions: cells treated with 10 μM mβCD-complexed chol-TOTA-$JF_{570}$ for 30 min, 5 min yellow LED irradiation (170,00 Lux), 10 mM propargylamine. **(B)** Concentration dependence of chol-POCA labeling when HEK293T cells are treated with mβCD-complexed chol-TOTA-$JF_{570}$ for 60 min and irradiated for 5 min (yellow LED) in the presence of PA (10 mM). The extent of protein labeling, inferred by fluorescence intensity, appears to saturate around 10 μM. **(C)** Irradiation time dependence of chol-POCA labeling when HEK293T cells are treated with mβCD-complexed chol-TOTA-$JF_{570}$ for 60 min and then irradiated for various amounts of time in the presence of PA (10 mM). **(D)** Comparison of labeling between mβCD-**1** (cholesterol-$JF_{570}$) and mβCD-**2** (palmitate-$JF_{570}$). HEK293T cells were treated with 10 μM mβCD-complexed chol-TOTA-$JF_{570}$ for 60 min, 5 min yellow LED irradiation (170,00 Lux), 10 mM propargylamine.

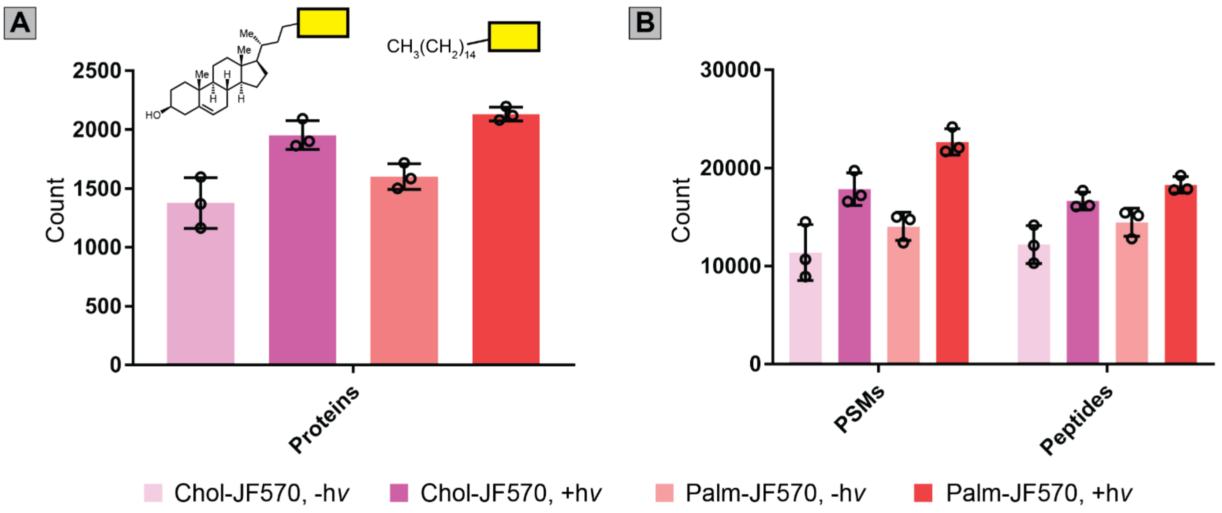


**<u>Figure S17</u>: POCA probe palmitate-$JF_{570}$ (2) effects modestly higher protein labeling than cholesterol-$JF_{570}$ (1) as assessed by proteomics. (A)** Counts of proteins identified by the cholesterol-$JF_{570}$ probe **1** (pink and light pink bars) and the palmitate-$JF_{570}$ probe **2** (red and light red bars). **(B)** Counts of peptide-spectrum matches (PSMs) and peptides identified by **1** (pink and light pink bars) and **2** (red and light red bars).Closed searches were performed using Fragpipe v20.0 with the *Homo sapiens* UniProtKB reviewed proteome as the database. Label-free quantification–match between runs (LFQ-MBR) within the IonQuant package was used to 1% FDR. MS experiments were conducted in HEK-293T cells using 3 replicates for each condition of +*hv* or –*hv*. All MS data can be found in **Data S2**.

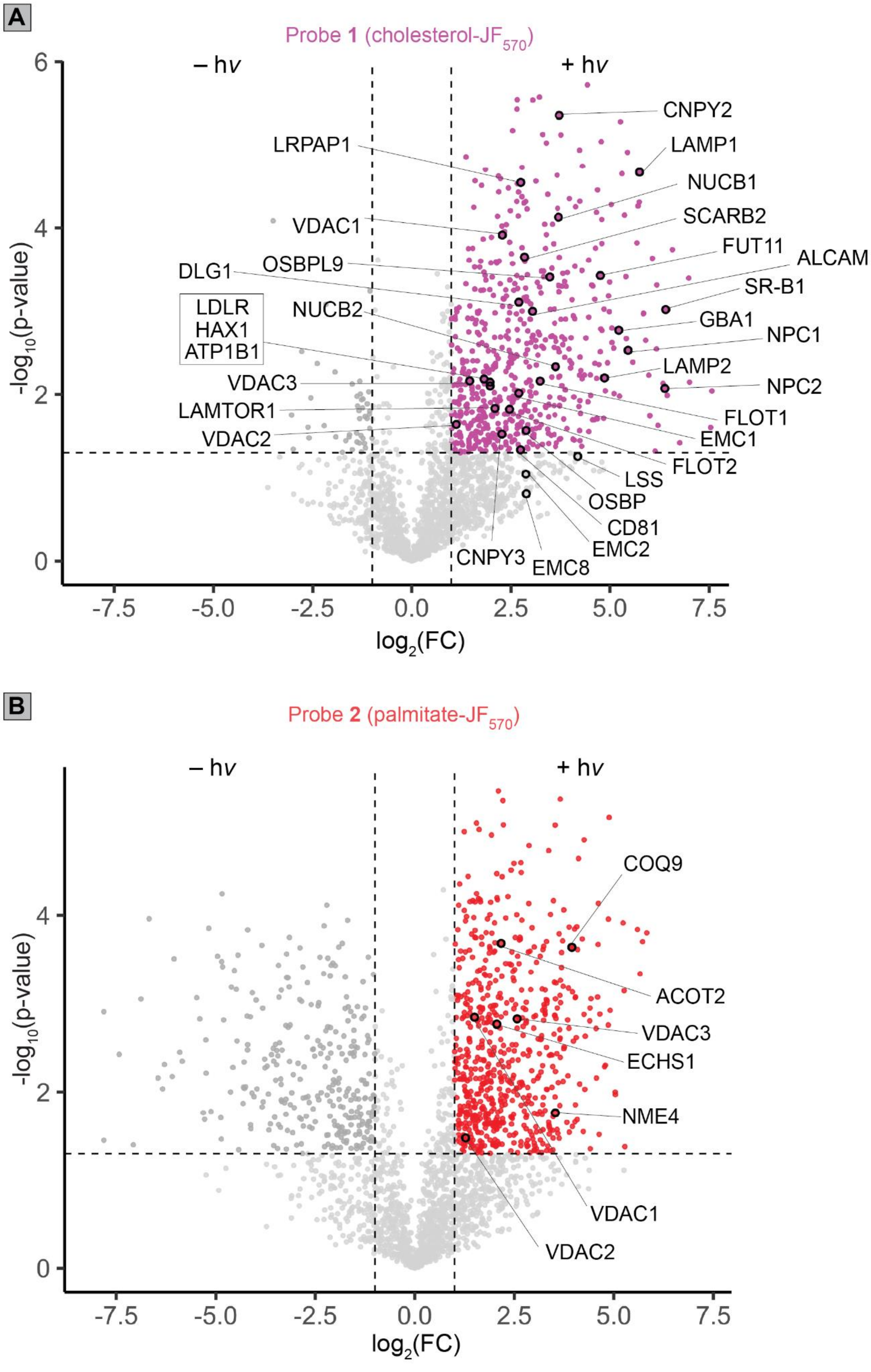


**Figure S18**: Photocatalytic POCA labeling enriches interacting proteins in a light-dependent manner using cholesterol-JF570 probe 1 (A) or palmitate-JF570 probe 2 (B).

Labeled proteins are those shown in **Figures 1E** and **1F** of the main text. HEK293T cells were treated with mβCD-**1** or **2** (10 µM, 60 min), washed twice, treated with propargylamine (10 mM, PA) and irradiation with a yellow LED (170,000 Lux, 5 min, +hv group) or kept in the dark (-hv group). After lysis, click conjugation to biotin-azide, cleanup and enrichment, tryptic digest, and LC-MS/MS analysis, protein LFQ measures of abundance were determined by label-free quantification–match between runs (LFQ-MBR)[6]. For measures of statistical significance, variances were calculated for each sample-condition pairing and a corresponding two-tailed t-test was performed to generate *p*-values. Enrichment criteria were set as raw *p*-value < 0.05 and $\log_2(FC) > 1$. MS experiments were conducted in HEK293T cells in triplicate. All MS data can be found in **Data S2**.

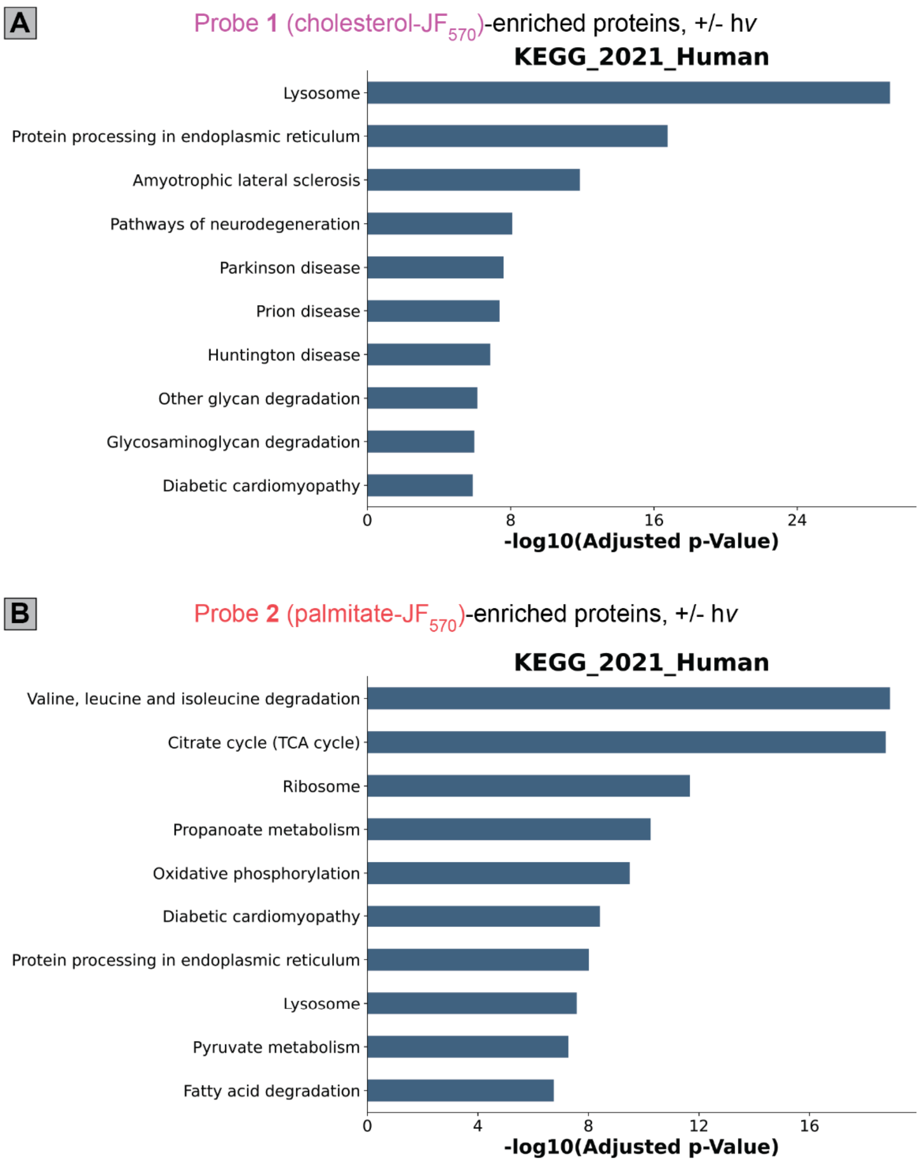


**<u>Figure S19</u>: Cholesterol-JF$_{570}$ probe 1 and palmitate-JF$_{570}$ probe 2 label proteins involved in differential cellular pathways. (A)** KEGG pathway analysis for probe **1** of the data presented in Figure 2C/2D. **(B)** KEGG pathway enrichment analysis for probe **2** of the data presented in Figure 2C/2D. Aggregate identified proteins for probe **1** in part ‘A’, or aggregate identified proteins for probe **2** in part ‘B’ were used as the background for the KEGG analyses. For measures of statistical significance, variances were calculated for each sample-condition pairing and a corresponding two-tailed t-test was performed to generate *p*-values. Enrichment criteria were set as raw *p*-value < 0.05 and $\log_2$(FC) > 1. MS experiments were conducted in HEK293T cells in triplicate. All MS data are found in **Data S2**.

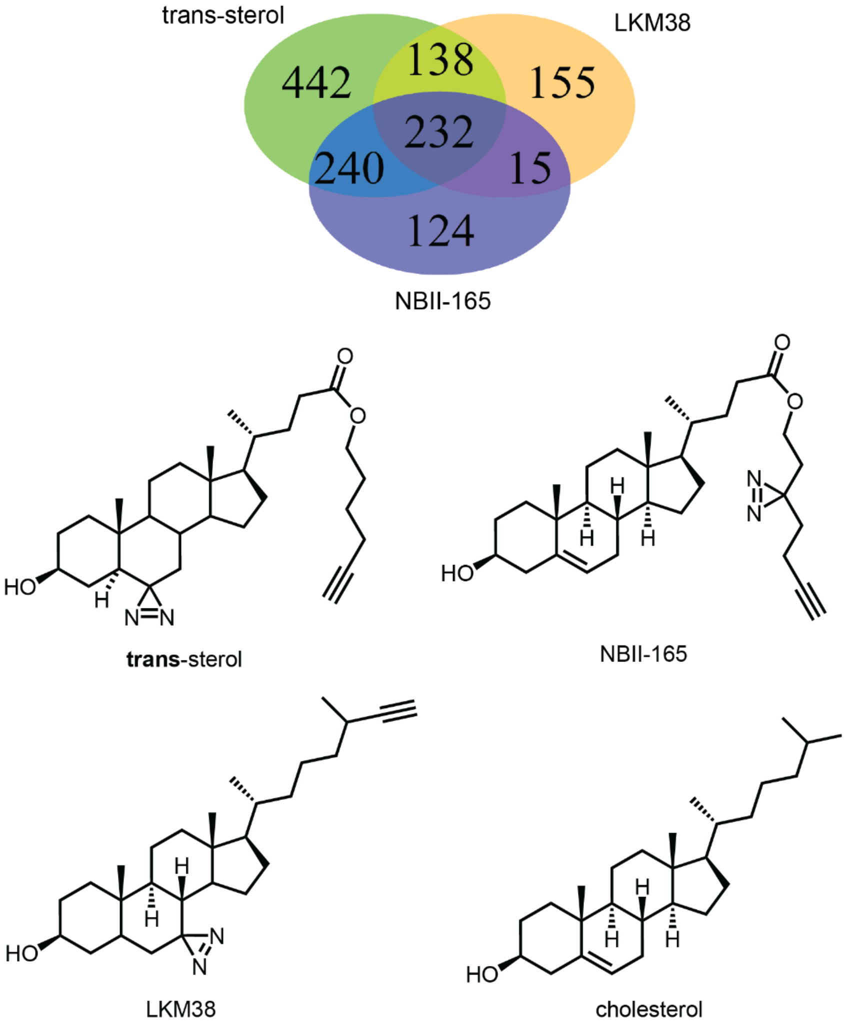


**Figure S20**: **The overlap of proteins enriched by the different cholesterol-diazirine probes [1,2] when applied to HEK-293T cells and their structures.** HEK293T cells were treated with mβCD-NBII-165, mβCD-LKM38, or mβCD-*trans*-sterol (10 µM, 60 min) and irradiation with UV light (15 min, +hv group) or kept in the dark (-hv group). After lysis, click conjugation to biotin-azide, cleanup and enrichment, tryptic digest, and LC-MS/MS analysis, protein LFQ measures of abundance were determined by label-free quantification–match between runs (LFQ-MBR) [6]. For measures of statistical significance, variances were calculated for each sample-condition pairing and a corresponding two-tailed t-test was performed to generate *p*-values. Enrichment criteria were set as raw *p*-value < 0.05 and $\log_2$(FC) > 1. MS experiments were conducted in HEK293T cells in triplicate. All MS data can be found in **Data S2**.

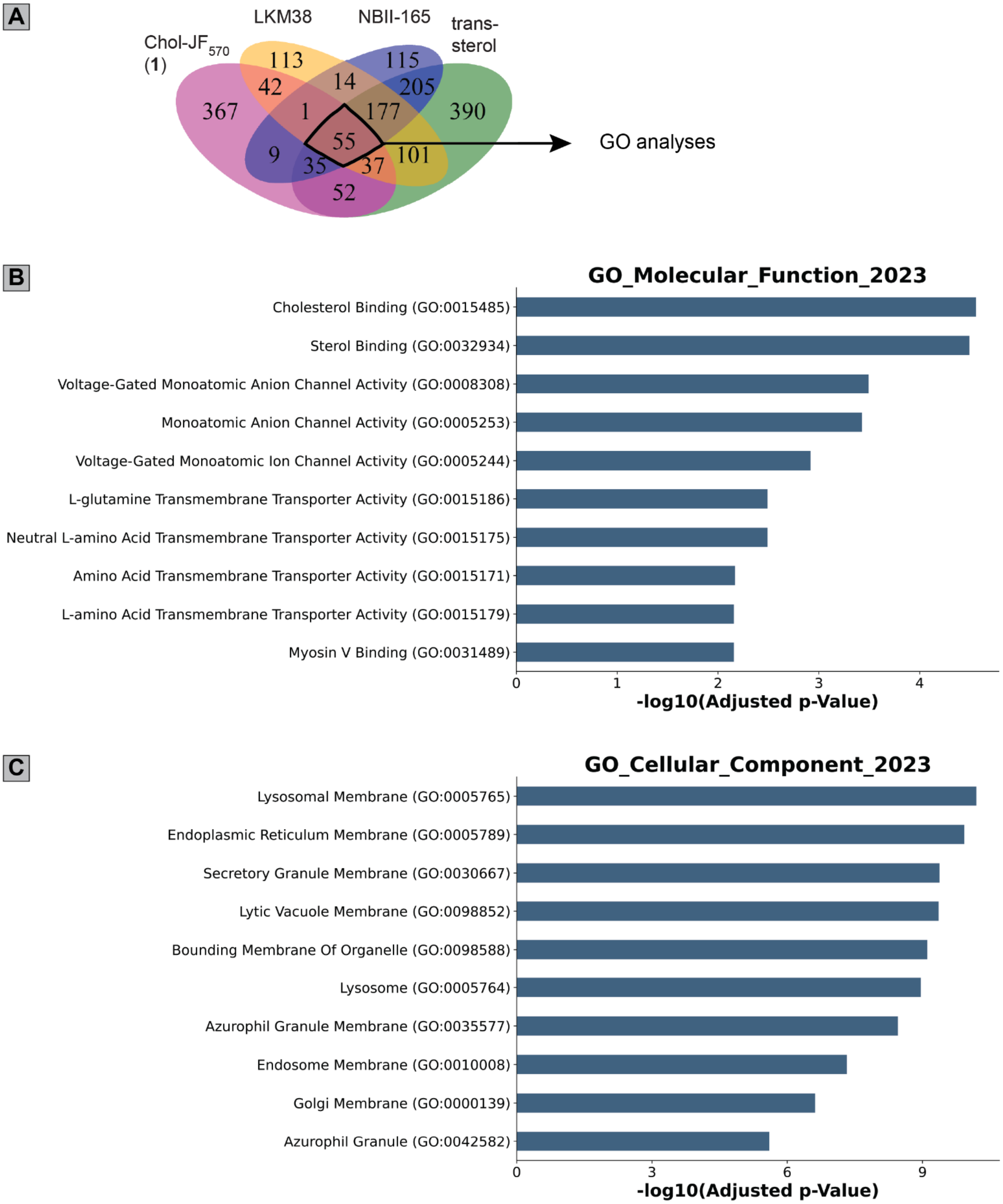
A
Chol-JF570 (1)
LKM38
NBII-165
trans-sterol
367
113
42
14
1
115
205
177
390
55
9
35
37
101
52
GO analyses
B
GO_Molecular_Function_2023
Cholesterol Binding (GO:0015485)
Sterol Binding (GO:0032934)
Voltage-Gated Monoatomic Anion Channel Activity (GO:0008308)
Monoatomic Anion Channel Activity (GO:0005253)
Voltage-Gated Monoatomic Ion Channel Activity (GO:0005244)
L-glutamine Transmembrane Transporter Activity (GO:0015186)
Neutral L-amino Acid Transmembrane Transporter Activity (GO:0015175)
Amino Acid Transmembrane Transporter Activity (GO:0015171)
L-amino Acid Transmembrane Transporter Activity (GO:0015179)
Myosin V Binding (GO:0031489)
0
1
2
3
4
-log10(Adjusted p-Value)
C
GO_Cellular_Component_2023
Lysosomal Membrane (GO:0005765)
Endoplasmic Reticulum Membrane (GO:0005789)
Secretory Granule Membrane (GO:0030667)
Lytic Vacuole Membrane (GO:0098852)
Bounding Membrane Of Organelle (GO:0098588)
Lysosome (GO:0005764)
Azurophil Granule Membrane (GO:0035577)
Endosome Membrane (GO:0010008)
Golgi Membrane (GO:0000139)
Azurophil Granule (GO:0042582)
0
3
6
9
-log10(Adjusted p-Value)

**<u>Figure S21</u>**: **Cholesterol probes enrich cholesterol-binding and membrane proteins**. **(A)** Venn diagram (replicated from Figure 2E) shows the coverage of shared and and uniquely enriched proteins captured by each probe and the 55 shared proteins enriched by all four cholesterol probes that were subjected to Gene Ontology (GO) analyses. **(B)** GO Molecular Function analysis results, and (C) GO Cellular Component analysis results. Aggregate identified proteins for all probes were used as the background for the GO analyses. For measures of statistical significance, variances were calculated for each sample-condition pairing and a corresponding two-tailed t-test was performed to generate *p*-values. MS experiments were conducted in HEK293T cells in triplicate. Enrichment criteria were set as raw *p*-value $< 0.05$ and $\log_2(FC) > 1$. All MS data can be found in **Data S2**.

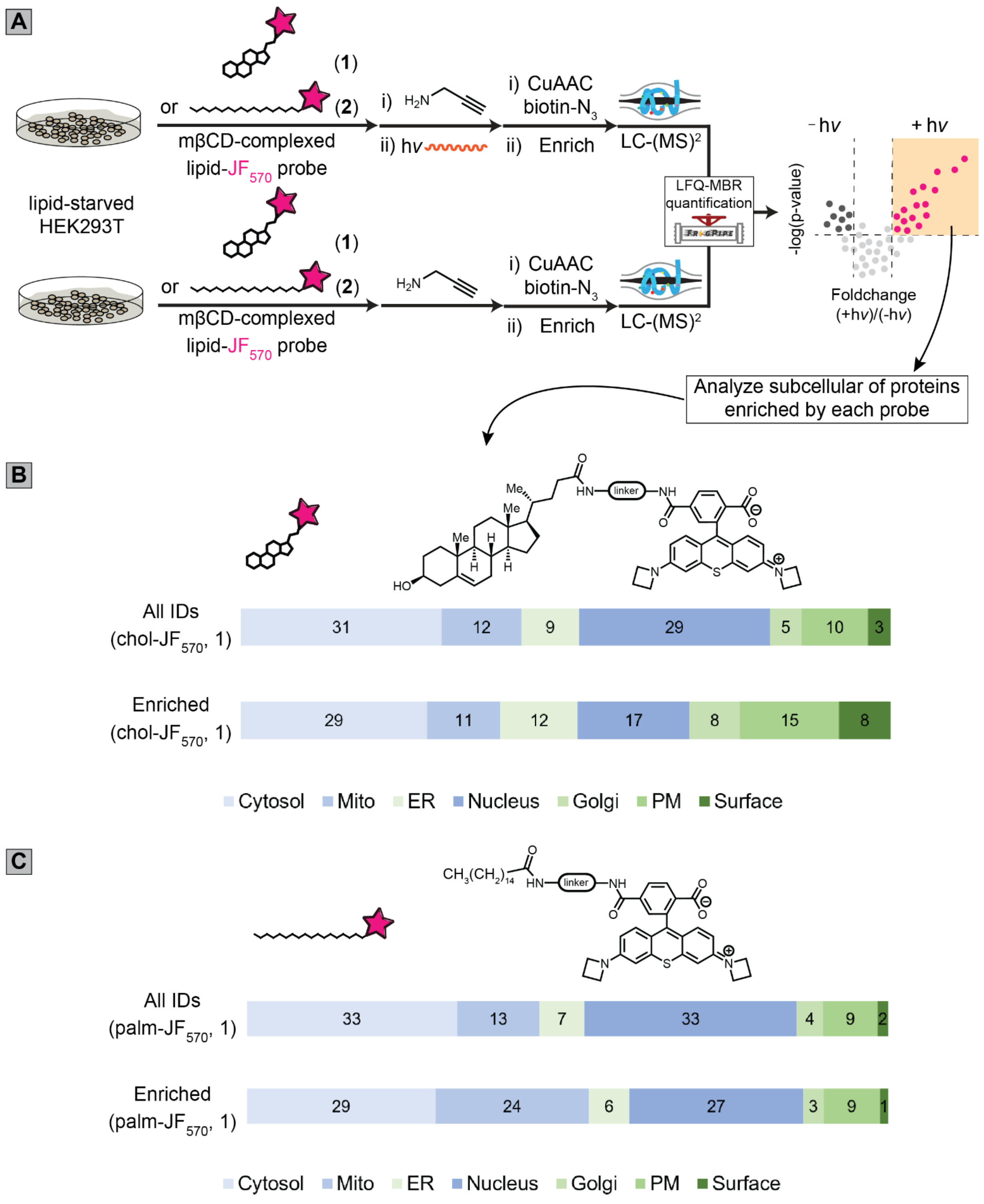


**<u>Figure S22</u>: Cholesterol-JF$_{570}$ (probe 1) and palmitate-JF$_{570}$ (probe 2) enrich distinct sets of proteins with differences in subcellular localization. (A)** Workflow for POCA-labeling with **1** and **2** and analysis of the subcellular localization of the proteins they enrich. **(B)** Subcellular localizations of all proteins identified and proteins enriched when **1** is used. **(C)** Subcellular localizations of all proteins identified and proteins enriched when **2** is used. Numbers in 'B' and

‘C’ are percentages of total. For measures of statistical significance, variances were calculated for each sample-condition pairing and a corresponding two-tailed t-test was performed to generate *p*-values. Enrichment criteria were set as raw *p*-value $< 0.05$ and $\log_2(FC) > 1$. MS experiments were conducted in HEK293T cells in triplicate. Lists used for subcellular annotations of proteins can be found in **Data S1**. MS data can be found in **Data S2**.

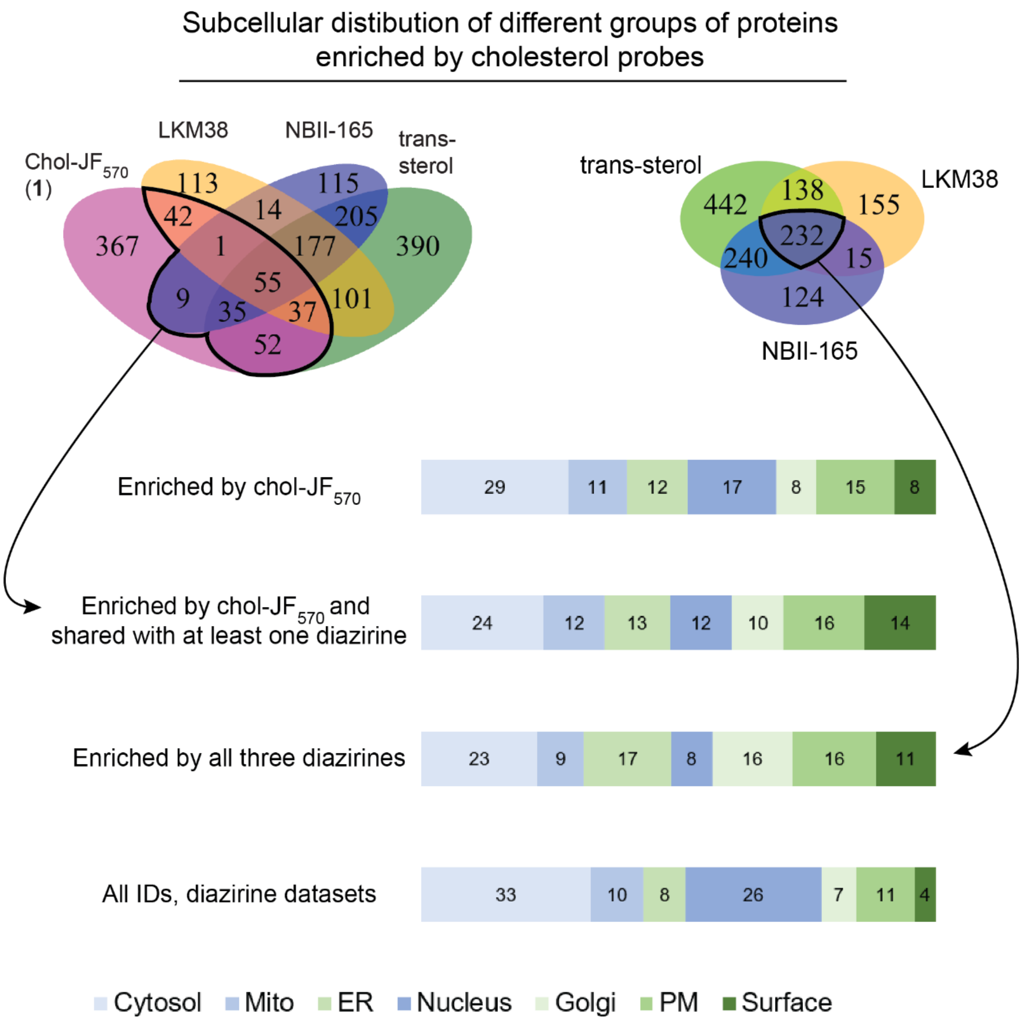


**<u>Figure S23</u>: Chol-POCA with probe 1 enriches proteins from similar cellular compartments as cholesterol-diazirine probes.** For measures of statistical significance, variances were calculated for each sample-condition pairing and a corresponding two-tailed t-test was performed to generate *p*-values. Enrichment criteria were set as raw *p*-value $< 0.05$ and $\log_2(FC) > 1$. Lists used for subcellular annotations of proteins can be found in **Data S1**. MS data can be found in **Data S2**.

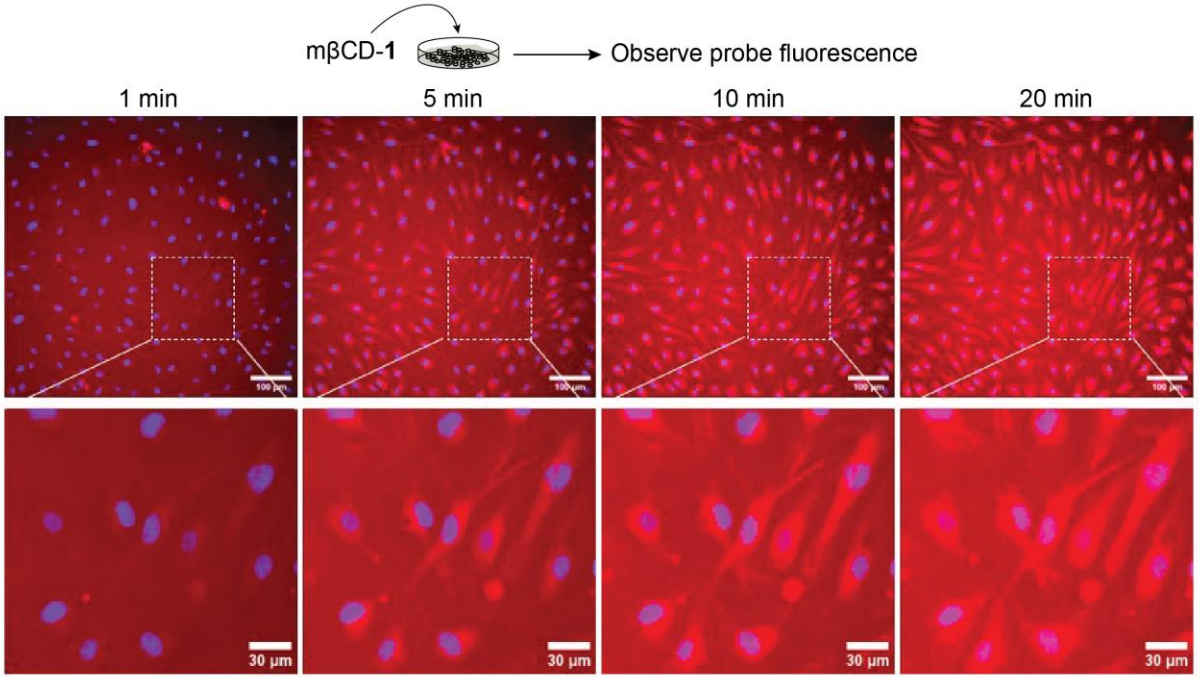


**<u>Figure S24</u>: mβCD-1 uptake and distribution are readily tracked using live-cell microscopy.** mβCD-**1** (2 μM) was added to human aortic endothelial cells and imaged at every 1 min in live cell imaging. Probe uptake was clearly visualized by time-lapse images. Scale bars = 100 μM (upper panels) and 30 μM (lower panels). Scale bars = 50 μM (upper panels) and 30 μM (lower panels). Time course is also provided as **Video S1**. Video plays at a speed of 5 frames per second, equivalent to 5 min elapsing each second.

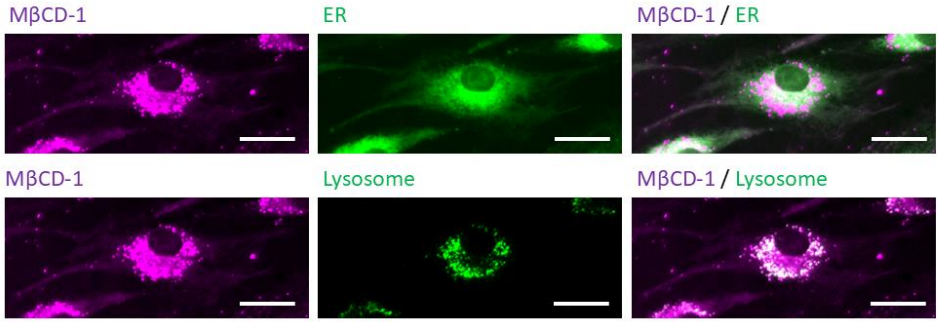


**<u>Figure S25</u>: Intracellular localization of mβCD-1 in human aortic endothelial cells.** Cells were treated with mβCD (2.5 mM, 15 min) to deplete cholesterol, washed, then treated with mβCD-1 (2 μM), ER Tracker green, and Lysotracker deep red for 30min. Then, cells were washed, and images were taken in live cell imaging. mβCD-**1** = magenta, ER = green (upper panel), Lysosome = green (lower panel). mβCD-**1** was localized to ER and lysosome in cells. Scale bars = 30 μm.

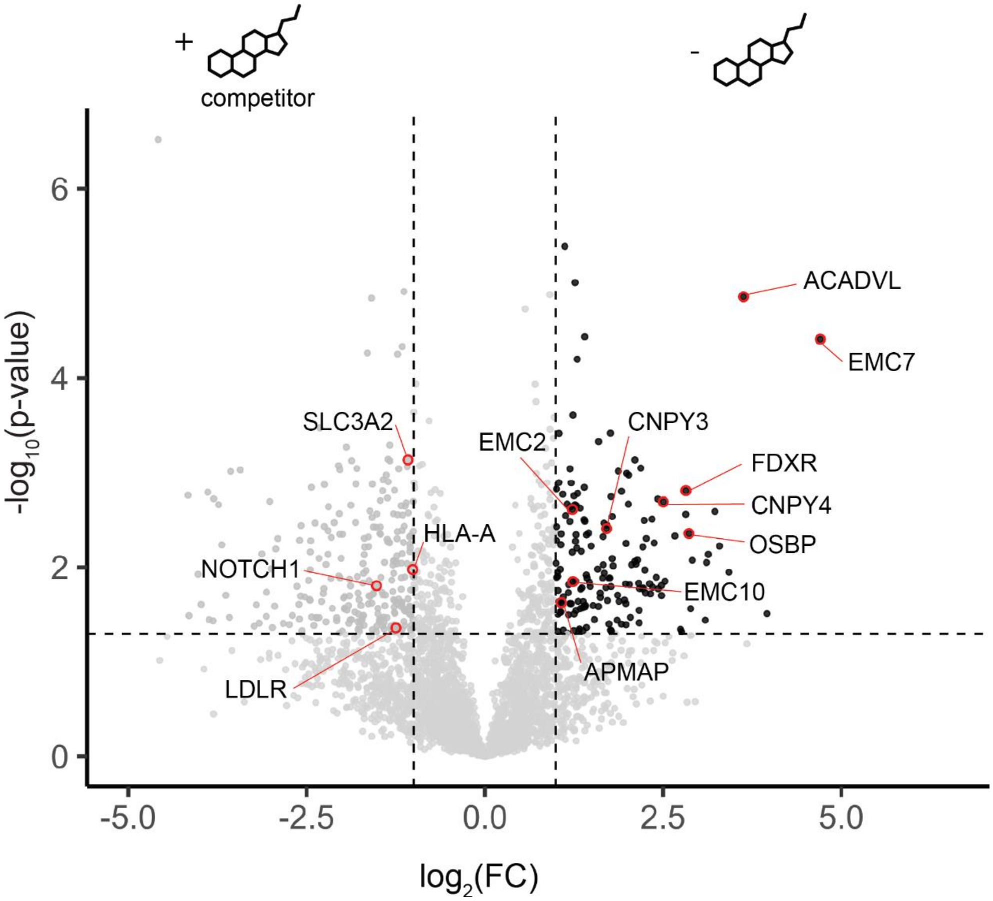


**<u>Figure S26</u>**: **Pretreatment with cholesterol competitor enables cholesterol-interacting proteins to be identified with chol-POCA.** Volcano plot of the proteomic data from treatment of HEK293T cells with mβCD-chol-TOTA-$JF_{570}$ (10 µM) and comparing the intensities of proteins labeled in the presence of 10x mβCD-cholesterol (100 µM) competitor (left side of volcano) to the intensities of proteins labeled in the absence of competitor (3 replicates, right side of volcano). Dashed lines are at the values of *p*-value = 0.05 and $\log_2$(FC) = 1. For measures of statistical significance, variances were calculated for each sample-condition pairing and a corresponding two-tailed t-test was performed to generate *p*-values. Enrichment criteria were set as raw *p*-value < 0.05 and $\log_2$(FC) > 1. MS experiments for each condition were conducted in 4 replicates (2 biological, 2 technical) in HEK293T cells. All MS data can be found in **Data S3**.

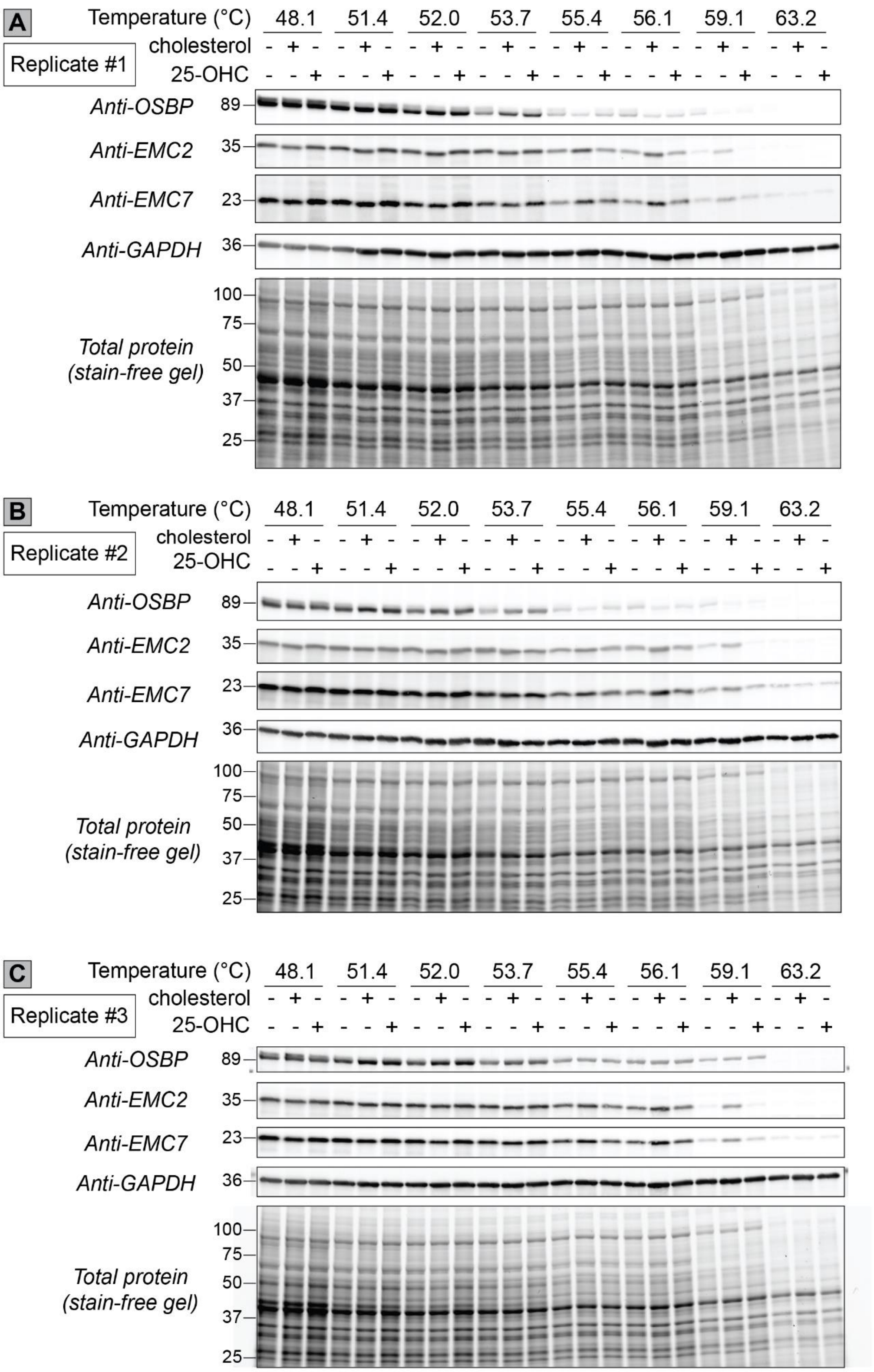


**<u>Figure S27</u>: EMC2 and EMC7 are stabilized in HEK293T cells by the addition of either mβCD-complexed cholesterol or 25-hydroxycholesterol (25-OHC).** Western blots from each replicate of in-cell CETSA analysis when HEK293T cells were treated with either mβCD-cholesterol (150 µM) or 25-OHC (10 µM) for 30 min before being heated at the indicated temperature for 3.0 min. After heating, cells were allowed to sit for 3.0 min at room temperature before they were snap frozen, thawed, and the EMC complex was solubilized by the addition of 0.4% NP40 substitute. Cells were subjected to three more freeze-thaw cycles to effect complete lysis before the lysates were cleared by centrifugation (21,000 *g* for 45 min at 4 °C). The soluble portion was normalized to the lowest concentration at each temperature point before separation by SDS-PAGE and subsequent Western blot.

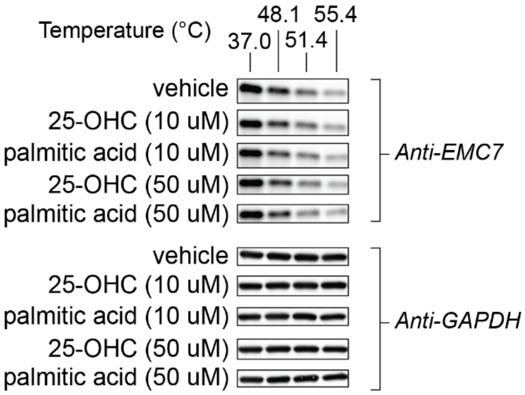


__Figure S28__: **Neither 25-hydroxycholesterol nor palmitic acid confer thermal stabilization to EMC7 in lysates.** HEK293T lysates (3.0 mg/mL, containing 0.4% nonionic surfactant) were heated at the indicated temperatures for 3.0 min and the protein abundance of EMC7 was assessed by Western blot. Protein abundance of EMC7 and GAPDH (loading control) when lysates were heated with the indicated concentrations of 25-OHC or palmitic acid.

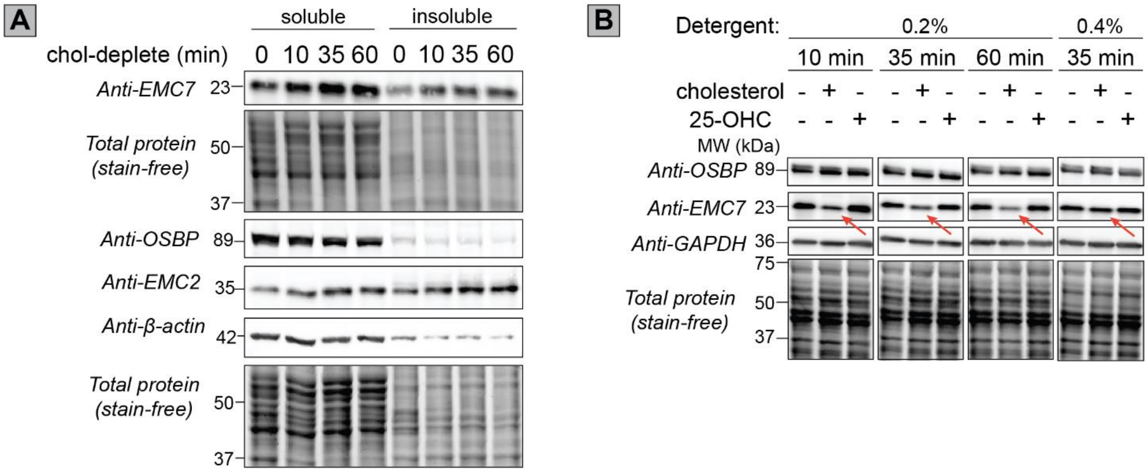


__Figure S29__: **Addition or subtraction of cholesterol leads to a time-dependent change in the amount of soluble EMC subunit(s). (A)** Immunoblots of EMC7, EMC2, and OSBP abundance as a function of time after cholesterol extraction by mβCD. HEK293T cells were treated with 2.5 mM mβCD for the indicated time, harvested, and lysed in the presence of 0.4% NP-40. **(B)**

Immunoblots of EMC7 and OSBP abundance as a function of time, cholesterol treatment, or treatment with 25-hydroxycholesterol (25-OHC, 10 μM). HEK293T cells were loaded with mβCD-cholesterol (150 μM) or 25-OHC for the indicated time, harvested, and lysed in the presence of 0.2% or 0.4% NP-40.

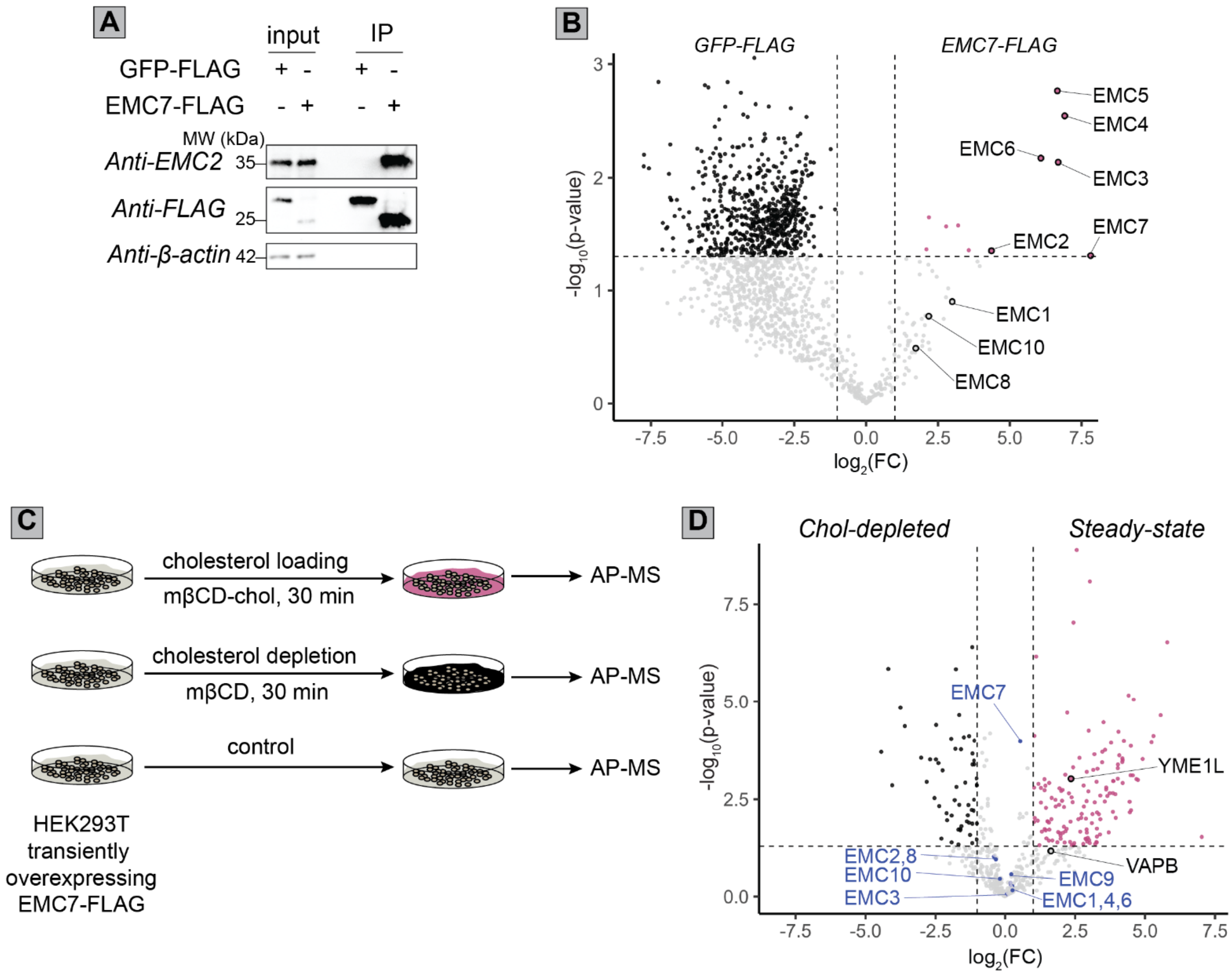


**<u>Figure S30</u>: Affinity-purification of the EMC reveals cholesterol-dependent interacting partners. (A)** Western blot showing the efficient pulldown of EMC2 using EMC7-FLAG as bait. **(B)** Plot showing the fold-changes and statistical measures of protein intensities from label-free quantification-match between runs [6] (LFQ-MBR) after affinity purification-mass spectrometry analysis using transiently-expressed GFP-FLAG or EMC7-FLAG as bait in HEK293T cells. **(C)** Plot showing the fold-changes and statistical measures of protein intensities from LFQ-MBR after affinity purification-mass spectrometry analysis using transiently expressed EMC7-FLAG bait in HEK293T cells that were either depleted of cholesterol by treatment with methyl-beta cyclodextrin (2.5 mM, 30 min) or left untreated. For measures of statistical significance, variances were calculated for each sample-condition pairing and a corresponding two-tailed t-test was performed to generate *p*-values. Enrichment criteria were set as raw *p*-value < 0.05 and $\log_2$(FC) > 1. MS experiments were conducted in HEK293T cells, n=3 for part 'B' and n=6 for part 'C'. All MS data can be found in **Data S3**.

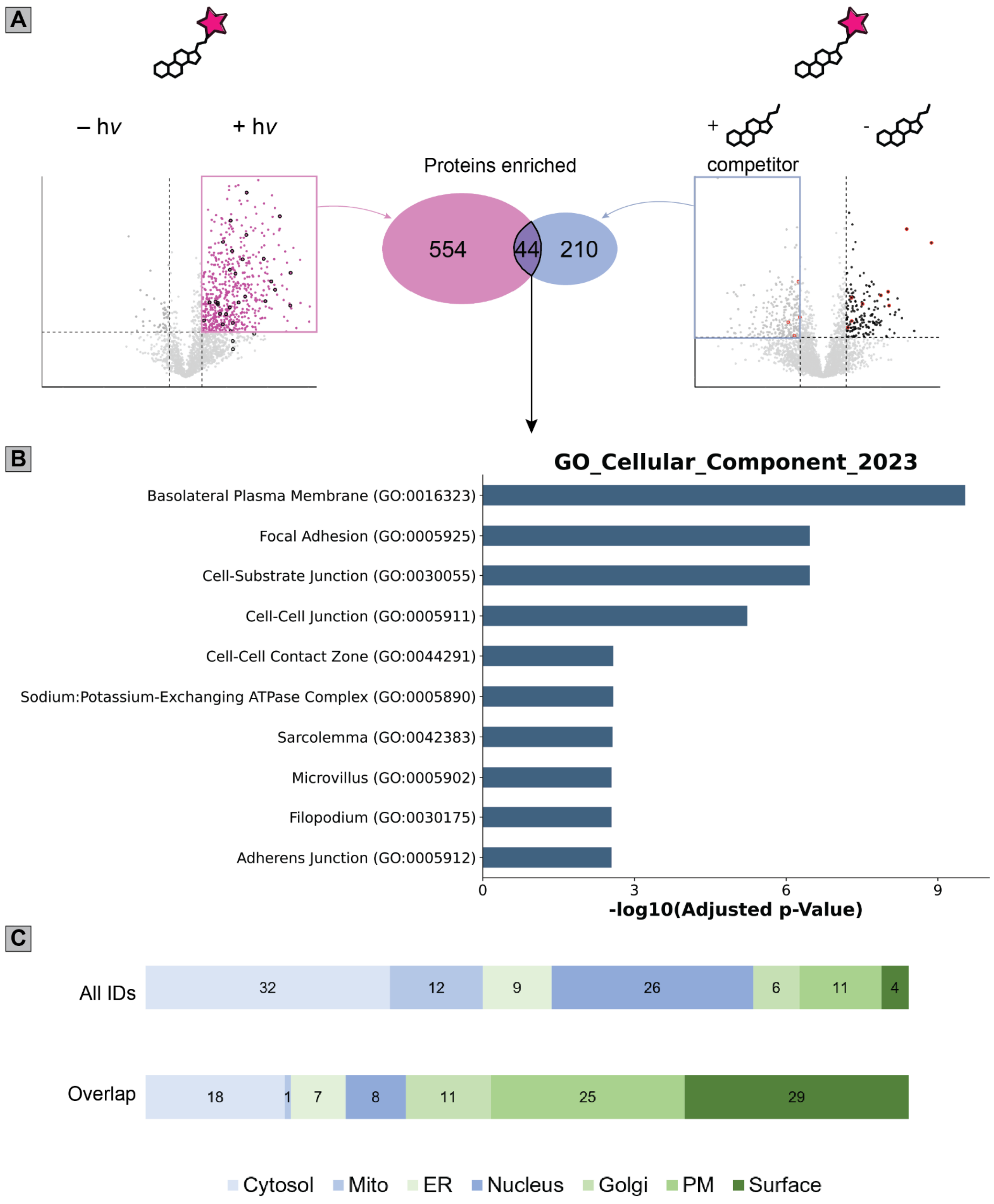


<u>Figure S31</u>: **Proteins preferentially enriched in the presence of cholesterol competitor are predominantly PM- or surface-annotated. (A)** Schematic showing the overlap of the proteins being analyzed in this figure. The left side depicts the proteins enriched by probe **1** in a light-dependent manner (data shown in **Figure 2** of the main text) while the right side depicts the proteins enriched by probe **1** in the presence of cholesterol competitor. **(B)** Cellular compartment

analysis (GO Cellular Compartment 2023) of the overlapping proteins. **(C)** Subcellular distribution of proteins enriched by probe **1** in a light-dependent manner and the proteins that had significantly more labeling in the presence of 100 µM mβCD-cholesterol competitor by mβCD-chol-$JF_{570}$ probe **1**. The increased labeling of plasma membrane proteins is hypothesized to occur due to the effects of the added cholesterol on the membrane. Aggregate identified proteins of both the competition and labeling experiments were used as the background for the GO analysis. For measures of statistical significance, variances were calculated for each sample-condition pairing and a corresponding two-tailed t-test was performed to generate *p*-values. Enrichment criteria were set as raw $p$-value < 0.05 and $\log_2$(FC) > 1. Competition experiments (+/- cholesterol) were performed with 4 replicates (2 biological, 2 technical) in HEK293T cells and labeling experiments (+/- hv) used 6 replicates in HEK293T cells. All MS data can be found in **Data S3**.

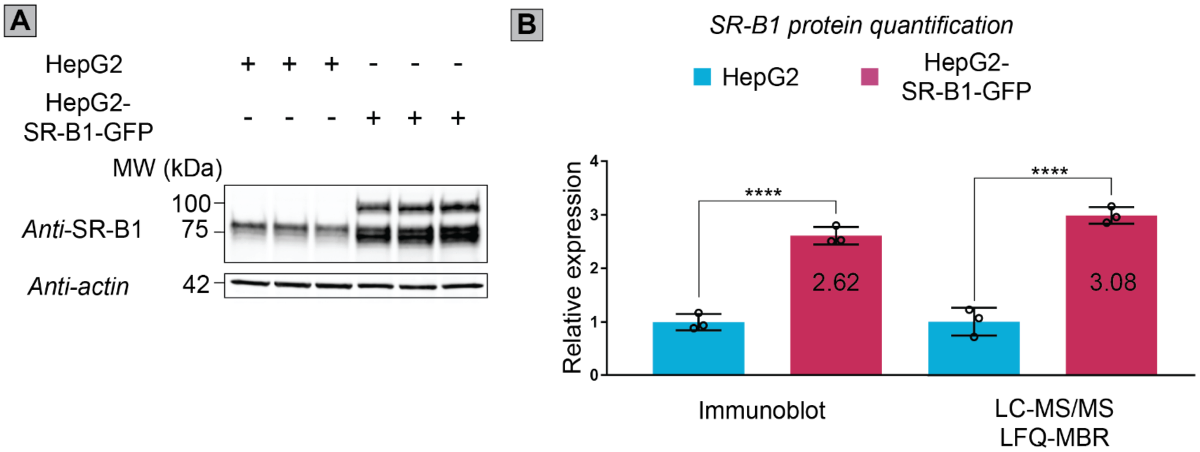


**Figure S32**: **Scavenger receptor class B member 1 (SR-B1) expression is increased in the HepG2-SR-B1-GFP cells relative to parental HepG2. (A)** Immunoblot showing the differences in SR-B1 expression. The same figure is shown in **Figure 4C** of the main text. **(B)** Quantification of the immunoblot and the LC/MS-MS LFQ-MBR measures of SR-B1 abundance. Immunoblot signals were quantified using the grayscale pixel densities, and the LC-MS/MS data were quantified by using the MS1 intensities (LFQ-MBR). In 'B', the mean value (n=3) and standard deviations are shown as the heights and error bars, respectively. Statistical significance was calculated with unpaired Student's t-tests. **** $p < 0.001$. MS data can be found in **Data S4**.

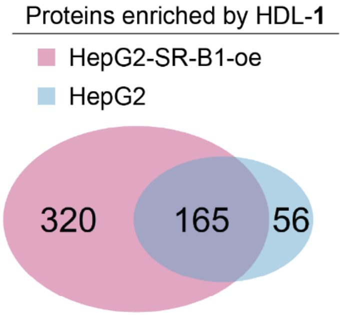


**Figure S33**: **Different and shared proteins are enriched by POCA with HDL-1 in HepG2 and HepG2 cells overexpressing the HDL ligand scavenger receptor class B member 1 (SR-B1).** Venn diagram of proteins enriched by HDL-POCA using chol-$JF_{570}$ probe **1** in either HepG2-

SR-BI-overexpressing cells (pink) or parental HepG2 cells (blue). For measures of statistical significance, variances were calculated for each sample-condition pairing and a corresponding two-tailed t-test was performed to generate *p*-values. Enrichment criteria were set as raw *p*-value < 0.05 and $\log_2$(FC) > 1. All MS data can be found in **Data S4**.

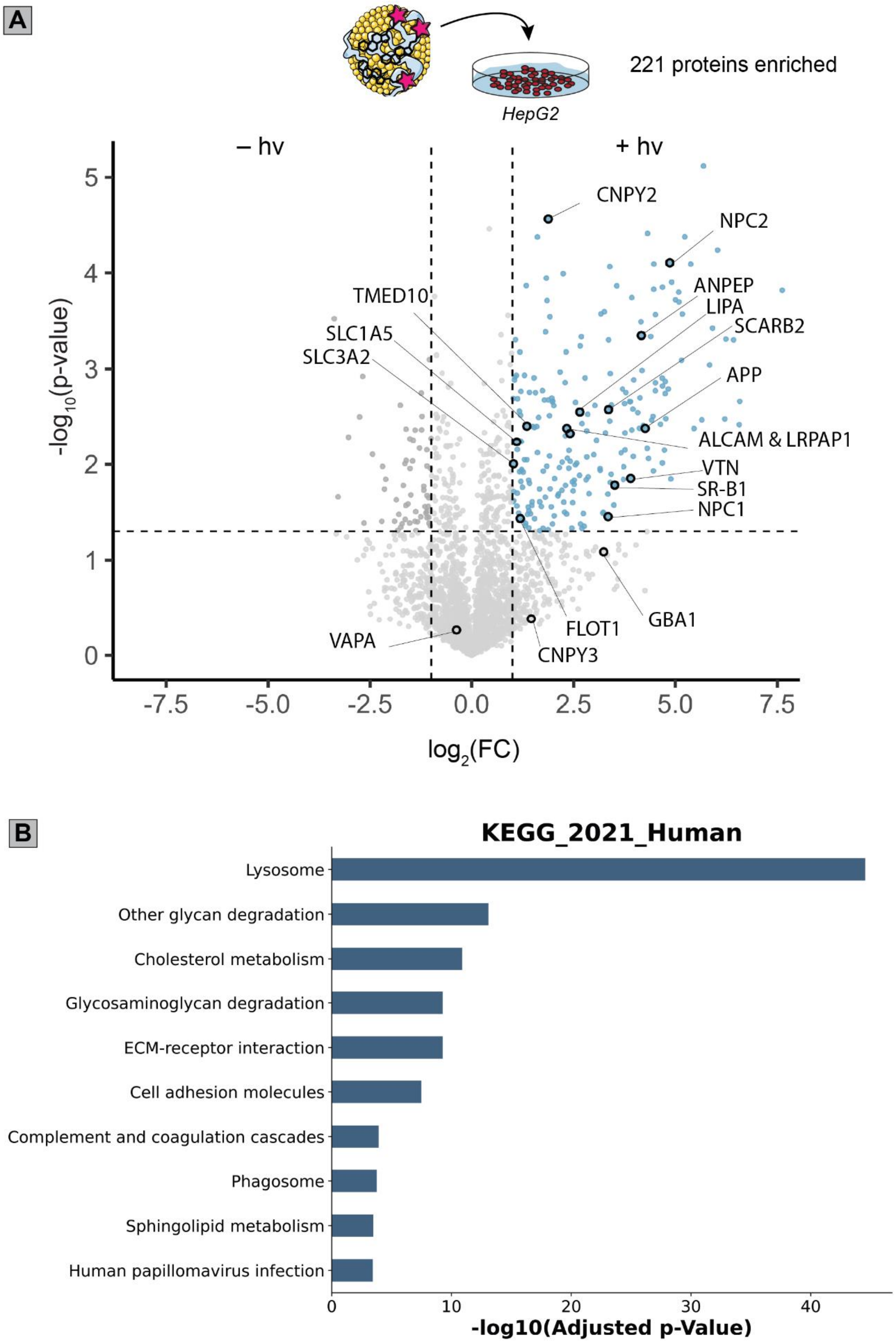

<u>Figure S34</u>: **Canonical cholesterol handling proteins are enriched by POCA using HDL-1 in parental HepG2 cells. (A)** Volcano plot of the data. **(B)** Pathway analysis of the enriched proteins (light blue data points in part 'A'). Labeled proteins are a combination of proteins enriched by mβCD-**1** in **Figure 2** of the main text and other proteins with known cholesterol associations. HepG2 cells were starved overnight of cholesterol then treated with HDL-**1** (100 ug/mL, 60 min), washed twice, treated with propargylamine (10 mM) and irradiation (15 W yellow LED, 170,000 Lux max intensity, 5 min, on ice), lysis, click conjugation to biotin-azide, enrichment and MS sample preparation procedures, then the tryptic peptides were analyzed by LC-MS/MS and the intensities were determined by LFQ-MBR [6]. Aggregate identified proteins in the experiment were used as the background for the KEGG analysis. For measures of statistical significance, variances were calculated for each sample-condition pairing and a corresponding two-tailed t-test was performed to generate *p*-values. Enrichment criteria were set as raw $p$-value < 0.05 and $\log_2(FC) > 1$. MS experiments for each condition were conducted in triplicate in HepG2 cells. All MS data can be found in **Data S4**.

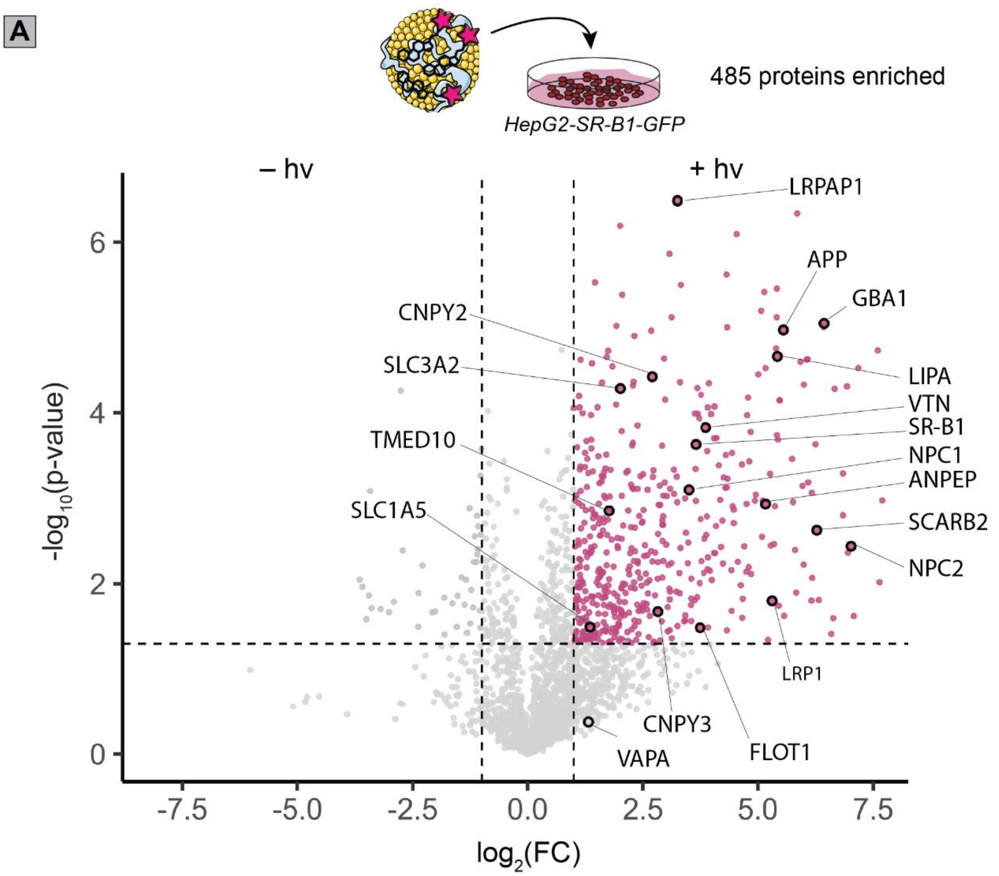


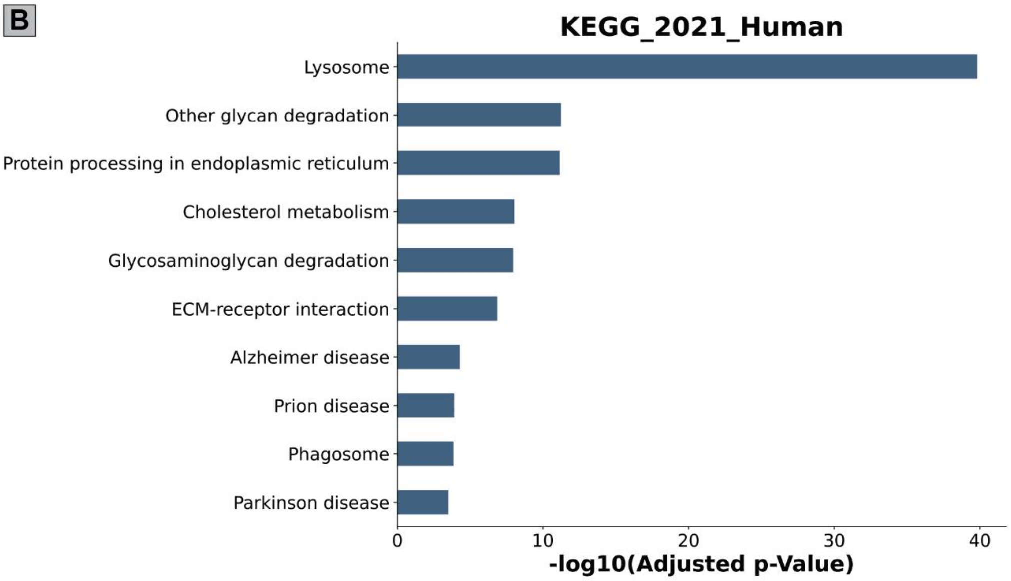


__Figure S35__: **Canonical cholesterol handling proteins are enriched by POCA using HDL-1 in HepG2 cells overexpressing SR-B1. (A)** Volcano plot of the data. **(B)** Pathway analysis of the enriched proteins (pink data points in part 'A'). Labeled proteins are a combination of proteins enriched by mβCD-**1** in **Figure 2** of the main text and other proteins with known cholesterol associations. HepG2-SR-B1-overexpressing cells were starved overnight of cholesterol then treated with HDL-**1** (100 ug/mL, 60 min), washed twice, treated with propargylamine (10 mM) and irradiation (15 W yellow LED, 170,000 Lux max intensity, 5 min, on ice), lysis, click conjugation to biotin-azide, enrichment and MS sample preparation procedures, then the tryptic peptides were analyzed by LC-MS/MS and the intensities were determined by LFQ-MBR[6]. Aggregate identified proteins in the experiment were used as the background for the KEGG analysis. For measures of statistical significance, variances were calculated for each sample-condition pairing and a corresponding two-tailed t-test was performed to generate *p*-values. Enrichment criteria were set as raw *p*-value < 0.05 and $\log_2$(FC) > 1. MS experiments for each condition were conducted in 3 biological replicates in HepG2 cells. All MS data can be found in **Data S4**.

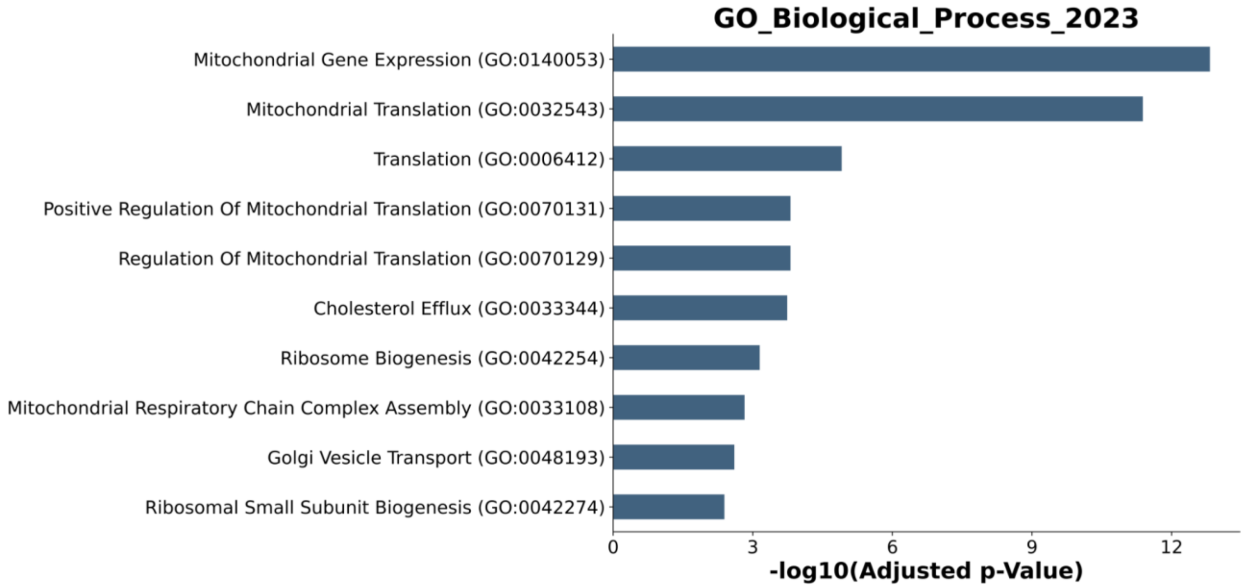


**Figure S36**: **HepG2-SR-B1-overexpressing cells upregulate mitochondrial and cholesterol efflux pathways relative to parental HepG2.** Pathway analysis (GO Biological Process 2023) of the 148 proteins with increased expression in HepG2-SR-BI-overexpressing cells (Figure 4 of the main text). Aggregate identified proteins in the experiment were used as the background for the KEGG/GO analysis. For measures of statistical significance, variances were calculated for each sample-condition pairing and a corresponding two-tailed t-test was performed to generate *p*-values. Enrichment criteria were set as raw *p*-value < 0.05 and log2(FC) > 1. All MS data can be found in **Data S4**.

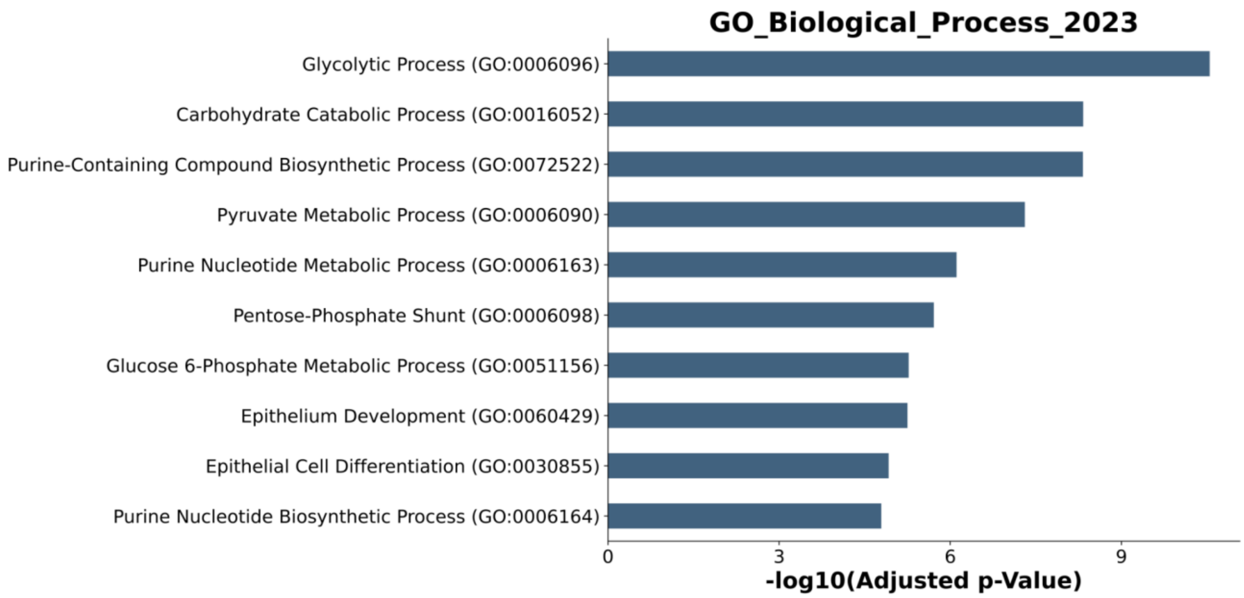


<u>Figure S37</u>: **HepG2 cells upregulate glycolytic and other metabolic pathways relative to the HepG2-SR-B1-overexpressing cells.** Pathway analysis (GO Biological Process 2023) of the 573 proteins with decreased expression in HepG2-SR-BI-overexpressing cells (Figure 4 of the main text). Aggregate identified proteins in the experiment were used as the background for the GO analysis. For measures of statistical significance, variances were calculated for each sample-condition pairing and a corresponding two-tailed t-test was performed to generate *p*-values. Enrichment criteria were set as raw *p*-value < 0.05 and $\log_2$(FC) > 1. All MS data can be found in **Data S4**.

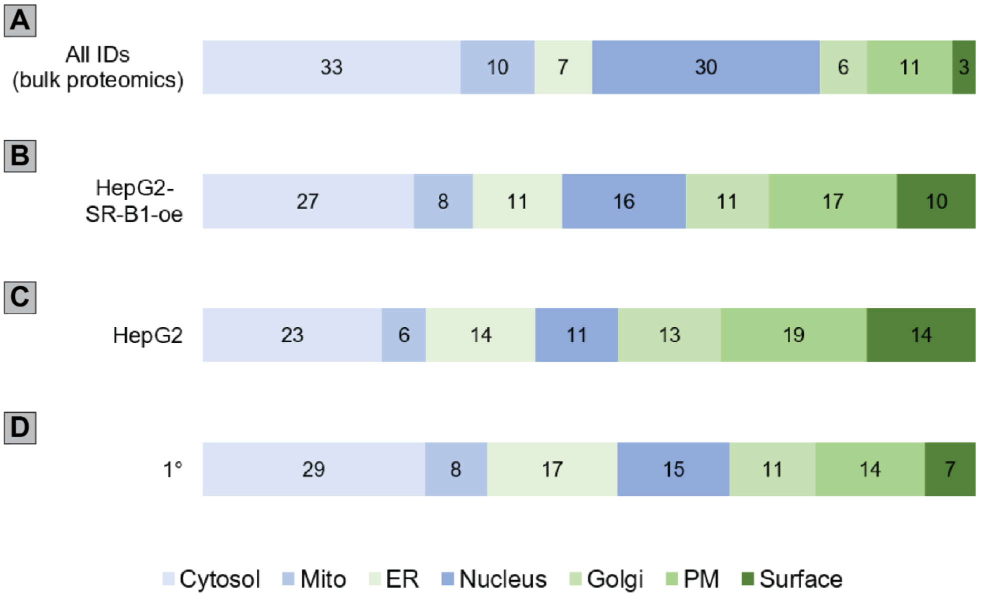


<u>Figure S38</u>: **POCA using HDL-1 identifies proteins from similar compartments across the three different cell lines studied.** Percentage of the proteins detected or enriched in different hepatocyte cell lines according to subcellular location. Cellular compartments in blue correspond

to a decrease in the prevalence between all IDs and an HDL-POCA experiment (cytosol, mitochondrial, and nuclear proteins), while cellular compartments in green correspond to an increase in the prevalence (endoplasmic reticulum, golgi, PM, and surface proteins). **(A)** Distribution of proteins detected in the bulk proteomics/expression analysis of both HepG2 and HepG2-SR-B1-overexpressing cells. **(B)** Distribution of proteins enriched by HDL-POCA in the HepG2-SR-B1-overexpressing cells. **(C)** Distribution of proteins enriched by HDL-POCA in the HepG2 cells. **(D)** Distribution of proteins enriched by HDL-POCA in the murine primary hepatocytes. All MS data can be found in **Data S4**.

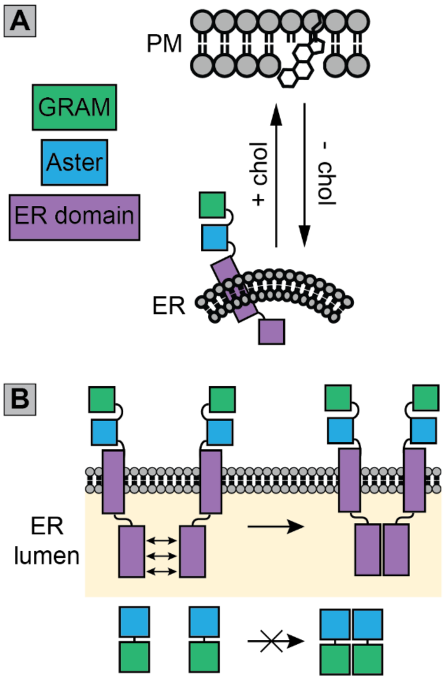


__Figure S39__: **Aster proteins translocate to ER-PM membrane contact sites upon cholesterol loading and oligomerizes via interactions between ER domains. (A)** Schematic diagram showing the translocation of an Aster protein to an ER-PM contact site upon loading with cholesterol, and the localization away from the PM in cholesterol-depleted conditions. **(B)** Schematic diagram showing the oligomerization of Aster within the ER membrane, mediated by interactions between lumenal portions of the ER domains of Aster proteins[18].

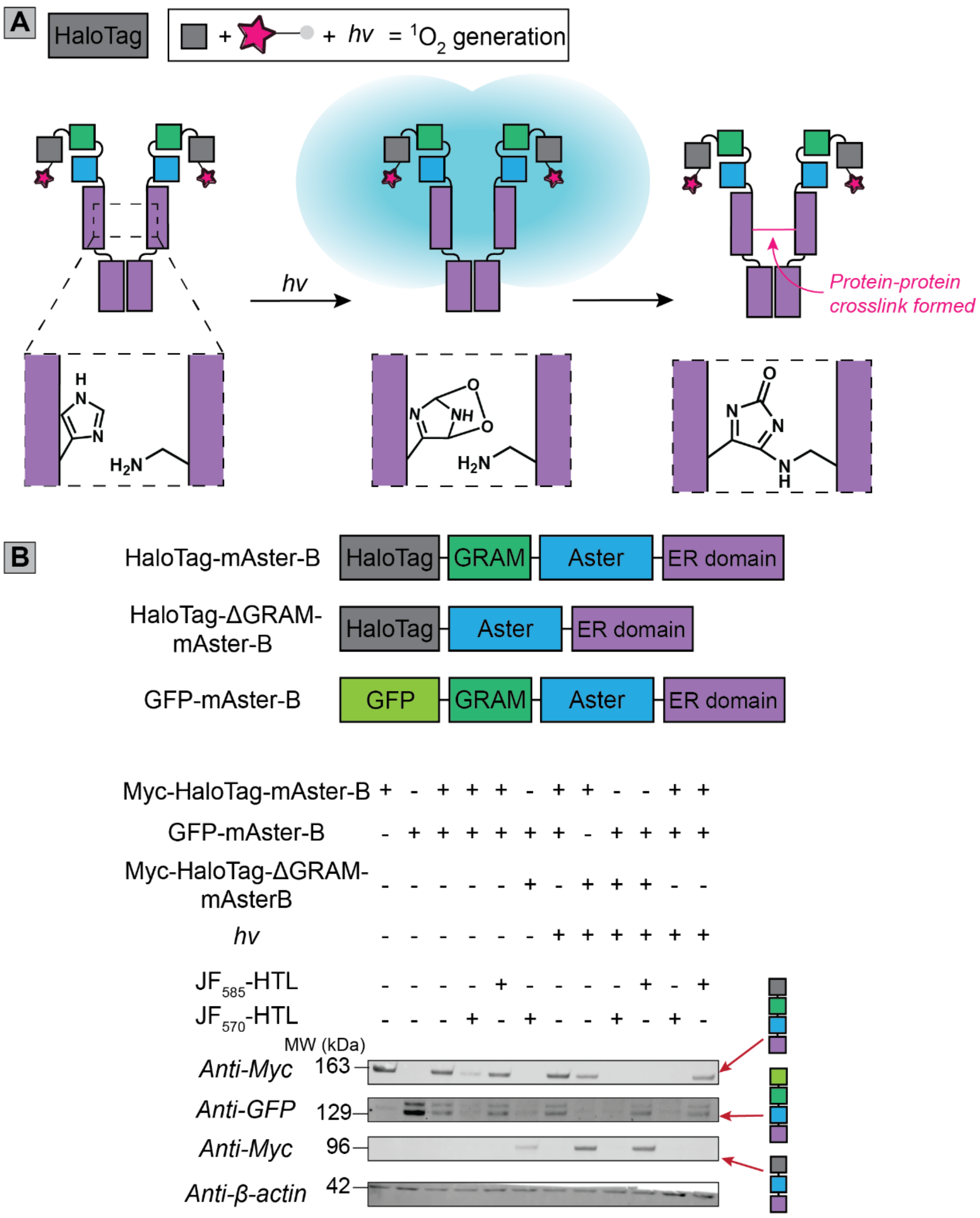


**<u>Figure S40</u>: Crosslinking of Aster constructs requires singlet oxygen and is not affected by the removal of the N-terminal GRAM domain. (A)** Schematic diagram of the different Aster constructs used and immunoblots of protein abundance. HEK293T cells transiently coexpressing different combinations of Aster fusion proteins had singlet oxygen induced by incubation with $JF_{570}$-HTL and irradiation or were treated with control conditions that do not produce large amounts of singlet oxygen: either no irradiation, no $JF_{570}$-HTL or irradiation after incubation with $JF_{585}$-HTL which is not a competent photosensitizer. **(B)** Schematic diagram showing the formation of protein-protein crosslinks upon singlet oxygen generation in the proximity of oligomerized Aster proteins. A histidine-lysine crosslink is shown in the ER domain as an example: the exact nature of the crosslinks formed and their location(s) were not deduced.

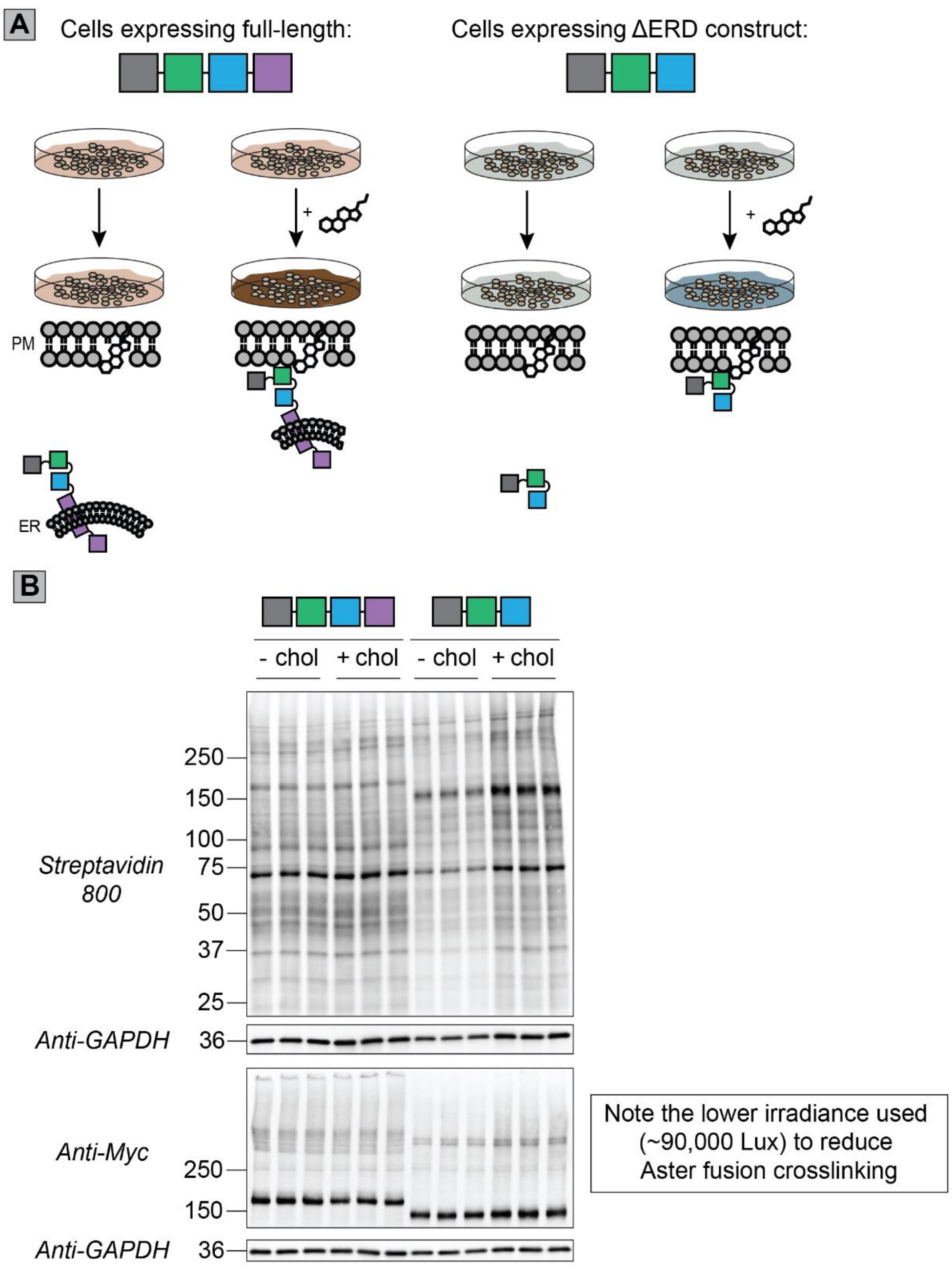


**Figure S41**: **Schematic depiction of the strategy for investigating the cholesterol-dependent interactome of Aster constructs using Halo-POCA**. **(A)** Schematic workflow for investigating the cholesterol-dependent interactome of Aster-B using Halo-POCA. Cells expressing either the full-length HaloTag-mAster-B or HaloTag-mAster-B-ΔERD will be subjected to Halo-POCA labeling in either cholesterol-starved or cholesterol-loaded conditions. **(B)** Visualization of protein labeling by streptavidin blot and fusion proteins for the Halo-POCA experiment with different HaloTag-Aster constructs in cholesterol-depleted or cholesterol-loaded HEK293T cells. HEK293T cells transiently expressing either HaloTag-mAster-B or HaloTag-mAster-B-ΔERD were starved overnight of cholesterol and then treated with $JF_{570}$-HTL (100 nM, 10 min) before they were washed and either loaded with mβCD-cholesterol (100 µM) in 1%

LPDS/DMEM or left untreated. After the media washout, cells were treated with propargylamine (10 mM) and irradiation (15 W yellow LED, 90,000 Lux max intensity, 5 min, on ice), lysis, click conjugation to biotin-azide, and a portion of the samples were set aside for separation by SDS-PAGE and subsequent immunoblotting. Experiments were conducted in triplicate in HEK293T cells.

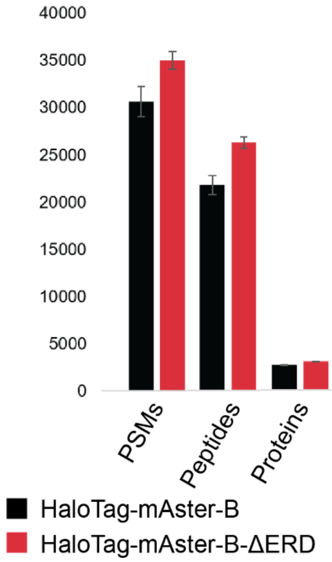


**<u>Figure S42:</u> POCA with two different HaloTag-Aster constructs labels with similar efficiency.** Counts of identified PSMs, peptides, and proteins identified by proteomics at 1% FDR for the Halo-POCA MS experiments using two different HaloTag-Aster fusions. Height of the bars represents the average of measurements and error bars represent the standard deviation of each measure. HEK293T cells transiently expressing either HaloTag-mAster-B or HaloTag-mAster-B-ΔERD were starved overnight of cholesterol and then treated with $JF_{570}$-HTL (100 nM, 10 min) before they were washed and either loaded with mβCD-cholesterol (100 µM) in 1% LPDS/DMEM or left untreated. After the media washout cells were treated with propargylamine (10 mM) and irradiation (15 W yellow LED, 90,000 Lux max intensity, 5 min, on ice), lysis, click conjugation to biotin-azide, enrichment and MS sample preparation procedures, then the tryptic peptides were analyzed by LC-MS/MS and searched in MSFragger[7]. Experiments were conducted in triplicate in HEK293T cells. All MS data can be found in **Data S5**.

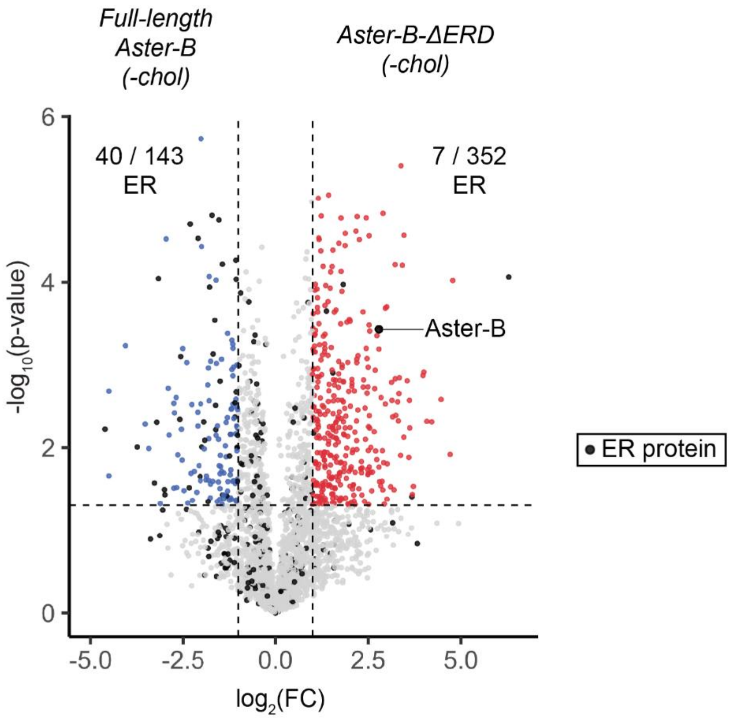


**<u>Figure S43:</u> Halo-POCA with full-length Aster-B enriches many more ER proteins compared to the ΔERD construct.** Volcano plot showing the differential protein labeling between HEK293T cells transiently expressing full-length HaloTag-mAster-B (left side, blue data points) and HaloTag-mAster-B-ΔERD (right side, red data points) in cholesterol-depleted conditions. ER-annotated proteins are represented by black data points. HEK293T cells transiently expressing HaloTag-mAster-B were starved overnight of cholesterol and then treated with $JF_{570}$-HTL (100 nM, 10 min) before they were washed and either loaded with mβCD-cholesterol (100 µM) in 1% LPDS/DMEM or left untreated. After the media washout cells were treated with propargylamine (10 mM) and irradiation (15 W yellow LED, 90,000 Lux max intensity, 5 min, on ice), lysis, click conjugation to biotin-azide, enrichment and MS sample preparation procedures, then the tryptic peptides were analyzed by LC-MS/MS and searched in MSFragger[7]. For measures of statistical significance, variances were calculated for each sample-condition pairing and a corresponding two-tailed t-test was performed to generate *p*-values. Enrichment criteria were set as raw *p*-value < 0.05 and $\log_2(FC) > 1$. Experiments were conducted in triplicate in HEK293T cells. All MS data can be found in **Data S5**.

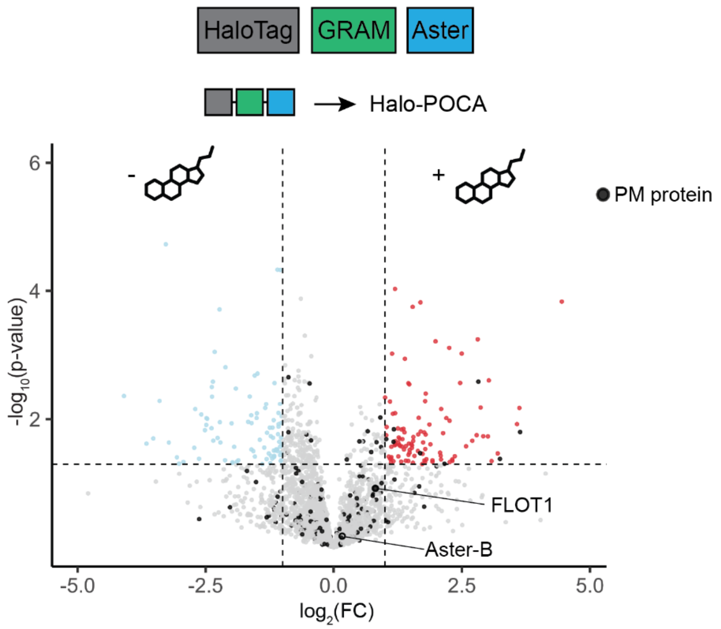


**<u>Figure S44:</u> Halo-POCA with full-length Aster-B enriches many more ER proteins compared to the ΔERD construct.** Volcano plot showing the differential protein labeling between HEK293T cells transiently expressing HaloTag-mAster-B-ΔERD when starved of cholesterol (left side, blue data points) and loaded with cholesterol (right side, red data points). PM-annotated proteins are represented by black data points. HEK293T cells transiently expressing HaloTag-mAster-B-ΔERD were starved overnight of cholesterol and then treated with $JF_{570}$-HTL (100 nM, 10 min) before they were washed and either loaded with mβCD-cholesterol (100 µM) in 1% LPDS/DMEM or left untreated. After the media washout cells were treated with propargylamine (10 mM) and irradiation (15 W yellow LED, 90,000 Lux max intensity, 5 min, on ice), lysis, click conjugation to biotin-azide, enrichment and MS sample preparation procedures, then the tryptic peptides were analyzed by LC-MS/MS and searched in MSFragger[7]. For measures of statistical significance, variances were calculated for each sample-condition pairing and a corresponding two-tailed t-test was performed to generate *p*-values. Enrichment criteria were set as raw *p*-value < 0.05 and $\log_2$(FC) > 1. Experiments were conducted in triplicate in HEK293T cells. All MS data can be found in **Data S5**.

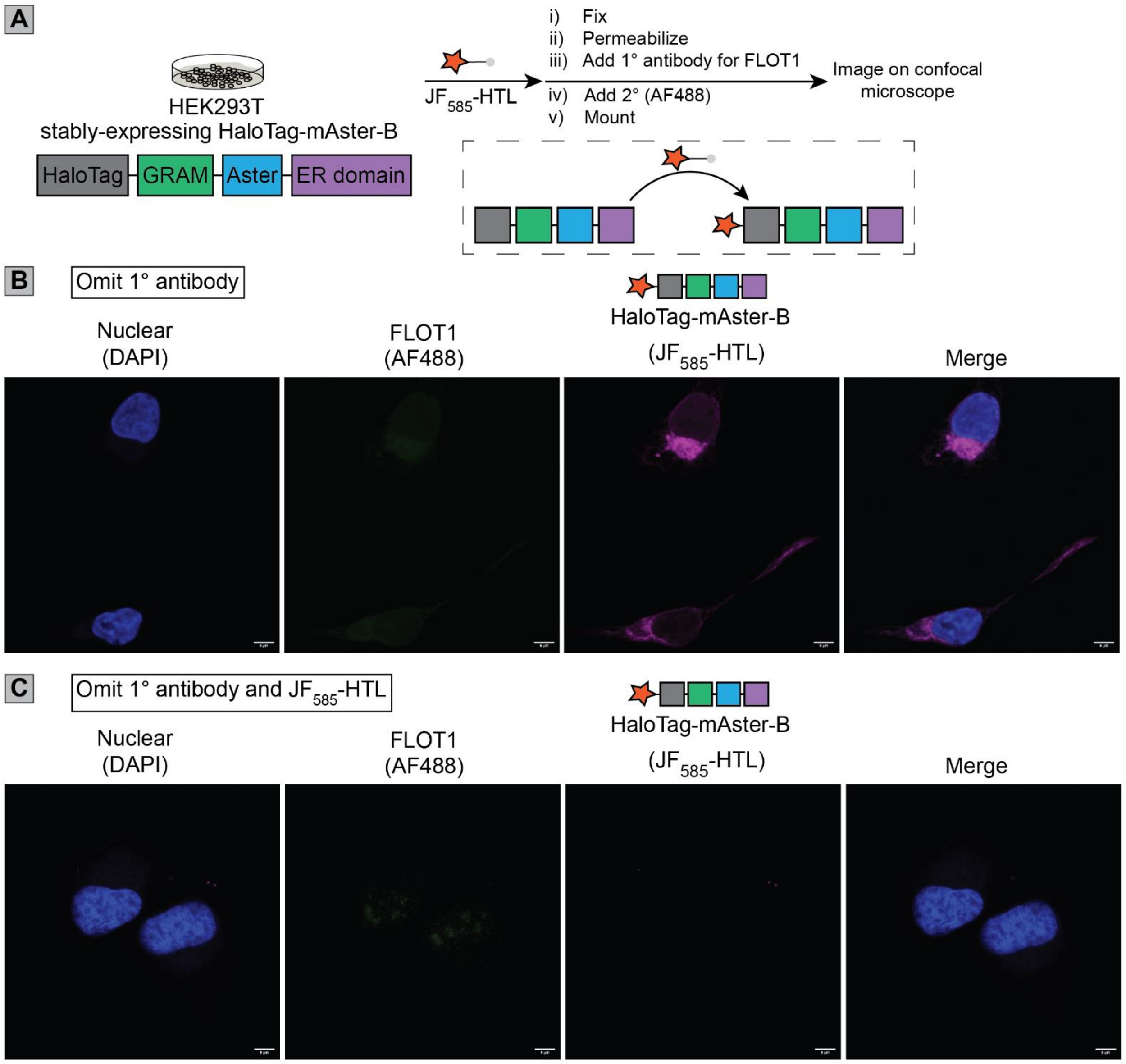


**<u>Figure S45:</u> Control images for FLOT1 immunofluorescence analysis. (A)** Schematic workflow for immunofluorescence of FLOT1 in cells stably expressing HaloTag-mAster-B. **(B)** Confocal microscopy images of cells that were not treated with anti-FLOT1 primary antibody. **(C)** Confocal microscopy images of cells that were not treated with anti-FLOT1 primary antibody nor $JF_{585}$-HaloTag ligand. HeLa cells stably expressing HaloTag-mAster-B were grown overnight, washed twice with HBSS, then treated with $JF_{585}$-HaloTag ligand (100 nM, 10 min) or vehicle for part 'C'. Cells were washed, fixed, permeabilized, washed, and incubated with blocking buffer overnight. After washing, cells were incubated with a 488-conjugated secondary antibody, washed, stained with DAPI, mounted on slides, and imaged the next day. Each image is a 1 µm-thick slice. Scale bars = 10 µm.

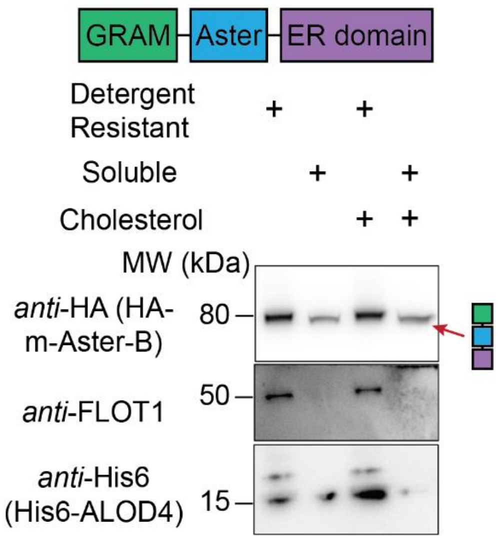


**<u>Figure S46:</u> Full length Aster-B is primarily found in detergent-resistant membranes (DRMs).** Experiment is analogous to that shown in **Figure 5G** of the main text but uses HA-mAster-B instead of HA-GRAM-mAster-B. HA-mAster-B preferentially associates with the DRM fraction. HEK293T cells transiently expressing HA-Aster-B were treated or not treated with methy-β-cyclodextrin-cholesterol complexes (100 µM, 1 h). Cells were washed and then incubated with His-tagged ALOD4[19] (20 µg/ml in DPBS) for 15 minutes at 37 °C. Cells were washed and lysed into TNE buffer by passing through a 23 G needle and nuclei were pelleted at 1000 g for 5 mins at 4 °C. Triton X-100 was added to the supernatant at a final concentration of 1%, gently mixed, and left on ice for 30 mins. The samples were adjusted to a concentration of 40% iodixanol, then sequentially overlaid with 40% iodixanol and TNE buffer. After ultracentrifugation, the top 50% of volume was removed and designated the detergent resistant fraction and the bottom 50% was designated the detergent soluble fraction and protein amounts in each fraction were assessed by immunoblot.

**General Methods**.

*Cloning and generation of HaloTag constructs*

List of plasmids with detailed information used in this study can be found in Table S2. PCR fragments were amplified using Phusion polymerase (Berkeley). The vector and PCR product(s) were digested using standard enzymatic restriction (37 °C, 2–3 h) in 10% rCutSmart buffer. Linearized DNA fragments were cleaned up after separation on a 1% agarose/TAE gel by excising the band and isolating the DNA (Zymo). Fragments were then joined by either Gibson Assembly (New England Biolabs) or by double-digestion with restriction enzymes, gel isolation of the cut fragments, and T4 ligation (New England Biolabs). Ligation, assembly, and gateway cloning products were transformed into TOP10 or Stbl3 competent cells and sent for sequencing (Azenta/Genewiz) to confirm the cloning using a standard primer (i.e. T7, M13F) or a custom primer (Primer 3, Table S1).

For plasmids, we thank Michael Huen for pMH-HaloTag (Addgene plasmid # 154144 ; http://n2t.net/addgene:154144 ; RRID:Addgene_154144), Kai Johnsson for pcDNA5/FRT/TO_CalR_HaloTag7_KDEL (Addgene plasmid # 175530 ; http://n2t.net/addgene:175530 ; RRID:Addgene_175530), Michael Lampson for pERB254 (Addgene plasmid # 67762 ; http://n2t.net/addgene:67762 ; RRID:Addgene_67762), and Eric Campeau & Paul Kaufman for pLenti CMV GFP Hygro (656-4) (Addgene plasmid # 17446 ; http://n2t.net/addgene:17446 ; RRID:Addgene_17446).

*Cell lines*

HEK293T (CRL-3216, Homo sapiens, female, embryonic kidney) and HepG2 (HB-8065, Homo sapiens, male, hepatocellular carcinoma) cells were cultured in DMEM supplemented with 10% fetal bovine serum (FBS) and 1% antibiotics (penn/strep, 100 U/mL). HepG2-SR-B1-GFP cells were cultured in DMEM supplemented with 15% FBS, 4 mM glutamine, and 10X amino acid mix. Murine primary hepatocytes were isolated from C57BL/6 mice and cultured according to an existing procedure[20] and used 4 h after isolation. Media was sterile filtered (0.22 μm) prior to use. Cells were free of visual contamination based on daily inspection. Cells were maintained in a humidified incubator at 37°C with 5% CO2.

*Primary hepatocytes isolation*

Primary hepatocytes were isolated from wild-type mice according to an established procedure[20]. The mice were anesthetized with 200 μl ketamine/xylazine solution by intraperitoneal injection before a cannula was inserted into the vena cava. The liver was perfused through the portal vein with 50 ml of solution A (Hanks's solution with 1 mM EGTA, 20 mM HEPES and 1× pen/strep), followed by 50 ml of solution B (William Medium E with 1× GlutaMax, 20 mM HEPES, 1× pen/strep and 50 μg/ml liberase) at a speed of 5 ml/min. The liver was placed in a dish, extraneous tissue was removed and the liver was dismantled using forceps to release hepatocytes into the medium. Using a 25-ml pipette, hepatocytes were filtered through a 70-μm cell strainer into a fresh 50-ml conical tube. Cells were centrifuged at 50 g at 4 °C for 3 min to gently pellet cells. Cells were gently centrifuged an additional two times to wash. Cells were plated in warmed washing/Plating medium (William Medium E medium with 1× GlutaMax, 20 mMHEPES, 1× pen/strep, 1 μM dexamethasone, 100 nM insulin and 5% FBS) at a density of $5 \times 10^5$ cells per well in collagencoated six-well plates. Cells were allowed to settle for 3–4 h in washing/plating medium before experiments were conducted.

*Generation of cell lines stably expressing HaloTag constructs*

For preparation of lentiviruses, HEK293T cells were cultured in antibiotic-free media (DMEM + 10% FBS) to a confluency of 80-90% in 10 cm plates. They were transfected by adding a mixture of 10 ug lentiviral vector p156RRL-sinPPT-CMV-[gene]-PRE/Nhe I, pVSVG (4 ug; Addgene #8454) and Δ8.9 (8 ug; Addgene #8455) and 66 uL of Turbo DNAfectin3000 (Lamda Biotech Inc.) in Opti-MEM (1.00 mL) to the media. The DNAfectin-containing media was replaced with fresh antibiotic-free media and the cells were incubated 24 hours for lentiviral generation. The lentivirus-containing media was collected and used directly to transduce cells at 70-80% confluency (1-2 mL virus-containing media per 6 cm plate) in the presence of polybrene (8 ug/mL) for 24-36 h before the virus-containing media was replaced with 10% FBS/DMEM. After 24 h, the media was swapped for selection media containing hygromycin B (HEK293T: 200 ug/mL, HeLa: 350 ug/mL) and the cells were maintained in selection media for at least 3 passages. The percentage of transduction-positive cells was assayed by seeding 50,000 cells in a 24-well plate, incubating for 16 h, and then replacing the media with 100 nM $JF_{585}$-HTL in HBSS and incubating for 10 min before imaging on an epifluorescence scope.

*Transient transfection*

Adherent cells were transfected at 65-80% confluency 18-24 h after seeding. For a 10-cm plate, plasmid (5.0 ug), serum-free DMEM (350 uL) and polyethylene imine (PEI, 35 uL of 1 mg/mL, 20k average MW) were mixed and incubated for 15 min at room temperature, then added dropwise to the cells. For a 6-cm plate, plasmid (2.0 ug), serum-free DMEM (120 uL), and PEI (16 uL of 1 mg/mL) were used. Cells were incubated with DNA complexes for 24-36 h before they were used further.

*Chol-POCA procedure*

Cholesterol probes were complexed in methyl-*beta*-cyclodextrin, average MW 21,000 (mβCD). Cholesterol probes were complexed in HDL according to the method of Robichaud et al [21], *vide infra* "Cholesterol probe HDL complexation" section.

When HEK293T cells were used, plates were coated with poly-D-lysine (0.1 mg/mL, 10 min at 37 °C, washed 3x with sterile H2O) and dried prior to seeding cells. For gel-based experiments, cells were grown overnight in the corresponding full media, washed once with PBS, then serum- and antibiotic-free DMEM (DMEM -/-) was added to the cells. Cells were incubated (37 °C, 5% CO2) for 4 h to briefly starve the cells of cholesterol and lipids. For MS-based experiments, overnight incubation in 1% LPDS/DMEM was used to starve the cells of cholesterol and lipids. Next, mβCD-complexed probe dissolved in either DMEM -/- (gel-based experiments) or 1% LPDS/DMEM (MS-based experiments) was added to the cells (10 µM unless indicated otherwise, 1 h). Cells were then carefully washed once with PBS, once with ice-cold PBS, and an ice-cold solution of propargylamine (10 mM unless otherwise stated) in PBS was added to the cells. Cells were immediately put on ice and irradiated using a yellow LED (SunLite PAR30, 15 W) at an illuminance of ~170,000 Lux as measured by a lux meter (Dr. meter cat. LX1330B) for 5 min. Cells were then aspirated, washed once with cold PBS, and harvested into cold PBS (1.5 mL/6-cm plate, 5 mL/10-cm plate) with a cell scraper. Cells were collected by centrifugation (400 *g*, 5 min, 4°C), the supernatant was aspirated, and cells were either used immediately or frozen and stored at -80 °C until use.

For the competition experiment shown in **Figures 3A** & **3B**, a solution of mβCD-cholesterol (100 µM) in 1% LDPS/DMEM was added to the cells and incubated for 30 min. A equal volume of 20 µM mβCD-**1** in 1% LDPS/DMEM was then added to the cells (10 µM final concentration) and then incubated for 30 min. Cells were then treated as described above.

*Halo-POCA procedure*

HEK293T cells transiently or stably-expressing a HaloTag construct were grown to 70–80% confluency in plates coated with poly-D-lysine. The media were aspirated and replaced with HBSS containing $JF_{570}$-HaloTag ligand[4] (100 nM) and cells were incubated for 10 min at 37 °C and 5% CO2. Cells were then washed once with HBSS and then full media was added. Cells were incubated for 20 min before the media were removed and the full media wash was repeated once more. Cells were then aspirated before an ice-cold solution of propargylamine (10 mM unless otherwise stated) in PBS was added to the cells. Plates were then irradiated using a yellow LED (SunLite PAR30, 15 W) at an illuminance of ~170,000 Lux for 5 min on ice. Cells were then aspirated, washed once with cold PBS, and harvested into cold PBS (1.5 mL/6-cm plate, 5 mL/10-cm plate) with a cell scraper. Cells were collected by centrifugation (400 *g*, 5 min, 4°C), the supernatant was aspirated, and cells were either used immediately or frozen and stored at -80 °C until use.

For the +/- cholesterol experiments with HaloTag-Aster fusions in **Figure 5E, Figure S39, S40,** and **S41**, the washout procedure was changed slightly to accommodate cholesterol loading. After incubation with $JF_{570}$-HTL, cells were washed once with HBSS and then incubated with 1% LPDS/DMEM supplemented with mβCD-cholesterol (100 µM) for 40 min. Samples in the -cholesterol condition were incubated with non-supplemented 1% LPDS/DMEM for 40 min. Cells were then aspirated, an ice-cold solution of propargylamine (10 mM) in PBS was added, and cells were then irradiated at a maximum illuminance of 90,000 Lux to prevent complete crosslinking of the "full-length" HaloTag-mAster-B fusion. Cells were then treated according to the rest of the Halo-POCA procedure described above.

A picture of a prototypical setup for irradiating cell plates is below:

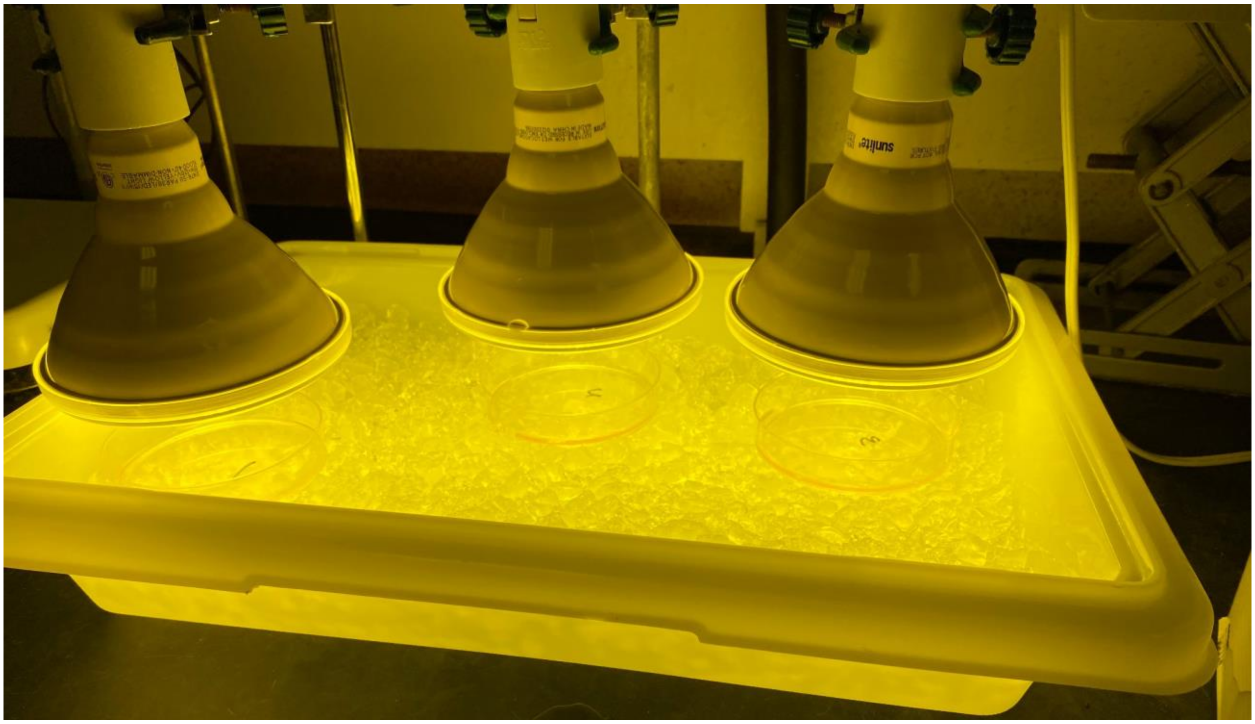

*Cell lysis and click chemistry*

Cells were lysed by the addition of either 0.3% CHAPS/0.5% NP40/PBS (gel-based experiments) and aging >30 min on ice or by the addition of 6 M Urea (MS-based experiments) and 3x freeze-thaw cycles using dry ice. Lysates were then clarified by centrifugation (20,000 *g*, 5 min, 4°C) and protein concentrations were determined using a DC protein assay kit (Bio-Rad, Cat# 5000111) and the lysate diluted to the indicated concentration. Click chemistry was performed by combining, in order, three parts (v/v) 1.7 mM TBTA (in 4:1 tBuOH:DMSO), one part 50 mM CuSO4 (aq.), one part 50 mM TCEP (aq.), and one part azide click partner in DMSO. TAMRA-azide: 200 µM, sulfo-Cy5-azide: 50 µM.

*SDS-PAGE, fluorescent gels, and immunoblots*

Samples were normalized to 1-5 mg/mL and separated on a pre-made tris-glycine gel (Bio-Rad, stain-free). Total protein on the gel was imaged using a Bio-Rad ChemiDoc imager with the stain-free [22] setting. Gels were transferred to a nitrocellulose (Bio-Rad) or PVDF membrane (Bio-Rad) using a semi-dry transfer system (2.5 A, 12V, 7 min, Bio-Rad) or traditional tank transfer (30 V, 16 h, 4 °C) and blocked in 5% (w/v) milk in Tris-buffered saline/0.1% Tween 20 (TBST) or 5% (w/v) BSA in TBST for >1 h at RT (see Table S3 for blocking & botting conditions for each antibody). Membranes were incubated with primary antibodies at the indicated dilution and temperature specified in Table S3, then rinsed 5 times with TBS and washed twice with TBST (2 x 5 min). Membranes were then incubated with secondary antibodies (1/5000 dilution) for 90 min at RT, rinsed 5 times with TBS and washed once with TBST (5 min) prior to imaging. For blots assessing biotin signal, membranes were incubated with a streptavidin-CW800 conjugate overnight at 4°C. Stain-free gel, fluorescent gel (rhodamine, Cy5), Streptavidin-800, and secondary antibody signals were imaged using a Bio-Rad ChemiDoc. Immunoblot signals were quantified in ImageJ (v1.54f) by taking the greyscale pixel density, calculating the inverted value

(255 - greyscale value), and subtracting the background. The inverted, background-corrected values for the actin signal of each replicate were used to normalize the inverted, background-corrected values of the SR-B1 signal for the corresponding replicate.

*mβCD-complexation of cholesterol diazirine probes or cholesterol-JF570 probes **1** & **2***

Diazirine probes were dissolved in either DCM (cholesterol diazirine probes) or 1:1 DCM:MeOH (probes **1** & **2**) at a concentration of 10 mM and then 50 uL was charged to an amber glass 1.5 mL screw-top vial. The solvent was evaporated with a gentle stream of nitrogen gas (~1 min) and then the vial was placed under high vacuum for 5 min to ensure removal of the solvent. 250 uL of 5% (w/v) mβCD as a solution in water was added to the residual probe. The vial was capped and parafilmed, vortexed briefly, then sonicated in a water bath for 30 min before it was incubated overnight (12–16 h) at 37 °C with shaking (250 RPM). The solution was used for experiments at a nominal concentration of 2 mM.

*mβCD-complexation of cholesterol*

To a 50 mL RBF with a magnetic stir bar is added methyl beta cyclodextrin (500 mg) and 10.0 mL of H2O under N2 to afford a 5% (w/v) solution. The solution is heated to 80 °C before 1.50 mL of an ethanolic cholesterol solution (10.0 mg/mL) is added dropwise over 2 min. The faintly turbid mixture is allowed to cool over 1 h to 22 °C before it is concentrated via rotary evaporation and the sticky residual film is dried overnight under high vacuum. The next day, the solids are dissolved in 15.54 mL H2O and the resulting solution is filtered with a 0.22 µm filter. The solution was stored at 4 °C and used for a maximum of 3 months.

*Cholesterol probe HDL complexation*

HDL were isolated by density gradient ultracentrifugation as described by Redgrave et al. [23]. Lipoprotein-deficient serum (LDPS) was treated with the additives thomerosol (3 mM), protease inhibitor cocktail (1 tablet, cOmplete), BHT (0.5% w/v), and trolox (0.1% w/v). Cholesterol probe (150 ug) as a solution in volatile organic solvent (EtOH, MeOH, DCM, or a mixture) was added to a 1.5 mL vial and dried down with a gentle stream of nitrogen. PBS (1 mL) was added, and the suspension was sonicated for 60 minutes at 37 °C. A solution of HDL (1 mg, human) was dissolved in 3 mL PBS and added to the residual cholesterol probe. Next, LDPS (3.0 mL) was added and the mixture was layered with nitrogen then incubated at 37 °C for 20 h. The density was adjusted to 1.215 g/mL with sucrose solution and then centrifuged at 50,000 rpm for 18 h. It was dialyzed three times to a pH 7.42 buffer containing 5 mM Tris, 50 mM NaCl, 5 mM EDTA using a 10k MWCO dialysis cassette. The buffer was replenished after 4 h, after 12 h, and a final time after another 4 h.

*CETSA and related experiments*

HEK293T cells were seeded in 10-cm plates and grown overnight to a confluency of 70–80%. The next morning, cells were treated with mβCD-cholesterol (150 µM in $H_2O$) or 25-OHC (10 µM in EtOH) for 30 min unless otherwise stated. After swapping into PBS, cells were harvested with a cell scraper and pelleted by centrifugation (300 *g*, 5 min,4 °C) before they were resuspended in PBS (300 uL) and the cell suspensions were aliquoted into different 0.2 mL PCR tubes (30 uL aliquots) and placed in a PCR block preheated to a spatial temperature gradient. After 3.0 min, cells were removed and allowed to sit at ambient temperature for 3.0 min before they were snap frozen with liquid nitrogen. After thawing, 5.0 uL of a solution of non-ionic surfactant ("NP-40 substitute", polyethylene glycol nonylphenyl ether, CAS 9016-45-9) was added to give the desired concentration. Samples were then snap frozen and thawed three more times before the lysates were transferred to 1.5 mL microcentrifuge tubes, whereupon precipitated material was pelleted by centrifugation (20,000 *g*, 45 min, 4 °C). The supernatants were transferred to fresh tubes and the protein concentration was assayed. Samples were normalized to the lowest concentration at each temperature and then separated by SDS-PAGE.

*Proteomic sample preparation: protein-level enrichment*

Alkynylated lysate (200 uL of 1.0 mg/mL) had biotin appended by CuAAC by adding 24 uL of a freshly-prepared click mix consisting of TBTA (4 volumes of 1.7 mM stock in DMSO/t-butanol 1:4, final concentration = 100 μM), CuSO4 (1 volume of 50 mM stock in water, final concentration = 1 mM), TCEP (1 volume of fresh 50 mM stock in water, final concentration = 1 mM), and biotin-C3-azide (1 volume of of 200 mM stock in DMSO, final concentration = 4 mM) and incubating for 1h at RT in the dark. SDS was added (26 uL of 10% solution in PBS, 1.2% final concentration) and the samples were briefly vortexed before they were taken through SP3 cleanup[14]. 20 μL Sera-Mag SpeedBeads Carboxyl Magnetic Beads, hydrophobic (50 μg/μL, total 1 mg) and 20 μL Sera-Mag SpeedBeads Carboxyl Magnetic Beads, hydrophilic (50 μg/μL, total 1 mg) were mixed and washed with water three times. The bead slurries were then transferred to the biotinylated samples, incubated for 5 min at RT with shaking (1000 rpm) and washed three times with 1 mL 80% EtOH/20% $H_2O$ on a magnetic rack. Proteins were eluted from the SP3 beads with 100 μL of 0.2% SDS in PBS for 30 min at 37°C with shaking (1000 rpm). The elution was repeated once and combined with the initial eluent. Streptavidin-agarose beads (50 uL suspension per sample) were washed twice with 0.2% SDS/PBS, resuspended in 0.2% SDS/PBS (500 uL per sample) and 500 uL of the streptavidin-agarose suspension was added to each sample. Sample concentrations were determined by DC assay and an equal protein mass from each sample were added to 500 uL of the streptavidin bead suspension then rotated at 22 °C for 2 h. Beads were pelleted by centrifugation (4,000 g, 1 min), the supernatant was removed, and the beads were washed successively with 1 mL of 6 M urea/0.2% SDS/PBS, 0.2% SDS/PBS, 3 times with PBS, and 3 times with water. The beads were then resuspended in 200 uL 6 M urea/PBS and cysteines reduced with DTT (10.5 uL of a 200 mM solution, final concentration 10 mM) for 30 min at 37 °C and then alkylated with iodoacetamide (11.2 uL of a 400 mM solution, final concentration 20 mM) for 60 min at 22 °C. The suspension was diluted with 400 uL PBS (to achieve a concentration of approximately 2 M urea), the beads were spun down and supernatant removed. The beads were resuspended in 2 M urea/8 mM $CaCl_2$/PBS and 30 ng of trypsin (TPCK treated) was added and the digestion was allowed to proceed overnight at 37 °C with shaking (250 rpm). The peptide-containing supernatant was collected and then desalted with a plug of C18-silica according to the manufacturer's protocol. The desalted peptide solutions were dried with a centrifugal vacuum apparatus then reconstituted with 5% acetonitrile and 1% FA in MB water prior to analysis by LC-MS/MS.

*Liquid chromatography-tandem mass spectrometry (LC-MS/MS) analysis*

Peptide samples were analyzed by liquid chromatography tandem mass spectrometry using a Thermo Scientific™ Orbitrap Eclipse™ Tribrid™ mass spectrometer. Peptides were fractionated online using an 18 cm long, 100 µM inner diameter (ID) fused silica capillary packed in-house with bulk C18 reversed phase resin (particle size, 1.9 µm; pore size, 100 Å; Dr. Maisch GmbH). The 74 min water-acetonitrile gradient was delivered using a Thermo Scientific™ EASY-nLC™ 1200 system at different flow rates (solvent A: water with 3% DMSO and 0.1% formic acid and solvent B: 80% acetonitrile with 3% DMSO and 0.1% formic acid). Gradient parameters are detailed in Table S5. Data were collected with charge exclusion (1, 8,>8). Data were acquired using a Data-Dependent Acquisition (DDA) method consisting of a full MS1 scan (Resolution = 120,000) followed by sequential MS2 scans (Resolution = 15,000) to utilize the remainder of the 3 second cycle time. Precursor isolation window was set as 1.6 and normalized collision energy was set as 30%.

*Protein identification and quantification*

Raw data collected by LC-MS/MS were searched with MSFragger (v3.5 or 3.8) and FragPipe (v18.0 or 20.0) with the UniProt reviewed (*Homo sapiens* downloaded 01 January 2020, *Mus musculus* downloaded 22 May 2023) *Homo sapiens* database (downloaded 01 January 2020). For the murine primary hepatocyte data in Figure 4, the *Mus musculus* database (downloaded 22 May 2023) was used and mouse genes were mapped to their human orthologs (*vide infra*). Precursor and fragment mass tolerance was set as 20 ppm. Fully tryptic (KR) search space with up to 2 missed cleavages was used. Peptide length was set 7 - 50 and peptide mass range was set 500 - 5000. Variable modification mass shift and max occurrences per peptide were as follows: N-terminal acetylation 42.0106 (1), methionine oxidation 15.9949 (3), histidine oxidation 15.9949 (1), histidine singlet oxygen and capture by water 31.9898 (2), cysteine carbamidomethylation 57.0215 (3). Peptide spectrum match validation was performed using Percolator[ref] to a minimum probability of 50% and MSBooster[9] and protein validation was performed using ProteinProphet[8]to an FDR of 1%. The MS1 intensity ratio of protein abundances using LFQ-MBR were determined with Ionquant[6] to an ion FDR of 1%. For feature detection and peak tracing, a minimum of 3 scans and two isotopes, 10 ppm mass tolerance, and a 0.4 min retention time tolerance were used. For MBR settings, a tolerance of 1 min, 0 minimum correlation, 1% peptide and 1% protein FDR, and 100,000 top runs were used. Perseus v2.0.7.0 was used for filtering and missing value imputation of the search results. The “combined_protein.tsv” output file from each LFQ-MBR search was opened in Perseus and the protein intensities were imported. The columns comprising the two groups (generally, control and treatment, i.e. “control” would be -light and “treatment” would be +light) were annotated, zero (0) values were converted to NaN, and rows were filtered based on valid values (i.e. non-NaN values are valid) with the criterion of at least 2 valid values in at least one group (‘group’ referring to either ‘control’ or ‘treatment’) when 3 replicates were used. For the competition experiment using 4 replicates, the criterion of 3 valid values was used. For experiments using 6 replicates, the criterion of 4 valid values was used. The filtered set of protein intensities were then transformed to the logarithm base 2 of their values. The remaining missing values were imputed separately for each column under the assumption they were normally distributed using the default values of 0.3 width and a downshift of 1.8 standard deviations. The resulting data matrix was exported as a .csv and custom R scripts were used to compile the intensities of proteins and perform statistical analysis. An example R script with relevant annotations to guide data processing is available in **Data File S1**. UniProt protein IDs were used to establish unique protein identities for the purposes of filtering, comparisons, and subcellular annotation lookups.

*Database construction*

Subcellular annotations for proteins were generated as described in Yan et al.[24], which aggregated protein localization information from the Human Protein Atlas[25] (v21.1), UniprotKB[26] (August 2022 release), and CellWhere[27] (accessed August 2022). Cell surface proteins were retrieved from the Surfaceome prepared by Wollscheid and coworkers [28]. Unique proteins were established using UniProt alphanumeric protein identifiers (referred to as UniProt accession numbers). CellWhere localization, HPA main location and UniProt subcellular location columns were mined for specific location keywords (ex. 'golgi'). Proteins containing these keywords are reported in **Data S1**. Surface proteins were retrieved from the SURFY surfaceome database as reported by the Wollscheid lab[28] (accessed 24 October 2023). In **Figure 2**, Sterol-binding proteins were annotated based on a list of proteins (62 ct.) containing the GO terms "cholesterol-binding"(GO:0015485), "sterol-binding" (GO:0032934), and "oxysterol-binding" (GO:0008142). Cholesterol-related proteins were based on a list of proteins (462 ct.) containing 180 GO biological process/molecular function terms. Specific terms used and the complete lists of proteins are available in **Data S2**.

*Mouse-human homolog mapping*

For the data generated from murine (C57BL/6) primary hepatocytes shown in Figure 4, identified mouse proteins were mapped to their human homologs using homology data provided by the Alliance of Genome Resources (retrieved 02 May 2023, v5.4, https://www.alliancegenome.org/downloads#orthology) that was curated by the Jackson Laboratory for mouse and human homology data (retrieved 02 May 2023, http://www.informatics.jax.org/downloads/reports/HOM_MouseHumanSequence.rpt). A custom R script (Supplemental Data File S2) matched the UniProt accession number to the Entrez gene ID for each mouse gene, retrieved the matching 'DB Class Key' value for the human homolog, and returned the corresponding UniProt accession number and Gene identifier. Lookup table and results are available in Data **S4**.

*Protein labeling for recombinant site of labeling analysis*

BSA (5 mg/mL) dissolved in PBS containing 1 mM propargylamine and 6-carboxy-JF$_{570}$ (10 µM) was irradiated with a yellow LED at 170,000 Lux for 5 min at room temperature. Samples had either biotin-C3-N3 or biotin-C4-N3 appended via CuAAC for 1 h and were taken through the sp3-cleanup protocol described above. After overnight trypsin digestion, biotinylated peptides were enriched by the addition of neutravidin-agarose resin, which was washed twice with PBS and twice with H2O. Bound peptides were eluted by three successive additions and collections of 80/20/0.1 MeCN/H2O/formic acid; first for 10 min at room temperature, next for 10 min at 72 °C, and finally for 2 min at room temperature. After concentration and redissolution in MS sample buffer, peptides were analyzed via LC-MS/MS as described above and then searched for biotinylated modifications using. The data were searched with the same parameters as described above, with differences in the variable mass modifications and were not subjected to LFQ-MBR quantification and missing value imputation. Variable modification mass shift and max occurrences per peptide were as follows: N-terminal acetylation 42.0106 (1), methionine oxidation 15.9949 (3), histidine oxidation 15.9949 (1), histidine singlet oxygen and capture by water 31.9898 (2), histidine POCA labeling and click with biotin-C3-N3 395.1740 (2), histidine POCA labeling and click with biotin-C4-N3 409.1897 (2). Cysteine carbamidomethylation 57.0215 (3) was used as a fixed modification. Modified peptide spectra were scored using PTMProphet[15], filtered to score >0.99, and visualized with PDV viewer[16].

*Statistics*

For bar plots, average of replicates was reported as indicated. Displayed error bars represent the standard deviation of each measure. Statistical significance was calculated with unpaired Student's t-tests with unequal variances using Graphpad PRISM (9.4.1). $* p<0.05$, $** p<0.01$, $*** p<0.005$, $**** p<0.0001$, NS $p>0.05$. For all volcano plots, variances were calculated for each sample-condition pairing and a corresponding two-tailed t-test was performed using R stats (4.2.2) to generate *p*-values.

*Microscopy and Immunofluorescence*

Primary human aortic endothelial cells (HAECs) (Cell Applications S304-05a) were used from P4-P7. For plating of cells on 8-well chamber slides for live-cell imaging, 0.1% gelatin coating was first applied, and EGM-2 medium supplemented with 10% FBS was used. Cells were cultured in a 37°C incubator with 5% $CO_2$. To capture mβCD-**1** probe uptake, after 20 min incubation with NucBlue nuclear probe, mβCD-**1** (2 μM) in media was added to confluent HAEC monolayer, and live cell imaging was conducted. Fluorescence images were acquired using Zeiss Observer Z1 with Colibri 7 LED light source, CMOS camera (Photometrics Prime 95B), 20X objective and ZEN Blue software 3.5 (Zeiss). Images were acquired every 60 sec for a total of 20 min. To observe cellular localization of the probe in cholesterol depleted cells, HAECs were first treated with mβCD (2.5 mM) for 15 min to deplete cellular cholesterol, washed, and then treated with mβCD-1 (2 μM) in media for 5 min or 60 min. At the designated time point the cells were washed and live cell images acquired with a 40X oil-immersion objective.

HEK293T or HeLa cells (50,000) were seeded onto sterile, poly-D-lysine coated coverslips within the wells of a 24-well plate and grown for 16 h before further steps were performed. After respective treatments, cells were washed with PBS (2 x 1 min) and then fixed by incubation with formaldehyde (3.7% v/v in PBS) for 15 min at 22 °C. Cells were then washed with PBS (2 x 5 s) and permeabilized by treatment with triton-X 100 (0.1% v/v in PBS) for 6.0 min at 22 °C and then washed with PBS (3 x 1 min). Cells were taken through either immunofluorescence (IF) or visualization of alkynylated proteins with click chemistry workflows. For IF, cells were incubated with blocking solution (2% w/v BSA in PBS supplemented with 0.1% v/v Tween-20 (PBST)) for 1 h at 22 °C before treating with primary antibody solution (1:100 dilution in 2% BSA/PBST) overnight at 4 °C. After washing with PBS (3 x 15 min) at 22 °C, fixed cells were incubated with secondary antibody solution (1:1000 dilution in 2% BSA/PBST) for 1 h. After washing with PBST (3 x 5 min) cells were incubated with DAPI solution (1 ug/mL in PBS) for 5 min at 22 °C, washed with PBS (3 x 1 min), then coverslips were mounted on glass slides using Aqua-Poly/Mount mounting media (Polysciences, Inc.) and kept in the dark for 16–24 h before imaging.

For click chemistry visualization of alkynylated proteins, cells were incubated with 300 uL click mixture for 1 h in the dark at 22 °C. The click mixture was prepared by mixing the following solutions: 150 uL TBTA (100 mM in DMSO, 760 µM final), 200 uL CuSO4 (50 mM in H2O, 500 µM final), 710 uL TCEP (50 mM in H2O, 1.77 mM final), and fluorophore-azide (either TAMRA-azide or sulfonylated Cy5-azide, 1.25 mM in DMSO, 15 µM final). Clicked cells were then washed (2 x 5 min) with washing buffer (0.1% tween-20, 0.5% triton-X-100 in PBS) and stained with DAPI (1 ug/mL in PBS) for 5 min at 22 °C, washed with PBS (3 x 1 min), then coverslips were mounted on glass slides using Aqua-Poly/Mount mounting media (Polysciences, Inc.) and kept in the dark for 16–24 h before imaging.

*Detergent-resistant membrane (DRM) fractionation*

Cells were fractionated into detergent resistant and detergent soluble domains as described previously[29] to assess Aster-B and FLOT-1 dynamics after cholesterol loading. Briefly, human embryonic kidney 293T cells were transiently transfected with HA-Aster-B or HA-GRAM-Aster-B using FuGENE 6 in OptiMEM. Twenty-four hours later cells were treated with or without 100 µM mβCD-cholesterol complexes for 1 h. After incubation with cholesterol, cells were washed twice with DPBS containing calcium and magnesium before being incubated with His-tagged ALOD4 (20 µg/ml in DPBS) for 15 minutes at 37 degrees Celsius. ALOD4 is a peptide that binds to 'accessible' cholesterol in the plasma membrane of cells[19]. Cells were then washed twice with DPBS containing calcium and magnesium and once with TNE buffer (50 mM Tris–HCl, pH 7.4, 150 mM NaCl, 2 mM EDTA,). Cells were lysed in TNE buffer by passing through a 23 G needle and nuclei were pelleted at 1000 g for 5 mins at 4 degrees. Triton X-100 was added to the supernatant at a final concentration of 1%, gently mixed, and left on ice for 30 mins. The samples were adjusted to a concentration of 40% iodixanol by mixing with cold Optiprep and added to the bottom of ultracentrifuge tubes. The samples were sequentially overlaid with 40% iodixanol followed by TNE buffer before being centrifuged at 259, 000 g for 2 h. The top 50% was designated the detergent resistant fraction and the bottom 50% was designed the detergent soluble fraction. Samples were subjected to immunoblotting to quantify Aster proteins (HA primary), ALOD4 ($His_{6X}$ primary), and FLOT1 (FLOT1 primary).

*Cell viability assessment*

HEK293T cells were seeded into the wells (10,000 cells/well) of a 96-well flat-bottomed, white-walled plate and grown overnight. The next day, cells were treated in triplicate with propargylamine (or DMSO vehicle) by adding a 2X solution to achieve the indicated final concentration at a constant DMSO amount of 0.4% v/v. Treatments were performed in descending order, and then CellTiter-Glo® solution was added (reagent is 2X). Luminescence was immediately read by a plate reader, and counts were normalized to the DMSO control.

*General Synthetic Methods*

All reactions were performed in oven dried glassware and kept under a positive pressure of Ar or $N_2$ unless stated otherwise. Silica gel P60 (SiliCycle) was used for column chromatography, SiliCycle 60Å F-254 silica (precoated glass-backed sheets, 0.25 mm thickness) was used for analytical thin layer chromatography, and SilliCycle 60Å F-254 (precoated glass-backed sheets, 1.0 mm thickness, 20 x 20 cm) was used for preparative thin layer chromatography. Plates were visualized by fluorescence quenching under UV light or by staining with iodine, $KMnO_4$, or ninhydrin. Reagents were purchased from Sigma-Aldrich (St. Louis, MO), Alfa Aesar (Ward Hill, MA), EMD Millipore (Billerica, MA), Fisher Scientific (Hampton, NH), Cambridge Isotopes Laboratories (Tewksbury, MA), Enamine (Monmouth Junction, NJ) Oakwood Chemical (West Columbia, SC), Combi-blocks (San Diego, CA), or aablocks (San Diego, CA) and used without further purification unless otherwise noted. Concentration *in vacuo* refers to removal of solvent on a Heidolph rotary evaporator under reduced pressure.

*General Analytical Methods*

$^{1}$H NMR spectra were collected on a Bruker AV400 (400 MHz), AV500 (500 MHz), or NEO600 (600 MHz) spectrometer in the stated solvents as a reference for internal deuterium lock. $^{13}$C NMR spectra were collected on a Bruker AV400 (101 MHz), AV500 (126 MHz), or NEO600 (151 MHz) spectrometer in the stated solvents as a reference for internal deuterium lock. NMR instruments were provided by the UCLA Molecular Instrumentation Center (MIC). All chemical shifts are reported as δ in the standard notation of parts per million (ppm) using the peak of residual proton signals of the deuterated solvent as an internal reference. Coupling constant (*J*) units are in Hertz (Hz) to the nearest 0.1 Hz. Splitting patterns are indicated as follows: br, broad; s, singlet; d, doublet; t, triplet; q, quartet; m, multiplet; or combinations thereof. Low-resolution mass spectrometry was performed on an Agilent Technologies InfinityLab LC/MSD single quadrupole LC/MS (ESI source). High-resolution mass spectrometry was performed on a Waters LCT Premier coupled with an ACQUITY LC and autosampler (ESI source) provided by the UCLA MIC.

6-carboxy-$Ac_2$-DBF and $Ac_2$-DBF-HTL was synthesized according to a previously published procedure[3]. 4',5'-dibromofluorescein was synthesized according to a previously published procedure [30]. 6-carboxy-$JF_{570}$ was synthesized according to a previously published procedure[4].

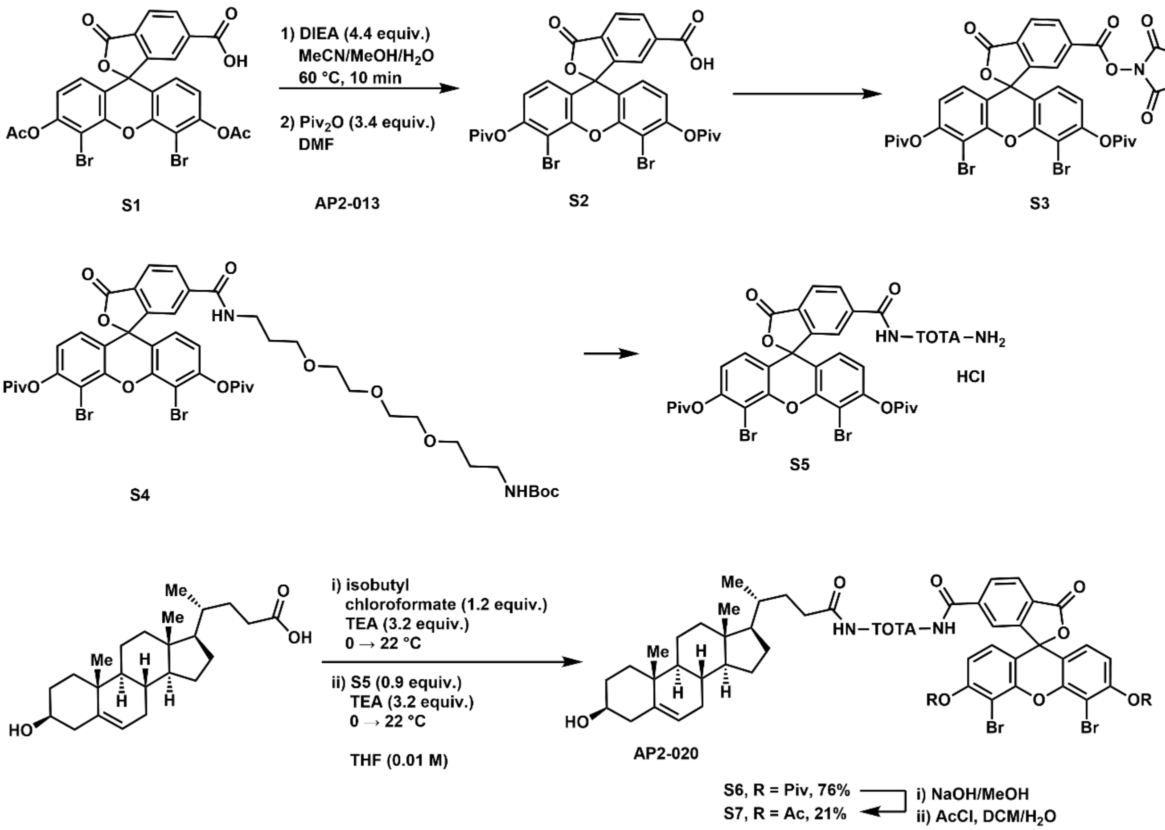


**(4',5'-dibromo-6-(((2,5-dioxopyrrolidin-1-yl)oxy)carbonyl)-3-oxo-3*H*-spiro[isobenzofuran-1,9'-xanthene]-3',6'-diyl bis(2,2-dimethylpropanoate) (S3)**. To a 50 mL round-bottomed flask equipped with a magnetic stir bar was added **S1** (812 mg, 1.31 mmol, 1.00 equiv.) as a pink solid, MeCN (10 mL), water (1 mL), MeOH (2 mL), and *N*,*N*-diisopropylethylamine (1.01 mL, 5.78 mmol, 4.40 equiv.). The headspace was purged with argon gas for ~5 min and the reaction mixture was heated to 60 °C and stirred for 5 min, after which the reaction was deemed complete by HPLC. The reaction mixture was then concentrated *in vacuo* and the residual red-orange oil was reconstituted in DMF (2 mL) and pivalic anhydride (906 µL, 4.47 mmol, 3.40 equiv.) was added. The reaction mixture was heated to 60 °C and stirred. After ~2 min, a lot of orange precipitate is observed, likely the pivaloylated product. Chloroform (3 mL) was added and the majority of solid dissolved. After 10 minutes, when the reaction was deemed complete by HPLC, the crude reaction mixture was diluted with EtOAc (~50 mL) and water (~20 mL), transferred to a separatory funnel, rinsing the vessel repeatedly with 0.1 M HCl and EtOAc (final volume ~200 mL). The aqueous layer was removed, and the organic layer was washed with 0.1 M $H_3PO_4$ (2 x 15 mL) and brine (2 x 20 mL), dried over anhydrous sodium sulfate, filtered, and concentrated *in vacuo* to yield **S2** (1.115 g, 1.588 mmol, 121%) as an orange solid that was used without further purification.

To a 100 mL round-bottom flask equipped with a magnetic stir bar was added **S2** (823 mg, 1.17 mmol, 1.00 equiv.), *N*-hydroxysuccinimide (216 mg, 1.87 mmol, 1.60 equiv.), and DMF (5 mL). The resulting solution was cooled to 0 °C in an ice-water bath and EDC-HCl (292 mg, 1.52 mmol, 1.30 equiv.) was charged as a solid. After ~5 min, the ice-water bath was removed and the reaction mixture was stirred overnight at ambient temperature. Silica gel (6.0 g) and MeCN were added and the crude reaction mixture was concentrated *in vacuo* for solid loading on silica column (45 °C water bath, max vacuum). Purification by silica gel chromatography (4.5 x 13 cm, solid load, collected in 1.6 x 15 cm tubes) eluting with 20-60% EtOAc/pet ether yielded 1.060 g of light orange solid contaminated with NHS and $Piv_2O$. This mixed material was repurified by silica gel chromatography (25% acetone/hexanes) to yield the title compound as an orange solid (459 mg, 1.17 mmol, 49%). $^1H$ NMR (400 MHz, $CDCl_3$) δ 8.41 (dd, *J* = 8.0, 1.4 Hz, 1H), 8.19 (dd, *J* = 8.0, 0.8 Hz, 1H), 7.94 (dd, *J* = 1.4, 0.8 Hz, 1H), 6.92 (d, *J* = 8.7 Hz, 2H), 6.78 (d, *J* = 8.7 Hz, 2H), 2.96 (s, 4H), 1.42 (s, 18H).

**4',5'-dibromo-6-((2,2-dimethyl-4-oxo-3,9,12,15-tetraoxa-5-azaoctadecan-18-yl)carbamoyl)-3-oxo-3*H*-spiro[isobenzofuran-1,9'-xanthene]-3',6'-diyl bis(2,2-dimethylpropanoate) (S4).** **S3** (120 mg, 0.151 mmol, 1 equiv.) was added to an oven-dried 5 mL round-bottom flask and dissolved in DCM (2.0 mL) under an atmosphere of argon. To a 1 dram septum-cap glass vial was added tert-butyl (3-(2-(2-(3-aminopropoxy)ethoxy)ethoxy)propyl)carbamate (142 mg, 0.444 mmol, 2.95 equiv., $NH_2$-TOTA-NHBoc) and DCM (0.5 mL). The amine solution was transferred to the solution of NHS ester dropwise over ~2 min, the color changing to a bright orange immediately after the first drop was added. The 1 dram vial was rinsed with DCM (0.5 mL). The resulting reaction mixture was stirred at 0 °C. A few minutes later, *N*,*N*-dimethylpyridin-4-amine (18 mg, 0.15 mmol, 1.0 equiv.) was added as a solid in a single portion. The headspace was purged with argon for ~2 min, and the reaction mixture was stirred at 0 °C for 5 min before the cooling bath was removed. After 30 min, when the starting material had been consumed as judged by HPLC analysis, the crude reaction mixture was concentrated and purified by silica gel chromatography (2.5 x 15 cm, wet load with ~5 mL DCM, collected in 1.6 x 10 cm tubes) eluting with 30% acetone/hexanes (400 mL) to yield the title compound (54 mg, 53 µmol, 35%). The de-pivaloylated material was also collected and converted back to **S4** by dissolving in DMF (2.0 mL), charging pivalic anhydride (95 mg, 0.51 mmol, 3.4 equiv.), pyridine (50 µL, 0.62 mmol, 4.1 equiv.) and heating the bright red-orange solution to 60 °C. HPLC analysis after 5 min showed consumption of the starting material. Silica gel (3.0 g) was charged to the solution and it was concentrated *in vacuo* to dryness at 50 °C. The crude material was purified twice by silica gel chromatography, eluting with 10–30% acetone/hexanes (400 mL), and combined with the initial material to yield the title compound as a yellow oil (103 mg, 0.102 mmol, 68%). $^1H$ NMR (400 MHz, $CDCl_3$) δ 8.22 (dd, *J* = 8.0, 1.4 Hz, 1H), 8.08 (dd, *J* = 8.1, 0.7 Hz, 1H), 7.74 (dd, *J* = 1.4, 0.8 Hz, 1H), 7.57 (t, *J* = 5.8 Hz, 1H), 6.90 – 6.79 (m, 4H), 6.41 (s, 1H), 4.88 (d, *J* = 31.1 Hz, 2H), 3.69 – 3.48 (m, 12H), 3.47 – 3.42 (m, 2H), 3.36 (ddd, *J* = 7.0, 3.9, 1.9 Hz, 3H), 3.34 – 3.29 (m, 2H), 3.21 (t, *J* = 6.2 Hz, 2H), 3.18 – 3.14 (m, 2H), 3.04 (s, 2H), 1.86 – 1.69 (m, 9H), 1.54 (p, *J* = 6.3 Hz, 2H), 1.41 (s, 18H), 1.18 (s, 7H).

**6-((3-(2-(2-(3-aminopropoxy)ethoxy)ethoxy)propyl)carbamoyl)-4',5'-dibromo-3-oxo-3*H*-spiro[isobenzofuran-1,9'-xanthene]-3',6'-diyl bis(2,2-dimethylpropanoate) (S5).** To a 2 dram glass vial containing **S4** (113 mg, 0.112 mmol, 1 equiv.) was added a magnetic stir bar, MeCN (3 mL), and HCl (0.75 mL, 6 M, aq.) dropwise at 22 °C. The resulting reaction mixture was heated to 35 °C and stirred for 60 min until complete consumption of starting material was judged by TLC.

The crude reaction mixture was concentrated *in vacuo* to yield the crude HCl salt of the title compound as a yellow solid which was used without further purification.

To a 2 dram glass vial was charged cholenic acid (46 mg, 0.12 mmol, 1.1 equiv.) and THF (4 mL) to suspend. The resulting suspension was cooled to 0 °C in an ice-water bath, and isobutyl chloroformate (18 µL, 0.13 µmol, 1.2 equiv.) and triethylamine (50 µL, 0.36 mmol, 3.2 equiv.) were added sequentially. The reaction was then stirred at 0 °C for 7 min before the ice-water bath was removed, allowing the reaction to warm to ambient temperature over ~10 min. To a 50 mL round-bottom flask containing **S5** (106 mg, 0.112 mmol, 1.00 equiv.) was added THF (6 mL) and triethylamine (50 µL, 0.36 mmol, 3.2 equiv.). The resulting solution was cooled to 0 °C in an ice-water bath, the suspension containing the mixed anhydride of cholenic acid was added, rinsing with THF (2 mL), and the resulting reaction mixture was stirred at 0 °C. The reaction stalled, so additional triethylamine (50 µL, 0.36 mmol, 3.2 equiv.) was added and the ice-water bath was removed. After 1 h, when the disappearance of starting material was observed by HPLC, the reaction was concentrated to ~1/3 volume *in vacuo* and a biphasic mix of water (10 mL) and DCM (10 mL) was added to the white suspension and stirred. The aqueous layer was acidified to pH~2 via HCl (6 M, aq.) and then transferred to a separatory funnel. The organic layer was removed, and the aqueous layer was extracted with DCM (2 x 10 mL). The organic components were combined, washed once with a mixture of brine and dilute HCl (~10:1 v:v) and once with brine (~20 mL), dried over anhydrous sodium sulfate, gravity filtered, and concentrated *in vacuo* to yield a crude yellow oil. Purification by silica gel chromatography (2.5 x 16.5 cm, wet load with ~2 mL DCM, collected in 1.6 x 10 cm tubes, eluting with 30% acetone hexanes (130 mL), 40% acetone hexanes (250 mL), and 50% acetone hexanes (500 mL)) yielded **S6** (107 mg, 84.8 µmol, 76%) as a white crystalline solid. HRMS *m/z* (ESI$^+$) calculated for $C_{65}H_{85}Br_2N_2O_{13}^+$ ([M+H]$^+$): 1261.4393, observed 1261.4600; calculated for $C_{65}H_{84}Br_2N_2NaO_{13}$ ([M+Na]): 1283.4212, observed 1283.4421.

**4',5'-dibromo-6-(((*R*)-18-((3*S*,8*S*,9*S*,10*R*,13*R*,14*S*,17*R*)-3-hydroxy-10,13-dimethyl-2,3,4,7,8,9,10,11,12,13,14,15,16,17-tetradecahydro-1*H*-cyclopenta[*a*]phenanthren-17-yl)-15-oxo-4,7,10-trioxa-14-azanonadecyl)carbamoyl)-3-oxo-3*H*-spiro[isobenzofuran-1,9'-xanthene]-3',6'- diyl diacetate (S7).** To a 2 dram glass vial containing **S6** (100 mg, 0.079 mmol, 1 equiv.) were added MeOH (2 mL) and NaOH (50 µL, 1 M, aq.), and the solution was stirred for 5 min before it was concentrated *in vacuo*. To the residual red-orange solid was added DCM (10 mL) and water (5 mL). Then, acetyl chloride (54 µL, 0.76 mmol, 9.7 equiv.) was added and reaction mixture was stirred vigorously for 18 h at ambient temperature, after which the reaction was judged complete by HPLC. The aqueous layer was acidified with HCl (1 M, aq.) and the biphasic mixture was transferred to a separatory funnel. The aqueous layer was removed, and the organic layer was washed with a mixture of brine (~15 mL) and dilute HCl (~2 mL). The organic component was then dried over anhydrous sodium sulfate, gravity filtered, and concentrated *in vacuo* to yield a pale yellow oil. Purification by silica gel chromatography (2.5 x 16 cm, wet load with ~3 mL DCM, collected in 1.6 x 10 cm tubes, eluting with 40% acetone/hexanes (120 mL) and 60% acetone/hexanes (400 mL)) (Rf 0.21 in 50% acetone/hexanes) yielded the title compound (20 mg, 17 µmol, 21%) as a white solid. TLC $R_f$ (50% acetone/hexanes): 0.21. HRMS *m/z* (ESI$^+$) calculated for $C_{59}H_{72}Br_2N_2NaO_{13}$ ([M+Na]): 1199.3278, observed 1199.3372. $^1$H NMR (400 MHz,

$CDCl_3$) δ 8.22 (dd, *J* = 8.0, 1.4 Hz, 1H), 8.09 (dd, *J* = 8.0, 0.7 Hz, 1H), 7.81 – 7.78 (m, 1H), 7.76 (t, *J* = 5.3 Hz, 1H), 6.92 (d, *J* = 8.7 Hz, 2H), 6.84 (d, *J* = 8.7 Hz, 2H), 6.00 (d, *J* = 5.7 Hz, 1H), 5.35 (d, *J* = 4.7 Hz, 1H), 3.54 (dt, *J* = 15.0, 5.5 Hz, 4H), 3.45 (dd, *J* = 5.9, 3.1 Hz, 2H), 3.41 – 3.27 (m, 4H), 3.25 – 3.08 (m, 6H), 2.39 (s, 6H), 2.34 – 2.24 (m, 1H), 2.17 (d, *J* = 2.3 Hz, 2H), 2.09 (s, 1H), 2.02 – 1.90 (m, 2H), 1.88 – 1.80 (m, 4H), 1.78 – 1.32 (m, 10H), 1.32 – 1.03 (m, 10H), 1.00 (s, 3H), 0.97 – 0.81 (m, 6H), 0.66 (s, 3H). $^{13}C$ NMR (101 MHz, $CDCl_3$) δ 173.84, 168.29, 168.11, 165.08, 153.26, 150.73, 148.87, 142.24, 140.90, 130.37, 127.45, 127.41, 125.65, 122.66, 121.79, 119.73, 117.70, 106.66, 81.41, 71.91, 70.49, 69.92, 69.72, 69.46, 69.29, 56.88, 55.99, 50.23, 42.51, 42.44, 39.91, 39.56, 37.40, 37.24, 36.64, 35.67, 33.70, 32.02, 31.97, 31.79, 29.37, 28.60, 28.30, 24.40, 21.22, 20.93, 19.54, 18.58, 12.03.

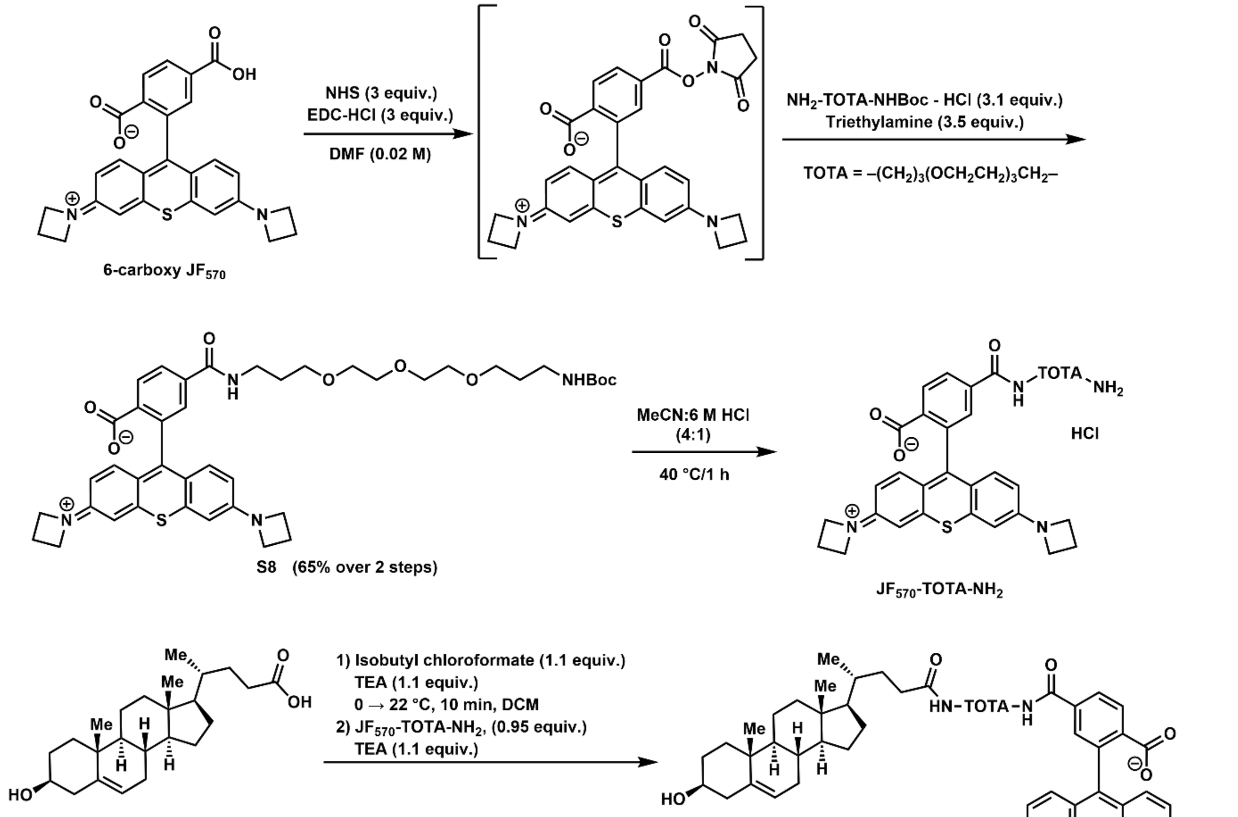


**2-(3-(azetidin-1-ium-1-ylidene)-6-(azetidin-1-yl)-3*H*-thioxanthen-9-yl)-4-((2,2-dimethyl-4-oxo-3,9,12,15-tetraoxa-5-azaoctadecan-18-yl)carbamoyl)benzoate (S8).** To a 20 mL glass vial containing 2-(3-(azetidin-1-ium-1-ylidene)-6-(azetidin-1-yl)-3H-thioxanthen-9-yl)-4-carboxybenzoate (44 mg, 0.094 mmol, 1.0 equiv., 6-carboxy $JF_{570}$) was added DMF (1 mL) and *N*-hydroxysuccinimide (32 mg, 0.28 mmol, 3.0 equiv.) and the resulting solution was cooled to 0 °C in an ice-water bath. After ~3 min, EDC-HCl (54 mg, 0.28 mmol, 3.0 equiv.) was charged as a solid. The headspace was purged with argon gas for ~2 min, the ice-water bath was removed, and the reaction mixture was stirred for 22 h under an inert atmosphere. At that point, HPLC analysis indicated 82% conversion to the NHS ester (via relative integration of the 280 nm UV chromatogram). A solution of tert-butyl (3-(2-(2-(3-aminopropoxy)ethoxy)ethoxy)propyl)carbamate hydrochloride (103 mg, 0.290 mmol, 3.10 equiv., $NH_2$-TOTA-NHBoc) in anhydrous DMF (0.4 mL) and triethylamine (47 µL, 0.34 mmol, 3.6 equiv.) was charged dropwise over ~2 min to the crude reaction mixture containing the NHS ester, and the resulting reaction mixture was stirred at ambient temperature for 2 h. At that point, HPLC analysis showed complete consumption of the NHS ester and generation of a new peak with the expected mass (744.0 m/z observed by LRMS). The crude reaction mixture was concentrated in vacuo, and MeCN (2 x ~5 mL) was added and evaporated to drive off most of the DMF. Purification by preparative TLC (30% MeOH/DCM) yielded the title compound (47 mg, 61 µmol, 65%) as a dark purple solid. HRMS *m/z* ($ESI^+$) calculated for $C_{42}H_{53}N_4O_8S^+$ ($[M+H]^+$): 773.3578, observed 773.3543. $^1H$ NMR (300 MHz, MeOD) δ 8.18 – 8.01 (m, 2H), 7.61 (d, *J* = 1.7 Hz, 1H), 7.27 (d, *J* = 9.4 Hz, 2H), 6.81 (d, *J* = 2.3 Hz, 2H), 6.57 (dd, *J* = 9.4, 2.3 Hz, 2H), 4.24 (t, *J* = 7.6 Hz, 8H), 3.70 – 3.40 (m, 15H), 3.35 (s, 2H), 3.10 (dt, *J* = 12.2, 6.8 Hz, 3H), 2.60 – 2.43 (m, 4H), 1.96 – 1.79 (m, 2H), 1.71 (dp, *J* = 19.8, 6.6 Hz, 3H), 1.41 (s, 10H).

**2-(6-amino-3-iminio-3*H*-thioxanthen-9-yl)-4-(((*R*)-18-((3*S*,8*S*,9*S*,10*R*,13*R*,14*S*,17*R*)-3-hydroxy-10,13-dimethyl-2,3,4,7,8,9,10,11,12,13,14,15,16,17-tetradecahydro-1*H*-cyclopenta[*a*]phenanthren-17-yl)-15-oxo-4,7,10-trioxa-14-azanonadecyl)carbamoyl)benzoate (1, Chol-$JF_{570}$).** To a solution of **S8** (47 mg, 0.061 mmol, 1.0 equiv.) in 2 mL MeCN was charged HCl (0.50 mL, 6 M, aq.) dropwise, and the resulting purple solution was heated to 40 °C and stirred for 50 min, at which point LCMS analysis showed complete consumption of the starting material. The crude reaction mixture was then concentrated *in vacuo*, azeotroping with toluene (3x). The crude solid was then taken up in DCM (2 mL) and triethylamine (34 µL, 0.25 mmol, 4.0 equiv.) was added to generate a solution of the free base ($JF_{570}$-TOTA-$NH_2$) which was cooled to 0 °C in an ice-water bath. In a separate 4 mL glass vial equipped with a magnetic stir bar, (R)-4-((3S,8S,9S,10R,13R,14S,17R)-3-hydroxy-10,13-dimethyl-2,3,4,7,8,9,10,11,12,13,14,15,16,17-tetradecahydro-1H-cyclopenta[a]phenanthren-17-yl)pentanoic acid (24.1 mg, 1.05 Eq, 64.3 µmol, cholenic acid) was suspended in THF (2 mL) and triethylamine (18 µL, 0.13 mmol, 2.1 equiv.) was added. The resulting suspension was cooled to 0 °C in an ice-water bath and stirred under an inert atmosphere. After 10 min, isobutyl chloroformate (8.8 µL, 0.067 mmol, 1.1 equiv.) was added, and the resulting solution was stirred at 0 °C for 10 min. The cold solution of the mixed anhydride was then added to the stirring solution of $JF_{570}$-TOTA-$NH_2$ free base, the ice-water bath was removed, and the reaction mixture was stirred for 12 h, whereupon HPLC analysis indicated complete consumption of starting material. The crude reaction mixture was concentrated *in vacuo* to yield a crude deep purple oil. Purification by preparative TLC (DCM:MeOH:AcOH 95:5:0.1) yielded the title compound (20 mg, 0.019 mmol, 33%) as a deep purple solid. TLC $R_f$ (DCM:MeOH:AcOH 95:5:0.1): 0.35. $^1H$ NMR (500 MHz, $CDCl_3$/MeOH-*d*4) δ 8.06 (br s, 1 H), 7.86 (s, 1H), 7.46 (s, 1H), 7.06 (s, 2H), 6.71 (s, 2H), 6.53 (s, 2H), 5.17 (d, *J* = 5.2 Hz, 1H), 3.46 (t, *J* = 4.7 Hz, 13H), 3.35 (ddd, *J* = 23.5, 10.1, 5.2 Hz, 6H), 3.20 (s, 4H), 3.07 (d, *J* = 6.5 Hz, 2H), 2.86 (q, *J* = 7.0 Hz, 1H), 2.07 (h, *J* = 9.6 Hz, 7H), 1.93 – 1.86 (m, 4H), 1.81 – 1.50 (m, 7H), 1.47 – 1.18 (m, 6H), 1.10 (q, *J* = 7.7 Hz, 4H), 0.98 – 0.86 (m, 2H), 0.84 (d, *J* = 3.4 Hz, 3H), 0.76 (d, *J* = 6.2 Hz, 4H), 0.51 (d, *J* = 3.7 Hz, 3H). $^{13}C$ NMR (126 MHz, $CDCl_3$)

δ 166.64, 153.25, 143.79, 140.77, 135.79, 135.43, 128.29, 127.31, 121.32, 119.54, 116.53, 104.10, 77.42, 77.36, 77.16, 76.90, 71.14, 70.21, 70.14, 69.91, 69.84, 69.62, 69.18, 56.61, 55.72, 49.99, 45.58, 42.22, 41.92, 41.70, 40.02, 39.63, 38.10, 37.13, 36.91, 36.35, 35.46, 33.30, 31.86, 31.74, 31.71, 31.04, 31.01, 28.87, 28.76, 27.98, 24.09, 20.90, 19.14, 18.11, 11.63, 8.44. HRMS *m/z* (ESI$^+$) calculated for $C_{61}H_{83}Cl_2N_4O_8S^+$ ([M+H+2HCl]$^+$): 1101.5303, observed 1101.5376.

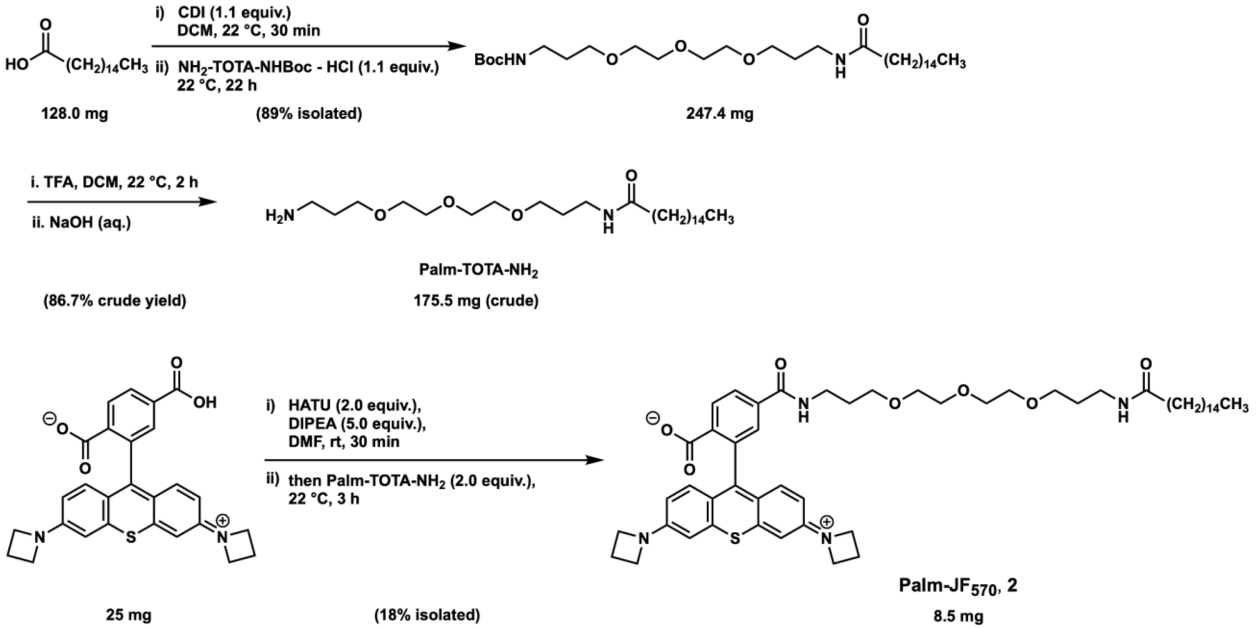


tert-butyl (15-oxo-4,7,10-trioxa-14-azatriacontyl)carbamate (**S9**).

To an oven-dried 5-mL flask equipped with a stir bar was added palmitic acid (128 mg, 1 Eq, 0.500 mmol), CDI (89.2 mg, 1.1 Eq, 550 µmol), and dry DCM (3.0 mL) under Ar. The mixture was stirred for 30 min, then tert-butyl (3-(2-(2-(3-aminopropoxy)ethoxy)ethoxy)propyl)carbamate (176 mg, 1.1 Eq, 550 µmol) was added, and the pale yellow solution was stirred under argon atmosphere for 22 h. The reaction was diluted with DCM, quenched with NH4Cl, then acidified to pH ~5 by adding a few drops of 1M HCl and transferred to a separatory funnel. The organic layer was collected and the product was stripped from the aqueous layer with DCM (3x). The organic layers were combined, washed with an equivalent volume of 1M HCl and extracted with DCM (3x). The organics were then combined, dried over anhydrous sodium sulfate, gravity filtered, concentrated by rotary evaporation, and placed under high vacuum (with gentle heating) for 5 minutes to yield tert-butyl (15-oxo-4,7,10-trioxa-14-azatriacontyl)carbamate (247 mg, 442.7 µmol, 89%) as a pale yellow gel that solidifies upon cooling. NMR matches the previously-reported[31] values. $^{1}$H NMR (400 MHz, MeOD) δ 3.64 (tdd, *J* = 5.1, 1.8, 1.0 Hz, 4H), 3.61 – 3.55 (m, 4H), 3.51 (td, *J* = 6.2, 3.3 Hz, 4H), 3.25 (t, *J* = 6.8 Hz, 2H), 3.12 (t, *J* = 6.8 Hz, 2H), 2.27 (t, *J* = 7.4 Hz, 1H), 2.17 (dd, *J* = 8.0, 6.9 Hz, 2H), 1.74 (dt, *J* = 13.1, 6.4 Hz, 4H), 1.60 (t, *J* = 7.4 Hz, 3H), 1.43 (s, 9H), 1.36 – 1.22 (m, 26H), 0.90 (t, *J* = 6.8 Hz, 3H).

N-(3-(2-(2-(3-aminopropoxy)ethoxy)ethoxy)propyl)palmitamide (**S10**).

To a scintillation vial containing tert-butyl (15-oxo-4,7,10-trioxa-14-azatriacontyl)carbamate (**S9**) (246.6 mg, 1 Eq, 441.3 µmol) was added DCM (1 mL), TFA (1 mL), and a stir bar. The resulting pink solution was stirred for 2 h. The stir bar was removed and the crude reaction mixture was concentrated by rotary evaporation to yield an off-white solid, which was then partitioned between DCM and 1M NaOH and extracted with DCM (3x). The organic layers were combined, concentrated by rotary evaporation, and dried under high vacuum to yield the free amine N-(3-(2-(2-(3-aminopropoxy)ethoxy)ethoxy)propyl)palmitamide (**S10**) (176 mg, 382.6 µmol, 87%) as a white solid. $^1$H NMR (600 MHz, MeOD) δ 3.67 (t, *J* = 5.5 Hz, 2H), 3.66 – 3.62 (m, 6H), 3.60 – 3.58 (m, 2H), 3.51 (t, *J* = 6.1 Hz, 2H), 3.25 (t, *J* = 7.1 Hz, 2H), 3.10 (t, *J* = 6.4 Hz, 2H), 2.28 (t, *J* = 7.4 Hz, 1H), 2.20 – 2.15 (m, 2H), 1.93 (qd, *J* = 6.1, 5.1 Hz, 2H), 1.75 (tt, *J* = 7.0, 6.1 Hz, 2H), 1.64 – 1.55 (m, 3H), 1.29 (s, 26H), 0.90 (t, *J* = 7.0 Hz, 3H).

2-(3-(azetidin-1-ium-1-ylidene)-6-(azetidin-1-yl)-3H-thioxanthen-9-yl)-4-((15-oxo-4,7,10-trioxa-14-azatriacontyl)carbamoyl)benzoate ('Palm-$JF_{570}$', **2**).

To a 50-mL round-bottom flask containing a magnetic stir bar was added 1-(6-(azetidin-1-yl)-9-(2,5-dicarboxyphenyl)-3H-thioxanthen-3-ylidene) azetidin-1-ium (17 mg, 1 Eq, 36 μmol) and anhydrous DMF (6.0 mL). DIPEA (47 mg, 63 μL, 10 equiv., 0.36 mmol), HATU (27 mg, 2 equiv., 72 μmol) and N-(3-(2-(2-(3-aminopropoxy)ethoxy)ethoxy)propyl)palmitamide (33 mg, 2 equiv., 72 μmol) were added successively. After stirring overnight, the mixture was evaporated at 40 °C and the residual material was purified by preparative HPLC (5-95% acetonitrile/water with constant 0.1% formic acid additive) to yield 19 mg (58%) of Palm-$JF_{570}$ **2** as a dark purple solid. HRMS expected: ($C_{53}H_{75}N_4O_7S^+$): 911.5351 m/z; Observed: 911.5345 m/z $[M+H]^+$. $^1$H NMR (500 MHz, MeOD) δ 8.40 (d, *J* = 8.3 Hz, 1H), 8.19 (dd, *J* = 8.3, 1.8 Hz, 1H), 7.75 (d, *J* = 1.8 Hz, 1H), 7.15 (d, *J* = 9.4 Hz, 2H), 6.88 (d, *J* = 2.3 Hz, 2H), 6.62 (dd, *J* = 9.4, 2.3 Hz, 2H), 4.27 (t, *J* = 7.7 Hz, 8H), 3.72 – 3.57 (m, 2H), 3.60 – 3.54 (m, 4H), 3.55 – 3.40 (m, 6H), 3.25 (td, *J* = 6.9, 2.3 Hz, 2H), 3.21 (t, *J* = 6.9 Hz, 2H), 2.53 (p, *J* = 7.7 Hz, 2H), 2.17 (t, *J* = 7.6 Hz, 1H), 2.13 (t, *J* = 7.5 Hz, 2H), 2.03 (s, 1H), 1.93 (s, 1H), 1.86 (q, *J* = 6.4 Hz, 2H), 1.73 (dp, *J* = 23.8, 6.5 Hz, 4H), 1.28 (d, *J* = 10.0 Hz, 28H), 0.94 – 0.84 (m, 4H). $^{13}$C NMR (126 MHz, MeOD) δ 176.19, 167.68, 167.28, 162.01, 154.24, 144.87, 139.41, 139.31, 136.62, 132.91, 130.43, 129.52, 120.09, 114.71, 104.57, 71.53, 71.41, 71.39, 71.22, 71.21, 71.19, 71.12, 70.19, 69.89, 69.86, 66.60, 52.73, 39.01, 37.90, 37.73, 37.19, 37.17, 33.08, 30.81, 30.79, 30.77, 30.73, 30.64, 30.48, 30.44, 30.43, 30.36, 30.30, 27.07, 23.74, 16.84, 14.45.

**2-(3-(but-3-yn-1-yl)-3H-diazirin-3-yl)ethyl (R)-4-((3S,8S,9S,10R,13R,14S,17R)-3-hydroxy-10,13-dimethyl-2,3,4,7,8,9,10,11,12,13,14,15,16,17-tetradecahydro-1H-cyclopenta[a]phenanthren-17-yl)pentanoate — "NBII-165".** The alkynyl diazirine alcohol 2-(3-(but-3-yn-1-yl)-3H-diazirin-3-yl)ethan-1-ol was synthesized according to methods described by Li et al.[5] To an oven dried glass vial was added cholenic acid (50 mg, 1 Eq, 0.13 mmol) and DMAP (3.3 mg, 0.2 Eq, 27 μmol) under Ar. 2-(3-(but-3-yn-1-yl)-3H-diazirin-3-yl)ethan-1-ol (20 mg, 1.1 Eq, 0.15 mmol) in DCM (0.30 mL). The solution was then cooled in an ice-water bath and DCC (41 mg, 1.5 Eq, 0.20 mmol) in DCM (0.2 mL) was added dropwise. The ice-water bath was removed and the mixture was stirred for 18 h before it was clarified by filtering through cotton. The residual oil was purified by column chromatography (2.5 x 15 cm, wet load with DCM) eluting with 30% EtOAc/hexanes (200 mL) and 50% EtOAc/hexanes (200 mL) to yield 52 mg (79%) of 2-(3-(but-3-yn-1-yl)-3H-diazirin-3-yl)ethyl(4R)-4-(3-hydroxy-10,13-dimethyl-2,3,4,7,8,9,10,11,12,13,14,15,16,17-tetradecahydro-1H-cyclopenta[a]phenanthren-17-yl)pentanoate as a light yellow solid. HRMS expected: 495.3581 ($C_{31}H_{47}N_2O_3^+$); observed: 495.3625. $^1$H NMR (400 MHz, $CDCl_3$) δ 5.35 (d, *J* = 5.2 Hz, 1H), 3.97 (t, *J* = 6.4 Hz, 1H), 3.52 (dd, *J* = 14.6, 8.9 Hz, 1H), 2.37 (ddd, *J* = 15.4, 10.2, 5.2 Hz, 1H), 2.33 – 2.18 (m, 2H), 2.06 – 1.92 (m, 3H), 1.91 – 1.77 (m, 3H), 1.74 (t, *J* = 6.4 Hz, 1H), 1.68 (t, *J* = 7.4 Hz, 1H), 1.52 – 1.40 (m, 2H), 1.40 – 1.24 (m, 1H), 1.22 – 1.03 (m, 3H), 1.01 (s, 3H), 0.93 (t, *J* = 6.0 Hz, 3H), 0.68 (d, *J* = 2.5

Hz, 3H). $^{13}C$ NMR (101 MHz, $CDCl_3$) δ 174.19, 140.90, 121.81, 82.70, 71.94, 69.42, 59.06, 56.88, 55.90, 50.23, 42.53, 42.44, 39.89, 37.40, 36.64, 35.49, 32.48, 32.45, 32.04, 32.02, 31.81, 31.33, 31.09, 28.26, 26.51, 24.41, 21.22, 19.54, 18.47, 13.40, 12.03.

### 3. Supplementary Tables

**Table S1**: List of primers used in this study.

| Primer | Sequence | Note |
|---|---|---|
| 1 | tagaagacaccgactgccaccatggaacaaaagctga | Forward primer for subcloning the HaloTag-mAster-B insert into lentiviral backbone plasmid 7 by Gibson assembly. |
| 2 | gcggccgctttactttcaatgatagcggttcctcttctcg | Reverse primer for subcloning the HaloTag-mAster-B insert into lentiviral backbone plasmid 7 by Gibson assembly. |
| 3 | cccggacctgatcggcagcgagatcgcgcgctggc | Custom sequencing primer used for sequencing N-terminal HaloTag constructs contained in plasmid 1. |
| 4 | cacctgctacgggaacgagctgggcc | Forward primer for TOPO cloning to make an entry vector containing the ΔGRAM-mAster-B insert. |
| 5 | tcaatgatagcggttcctcttctcg | Reverse primer for TOPO cloning to make an entry vector containing the ΔGRAM-mAster-B insert. |
| 6 | caccggaggtggtggttctccgg | Forward primer for TOPO cloning to make an entry vector containing the mAster-B-ΔERD insert. |
| 7 | tcatgctgaagacttgctagccttctcc | Reverse primer for TOPO cloning to make an entry vector containing the mAster-B-ΔERD insert. |

Table S2: List of plasmids used in this study.

| # | Protein expressed/Gene insert | Backbone | Notes |
|---|---|---|---|
| 1 | HaloTag | pMH-HaloTag | Addgene #154144, Huen Lab. Gateway destination vector for making N-terminal HaloTag fusions. N-terminal Myc tag. |
| 2 | NUP153 | pDONR221 | Gateway entry clone. DNASU HsCD00860103 |
| 3 | NUP153-HaloTag | pMH-HaloTag | Gateway LR reaction between plasmids 1 & 2. |

| | | | |
|---|---|---|---|
| 4 | HaloTag-eGFP-mito (pERB254) | pEM791 | Addgene #67762, Ballister et al. C-terminal inner mitochondrial membrane targeting sequence. HaloTag-eGFP-mito. |
| 5 | mAster-B | pDONR221 | Reported previously [32]. The Aster-B gene was PCR amplified from *Mus musculus* C57BL/6J cDNA and cloned into pDonr221 by Gateway cloning (BP reaction, ThermoFisher). Gateway entry clone with N-terminal HA tag. |
| 6 | HaloTag-mAster-B | pMH-HaloTag | Gateway between plasmids 1 & 6. |
| 7 | eGFP | p156RRL-sinPPT-CMV-GFP-PRE/Nhe I | Addgene #17446. Campeau & Kaufmann |
| 8 | HaloTag-mAster-B | p156RRL-sinPPT-CMV-PRE/Nhe I | Lentiviral vector for transduction. Prepared using primers 1 & 2 on plasmid 6, then Gibson Assembly of amplicon into plasmid 7 to replace eGFP. |
| 9 | ΔGRAM-mAster-B | pENTR™/D-TOPO™ | Directional TOPO (ThermoFisher K240020) to make gateway entry vector using primers 4 & 5 on plasmid 5. |
| 10 | ΔGRAM-mAster-B | pMH-HaloTag | Gateway LR reaction between plasmids 1 and 9. |
| 11 | mAster-B-ΔERD | pENTR™/D-TOPO™ | Directional TOPO (ThermoFisher K240020) to make gateway entry vector using primers 6 & 7 on plasmid 5. |
| 12 | mAster-B-ΔERD | pMH-HaloTag | Gateway LR reaction between plasmids 2 and 11. |
| 13 | eGFP-mAsterB | pDest53 | Reported previously[32]. Prepared by gateway LR recombination into a pDest53 vector containing a CMV promoter and an N-terminal eGFP. |
| 14 | C-FLAG Destination | pRK5 | Gateway destination plasmid for the expression of C-terminally FLAG-tagged proteins. Gift from T Wucherpfennig. |

| | | | |
|---|---|---|---|
| 15 | pDONR_221_EMC7-FLAG | pENTR_223 | Gateway entry vector containing EMC7 (C15orf24). DNASU HsCD00515064 |
| 16 | EMC7-FLAG | pRK5 | Expression vector for C-terminally FLAG-tagged EMC7. |

**Table S3**: List of primary antibodies used in this study.

| Antibody | Source (Product #, production) | Validation | Dilution | Temp (°C) | Duration (ON = overnight) |
|---|---|---|---|---|---|
| β-actin | Cell Signaling Technology (3700S, mAb) | Yes | 1:20000 | 4 | ON |
| Myc | Cell Signaling Technology (2276S, mAb) | Yes (simpleChIP) | 1:133000 | 4 | ON |
| GFP | AbClonal (AE012, mAb) | Yes | 1:10000 | 20 | 1 h |
| GAPDH | Proteintech (60004-1-Ig, mAb) | Yes (KD/KO) | 1:20000 | 4 | ON |
| SR-B1 | Abcam (ab217318, recomb. mAb) | Yes (KO) | 1:4000 | 4 | 2 d |
| HaloTag | Promega (G9211, mAb) | Yes | 1:2000 | 4 | 2 d |
| FLAG | Thermo Scientific (F1804, mAb) | Yes | 1:10000 | 4 | ON |
| OSBP | Proteintech (11096-1-AP, pAb) | Yes (KD/KO) | 1:2000 | 4 | ON |
| EMC7 | Proteintech (27550-1-AP, pAb) | Yes (PMIDs: 32459176, 37199759) | 1:3000 | 4 | ON |
| EMC2 | Proteintech (25443-1-AP, pAb) | Yes (KD/KO) | 1:2500 | 4 | ON |
| FLOT1 (IF) | Cell Signaling (18634T, recomb. mAb) | Yes (PMID: 37996944) | 1:400 | 4 | ON |

| | | | | | |
|---|---|---|---|---|---|
| HA | Cell Signaling (3724S, recomb. mAb) | Yes | 1:1000 | 4 | ON |
| His-tag | Cell Signaling (2366S, mAb) | Yes | 1:1000 | 4 | ON |
| FLOT1 (WB) | Cell Signaling (3253S,mAb) | Yes | 1:1000 | 4 | ON |

**Table S4**. List of proteomics files used in this report, grouped by their respective association with Figures in the main text or supplementary information. Raw and search files can be found on the PRoteomics IDEntification Database (https://www.ebi.ac.uk/pride/) with the identifier PXD054875.

| Figure | File Name | Experiment | Description |
|---|---|---|---|
| 1 | HaloTag-ethynylaniline-1<br>HaloTag-ethynylaniline-2<br>HaloTag-ethynylaniline-3<br>HaloTag-NUP153-ethynylaniline-1<br>HaloTag-NUP153-ethynylaniline-2<br>HaloTag-NUP153-ethynylaniline-3 | Halo-POCA: HaloTag vs HaloTag-NUP153, 3EA capture, LFQ | HEK293T cells transfected with either free HaloTag (1–3) or HaloTag-NUP153 (4–6). Treated with $JF_{570}$-HTL, irradiated in the presence of 3-ethynylaniline. Protein-level enrichment |
| 1 | HaloTag-propargylamine-1<br>HaloTag-propargylamine-2<br>HaloTag-propargylamine-3<br>HaloTag-NUP153-propargylamine-1<br>HaloTag-NUP153-propargylamine-2<br>HaloTag-NUP153-propargylamine-3 | Halo-POCA: HaloTag vs HaloTag-NUP153, PA capture, LFQ | HEK293T cells transfected with either free HaloTag (1–3) or HaloTag-NUP153 (4–6). Treated with $JF_{570}$-HTL, irradiated in the presence of propargylamine. Protein-level enrichment |
| 2 | Chol-JF570_nohv_1<br>Chol-JF570_nohv_2<br>Chol-JF570_nohv_3<br>Chol-JF570_hv_1<br>Chol-JF570_hv_2<br>Chol-JF570_hv_3 | HEK293T mβCD-Chol probe **1,** +/- hv dataset #1 | HEK293T lipid starved in serum-free medium for 4 h, then incubated with 10 µM mβCD-chol-$JF_{570}$ (**1**) or mβCD-palm-$JF_{570}$ (**2**) for 1 h. 10 mM propargylamine/5 min hv. 200 ug input protein-level enrichment |
| 2 | Palm-JF570_nohv_1<br>Palm-JF570_nohv_2<br>Palm-JF570_nohv_3 | HEK293T mβCD-palm probe **2** | HEK293T lipid starved in serum-free medium for 4 h, then incubated |

| | | | |
|---|---|---|---|
| | Palm-JF570_hv_1<br>Palm-JF570_hv_2<br>Palm-JF570_hv_3 | | with 10 μM mβCD-palm-$JF_{570}$ (**2**) for 1 h. 10 mM propargylamine/5 min hv. 200 ug input protein-level enrichment |
| 2 | NBII-165_nohv_1<br>NBII-165_nohv_2<br>NBII-165_nohv_3<br>NBII-165_hv_1<br>NBII-165_hv_2<br>NBII-165_hv_3 | HEK293T mβCD-NBII-165 | HEK293T lipid starved in serum-free medium for 4 h, then incubated with 10 μM mβCD-NBII-165 for 1 h. Irradiated at 365 nm for 20 min. 200 ug input protein-level enrichment |
| 2 | LKM38_nohv_1<br>LKM38_nohv_2<br>LKM38_nohv_3<br>LKM38_hv_1<br>LKM38_hv_2<br>LKM38_hv_3 | HEK293T mβCD-LKM38 | HEK293T lipid starved in serum-free medium for 4 h, then incubated with 10 μM mβCD-LKM38 for 1 h. Irradiated at 365 nm for 20 min. 200 ug input protein-level enrichment |
| | trans-sterol_nohv_1<br>trans-sterol_nohv_2<br>trans-sterol_nohv_3<br>trans-sterol_hv_1<br>trans-sterol_hv_2<br>trans-sterol_hv_3 | HEK293T mβCD-trans-sterol | HEK293T lipid starved in serum-free medium for 4 h, then incubated with 10 μM mβCD-trans-sterol for 1 h. Irradiated at 365 nm for 20 min. 200 ug input protein-level enrichment |
| 3 | Chol-JF570_nohv_4<br>Chol-JF570_nohv_5<br>Chol-JF570_nohv_6<br>Chol-JF570_hv_4<br>Chol-JF570_hv_5<br>Chol-JF570_hv_6 | HEK293T mβCD-Chol probe **1,** +/- hv dataset #2 | HEK293T lipid starved in serum-free medium for 4 h, then incubated with 10 μM mβCD-chol-TOTA-$JF_{570}$ for 1 h. 10 mM propargylamine/5 min hv. 200 ug input protein-level enrichment |
| 3 | chol-JF570_noncompeted_1<br>chol-JF570_noncompeted_2<br>chol-JF570_noncompeted_3 | HEK293T mβCD-chol-TOTA-$JF_{570}$ | HEK293T lipid starved in serum-free medium for 4 h, then incubated |

| | | | |
|---|---|---|---|
| | chol-JF570_noncompeted_4<br>chol-JF570_competed_1<br>chol-JF570_competed_2<br>chol-JF570_competed_3<br>chol-JF570_competed_4 | cholesterol competition LFQ | with 10 µM mβCD-chol-TOTA-JF570 or probe + 100 µM mβCD-cholesterol for 1 h. 10 mM propargylamine/5 min hv. 200 ug input protein-level enrichment. Used 2x2 bio x technical replicates for each set |
| 3 | EMC7-FLAG_IP_notreat-1<br>EMC7-FLAG_IP_notreat-2<br>EMC7-FLAG_IP_notreat-3<br>EMC7-FLAG_IP_notreat-1_rep2<br>EMC7-FLAG_IP_notreat-2_rep2<br>EMC7-FLAG_IP_notreat-3_rep2<br>EMC7-FLAG_IP_chol-add-1<br>EMC7-FLAG_IP_chol-add-2<br>EMC7-FLAG_IP_chol-add-3<br>EMC7-FLAG_IP_chol-add-1_rep2<br>EMC7-FLAG_IP_chol-add-2_rep2<br>EMC7-FLAG_IP_chol-add-3_rep2<br>EMC7-FLAG_IP_chol-deplete-1<br>EMC7-FLAG_IP_chol-deplete-2<br>EMC7-FLAG_IP_chol-deplete-3<br>EMC7-FLAG_IP_chol-deplete-1_rep2<br>EMC7-FLAG_IP_chol-deplete-2_rep2<br>EMC7-FLAG_IP_chol-deplete-3_rep2 | EMC7-FLAG AP-MS with different amounts of cholesterol | HEK293T transiently overexpressing EMC7-FLAGcells were either loaded with mβCD-cholesterol (150 µM, 30 min), depleted of cholesterol with mβCD (2.5 mM, 30 min), or left untreated. Lysed into 0.4% NP40 and then proteins were pulled down with anti-FLAG resin. |
| 4 | Primary-cells_HDL-chol-JF570_nohv_1<br>Primary-cells_HDL-chol-JF570_nohv_2<br>Primary-cells_HDL-chol-JF570_nohv_3<br>Primary-cells_HDL-chol-JF570_hv_1<br>Primary-cells_HDL-chol-JF570_hv_2<br>Primary-cells_HDL-chol-JF570_hv_3 | 1° hepatocytes (wild-type), HDL-chol-TOTA-JF570 LFQ | Mouse primary hepatocytes, treated with 100 ug/mL [**HDL**-chol-TOTA-JF570], 3x +/- hv 10 mM propargylamine/5 min hv LFQ |
| 4 | HepG2_expression_1<br>HepG2_expression_2<br>HepG2_expression_3<br>HepG2-SR-B1-oe_expression_1<br>HepG2-SR-B1-oe_expression_2<br>HepG2-SR-B1-oe_expression_3 | HepG2-wt_SCARB1-oe_expression LFQ | HepG2 wild-type or SCARB1-overexpressing cells, 3 ug unenriched lysate analyzed |
| 4 | HepG2_HDL-chol-JF570_nohv_1<br>HepG2_HDL-chol-JF570_nohv_2<br>HepG2_HDL-chol-JF570_nohv_3<br>HepG2_HDL-chol-JF570_hv_1<br>HepG2_HDL-chol-JF570_hv_2 | HepG2-HDL-chol-TOTA-JF570 LFQ | HepG2 wild-type treated with 100 ug/mL [**HDL**-chol-TOTA-JF570], 3x +/- hv 10 mM propargylamine/5 min |

| | | | |
|---|---|---|---|
| | HepG2_HDL-chol-JF570_hv_3 | | hv LFQ |
| 4 | HepG2_SR-B1-oe_HDL-chol-JF570_nohv_1<br>HepG2_SR-B1-oe_HDL-chol-JF570_nohv_2<br>HepG2_SR-B1-oe_HDL-chol-JF570_nohv_3<br>HepG2_SR-B1-oe_HDL-chol-JF570_hv_1<br>HepG2_SR-B1-oe_HDL-chol-JF570_hv_2<br>HepG2_SR-B1-oe_HDL-chol-JF570_hv_3 | HepG2-SCARB1-oe HDL-chol-TOTA-$JF_{570}$ LFQ | HepG2-SRBI overexpressing treated with 100 ug/mL [**HDL**-chol-TOTA-$JF_{570}$], 3x +/- hv 10 mM propargylamine/5 min hv LFQ |
| 5 | HaloTag-Aster_1<br>HaloTag-Aster_2<br>HaloTag-Aster_3<br>HaloTag-Aster_pluschol_1<br>HaloTag-Aster_pluschol_2<br>HaloTag-Aster_pluschol_3 | HEK293T HaloTag-mAster-B, full-length, +/- cholesterol | HEK293T cells transiently expressing HaloTag-mAster-B lipid starved overnight then either loaded with mβCD-chol (100 µM) or not. Taken through Halo-POCA labeling protocol (total chol-loading time = 1 h) and protein-level enrichment and LFQ analysis. |
| 5 | HaloTag-GRAM-Aster_1<br>HaloTag-GRAM-Aster_2<br>HaloTag-GRAM-Aster_3<br>HaloTag-GRAM-Aster_pluschol_1<br>HaloTag-GRAM-Aster_pluschol_2<br>HaloTag-GRAM-Aster_pluschol_3 | HEK293T HaloTag-mAster-B-ΔERD, full-length, +/- cholesterol | HEK293T cells transiently expressing HaloTag-mAster-B-ΔERD lipid starved overnight then either loaded with mβCD-chol (100 µM) or not. Taken through Halo-POCA labeling protocol (total chol-loading time = 1 h) and protein-level enrichment and LFQ analysis. |
| S4 | C3_biotin-azide<br>C4_biotin-azide | BSA histidine capture *in vitro* | BSA (5 mg/mL in PBS) irradiated for 10 min in the presence of DBF (10 µM) and propargylamine (1 mM). Labeled proteins had either biotin-C3-N3 or biotin-C4-N3 appended by CuAAC. After cleanup and digestion, the modified peptides were enriched on |

| | | | |
|---|---|---|---|
| | | | neutravidin and then analyzed. |
| S15 | eGFP-HaloTag-mito_1<br>eGFP-HaloTag-mito_2<br>eGFP-HaloTag-mito_3<br>eGFP-HaloTag-mito_4 | HaloTag-eGFP-mito, +hv | HEK293T cells transfected with pERB254 HaloTag-eGFP-mito (pERB254). Subjected to Halo-POCA labeling and protein-level enrichment. |
| S30 | GFP-vs-EMC7_GFP-FLAG_IP_1<br>GFP-vs-EMC7_GFP-FLAG_IP_2<br>GFP-vs-EMC7_GFP-FLAG_IP_3<br>GFP-vs-EMC7_EMC7-FLAG_IP_1<br>GFP-vs-EMC7_EMC7-FLAG_IP_2<br>GFP-vs-EMC7_EMC7-FLAG_IP_3 | EMC7-FLAG AP-MS control experiment with GFP-FLAG comparator. LFQ-MBR | HEK293T cells transiently overexpressing either GFP-FLAG or EMC7-FLAG were lysed into 0.4% NP40 and then proteins were pulled down with anti-FLAG resin. |

**Table S5**: Gradient details for the LC-MS/MS method. Solvent A is 96.9% $H_2O$, 3% DMSO, and 0.1% formic acid. Solvent B is 80% acetonitrile, 16.9% $H_2O$, 3% DMSO, and 0.1% formic acid.

| Time (min) | %A | %B | Flow rate (nL/min) |
|---|---|---|---|
| 0 | 99 | 1 | 300 |
| 3 | 90 | 10 | 220 |
| 63 | 60 | 40 | 220 |
| 73 | 50 | 50 | 220 |
| 74 | 5 | 95 | 250 |
| 80 | 5 | 95 | 250 |
| 81 | 99 | 1 | 250 |
| 100 | 99 | 1 | 250 |

NMR SPECTRA

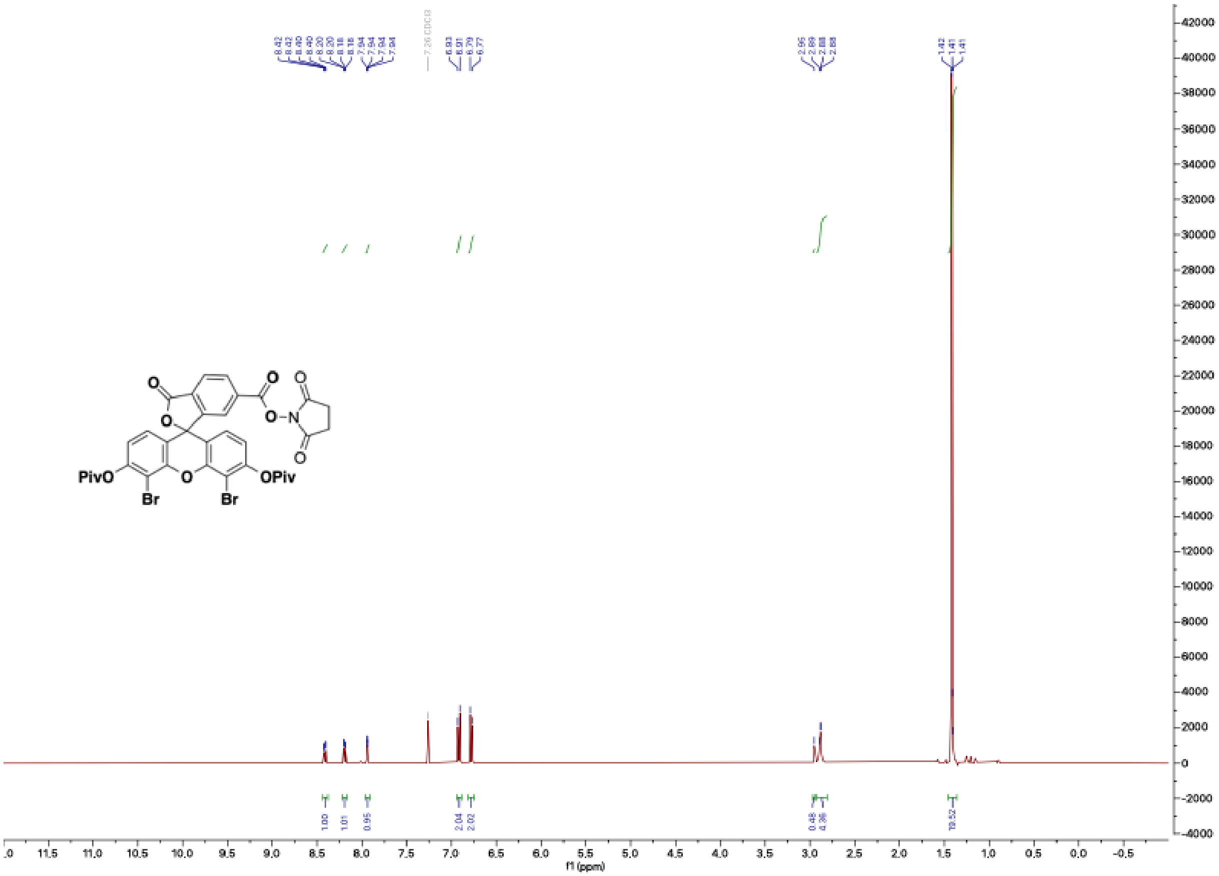

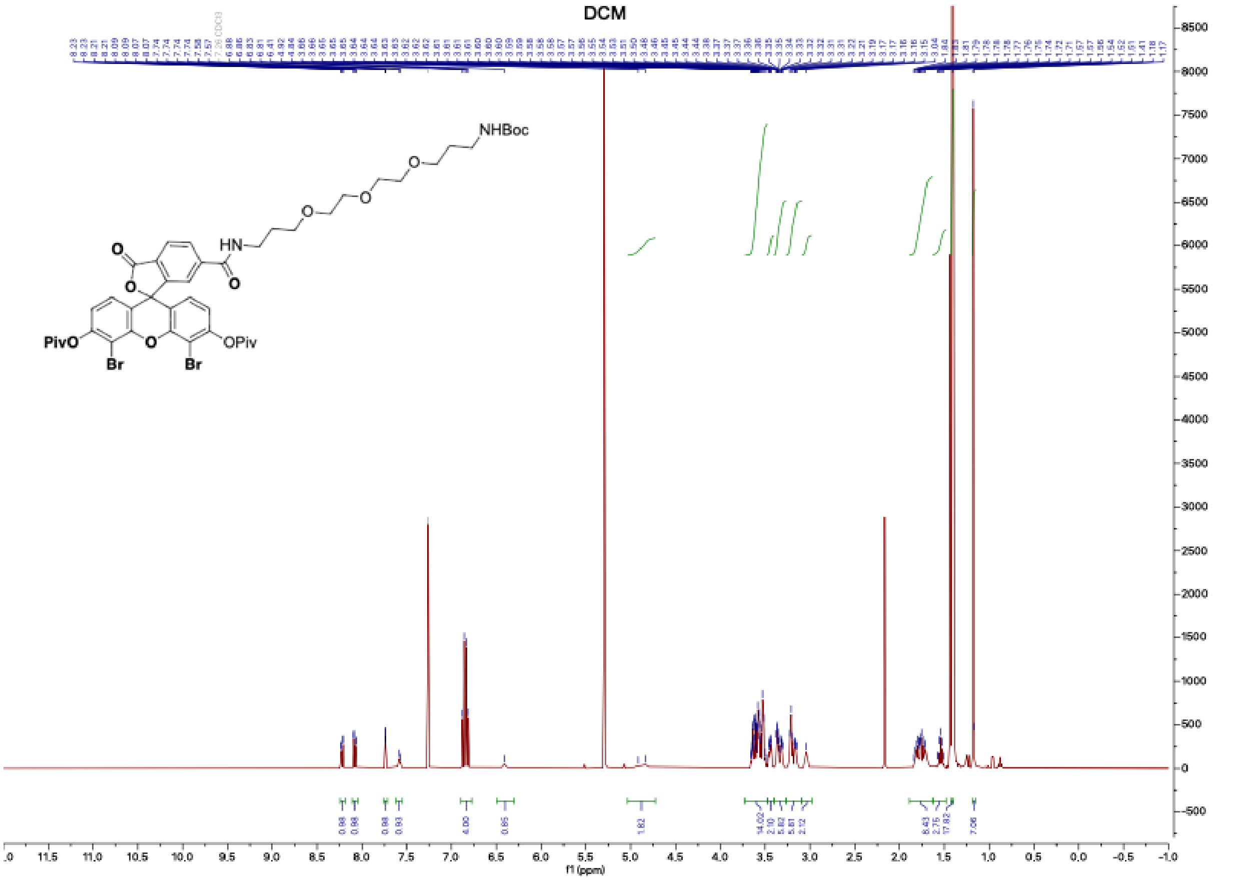
DCM
NHBoc
HN
PivO
OPiv
Br
Br
f1 (ppm)

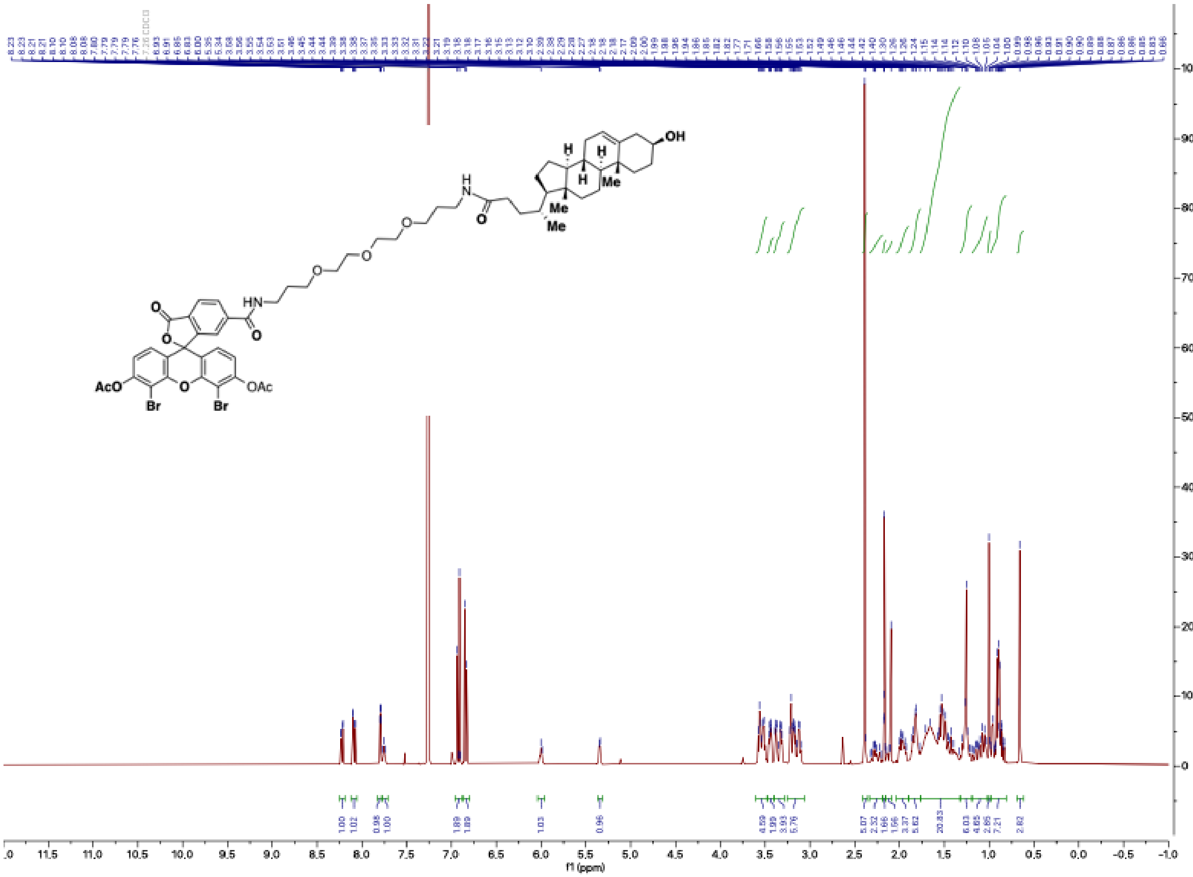

OH
H
H
H
Me
Me
Me
H
N
O
O
O
O
HN
O
O
O
AcO
OAc
Br
Br
O
f1 (ppm)

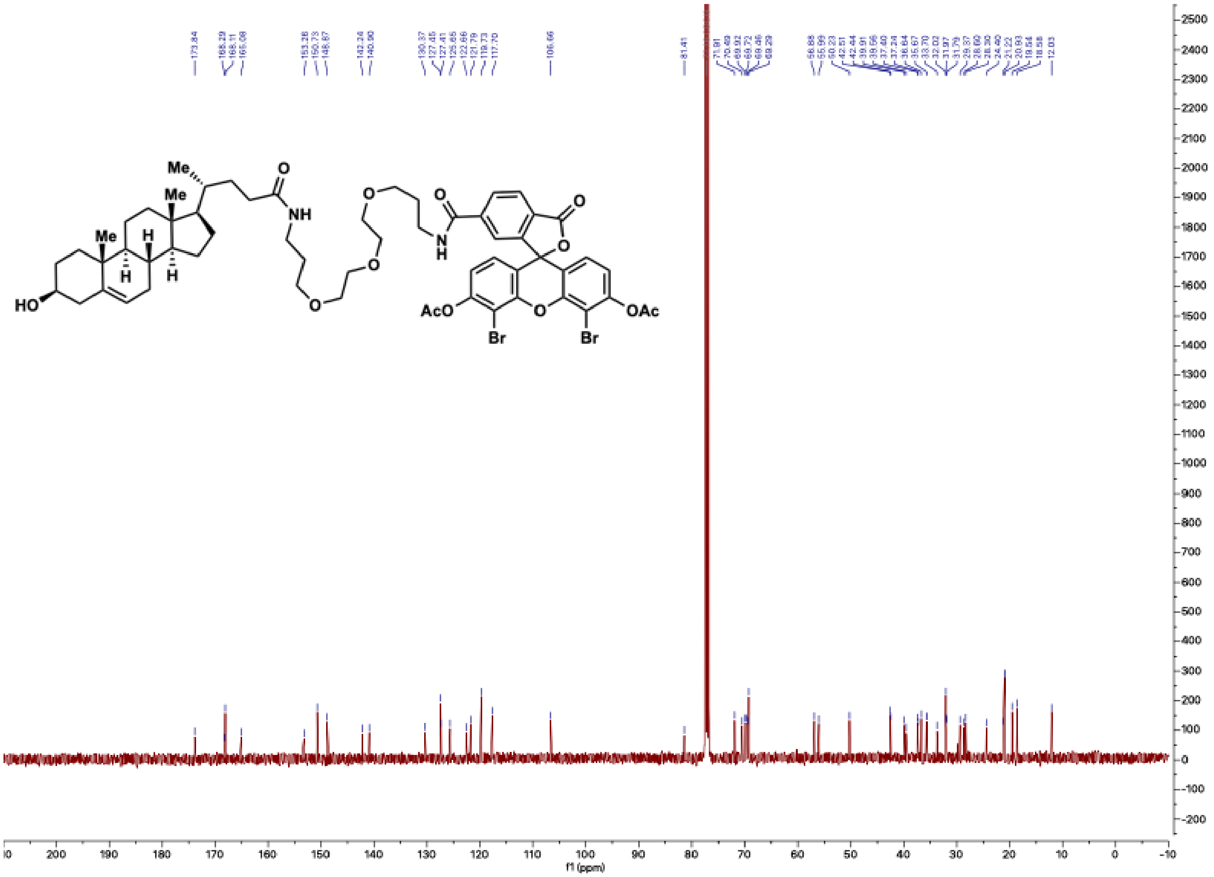

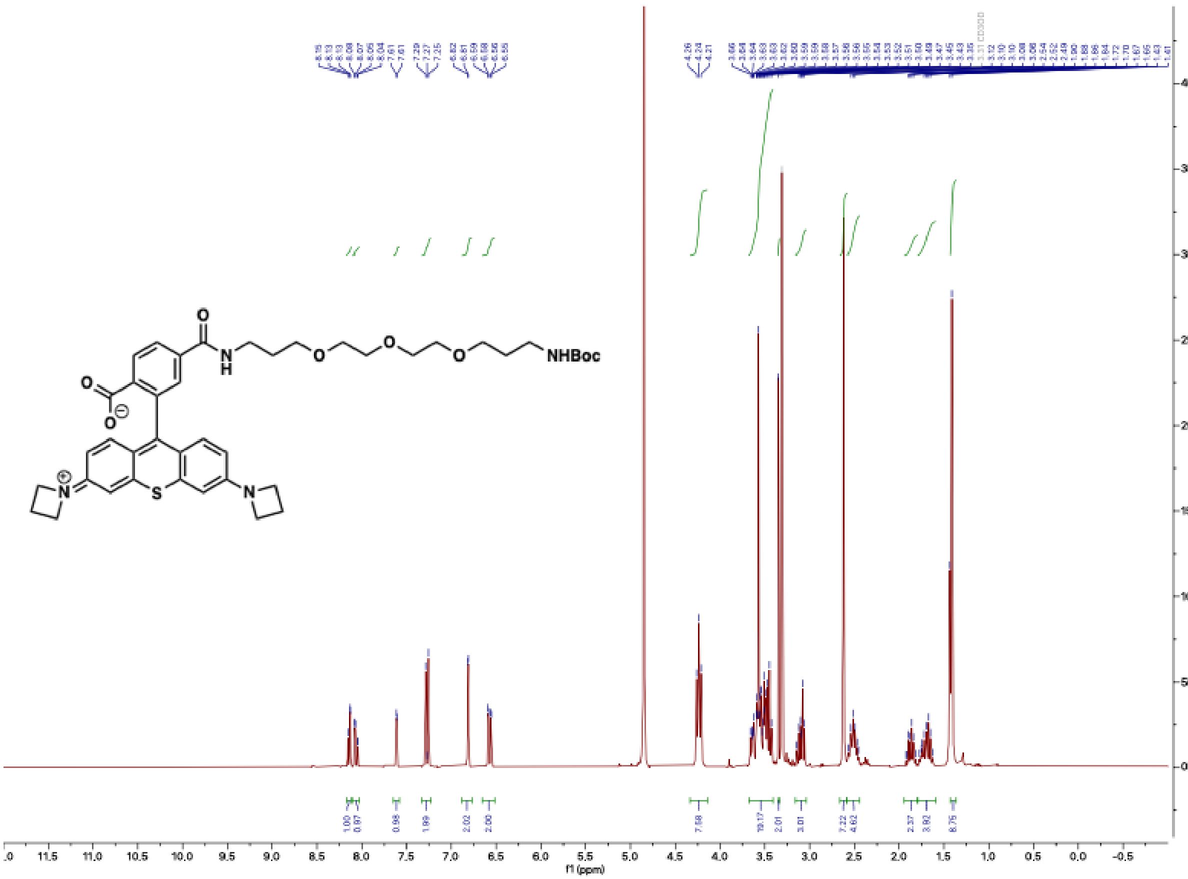

NHBoc
f1 (ppm)

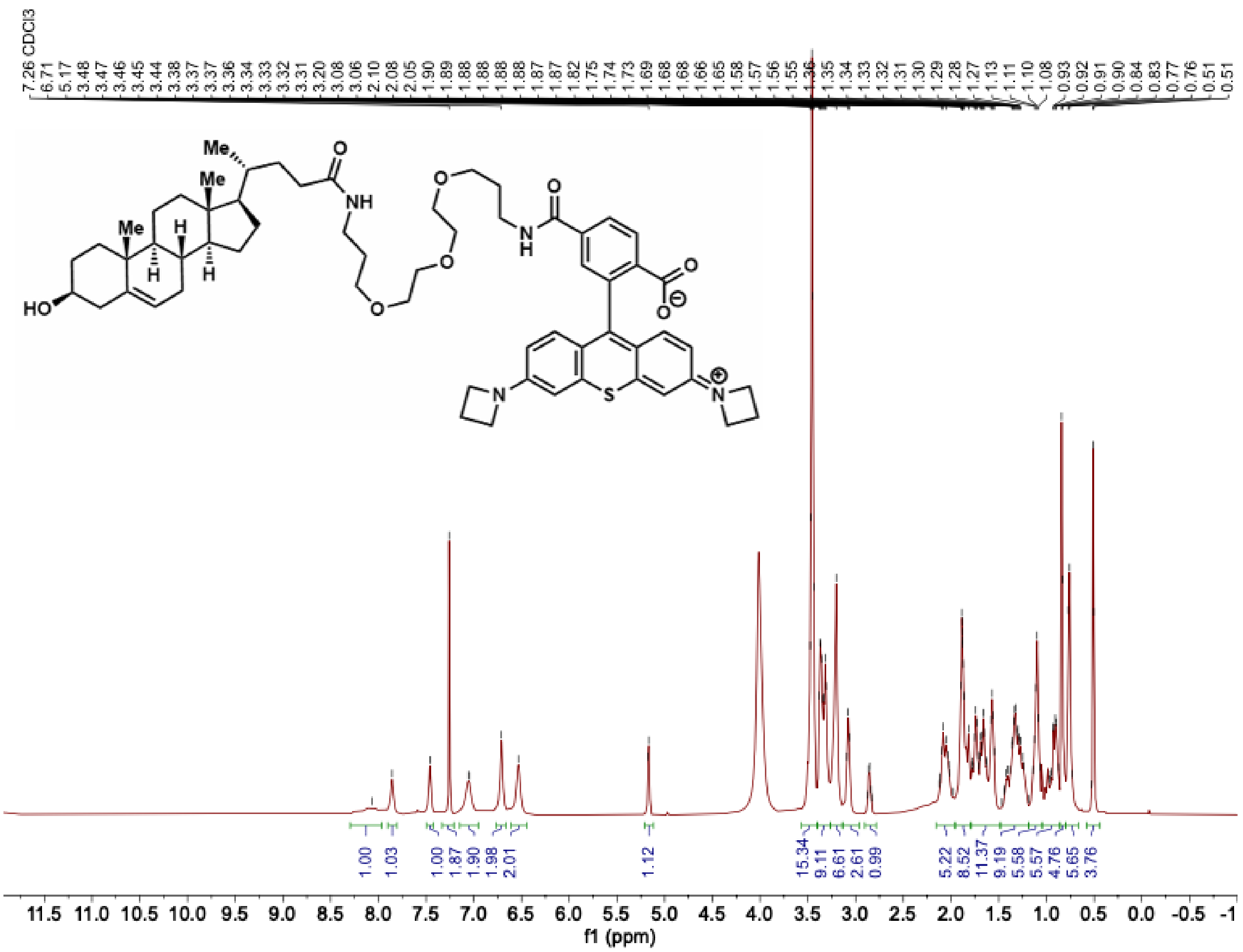

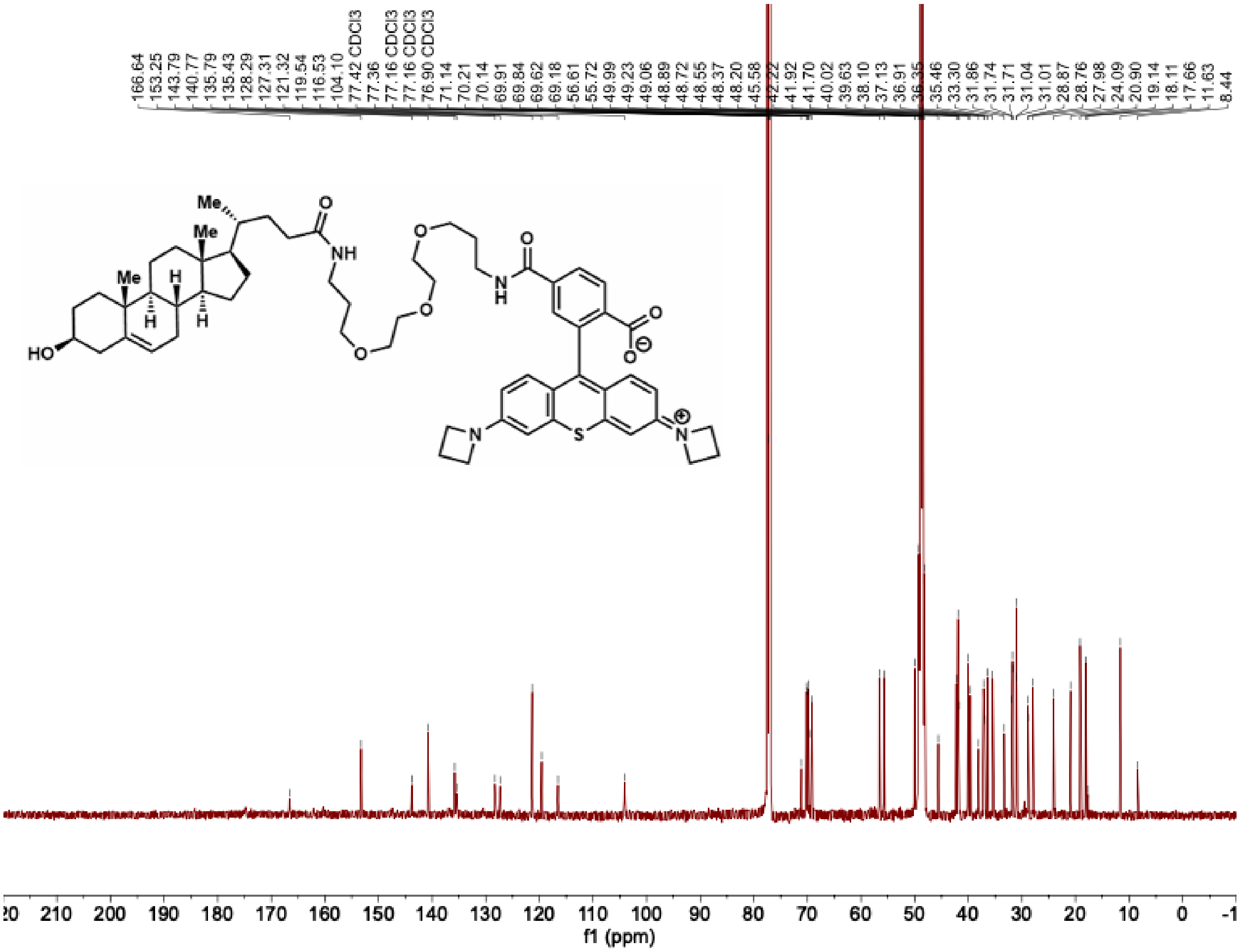
166.64
153.25
143.79
140.77
135.79
135.43
128.29
127.31
121.32
119.54
116.53
104.10
77.42 CDCl3
77.36
77.16 CDCl3
77.16 CDCl3
76.90 CDCl3
71.14
70.21
70.14
69.91
69.84
69.62
69.18
56.61
55.72
49.99
49.23
49.06
48.89
48.72
48.55
48.37
48.20
45.58
42.22
41.92
41.70
40.02
39.63
38.10
37.13
36.91
36.35
35.46
33.30
31.86
31.74
31.71
31.04
31.01
28.87
28.76
27.98
24.09
20.90
19.14
18.11
17.66
11.63
8.44
Me
Me
Me
H
H
H
HO
O
NH
O
O
O
O
N
H
O
O
N
S
N
f1 (ppm)

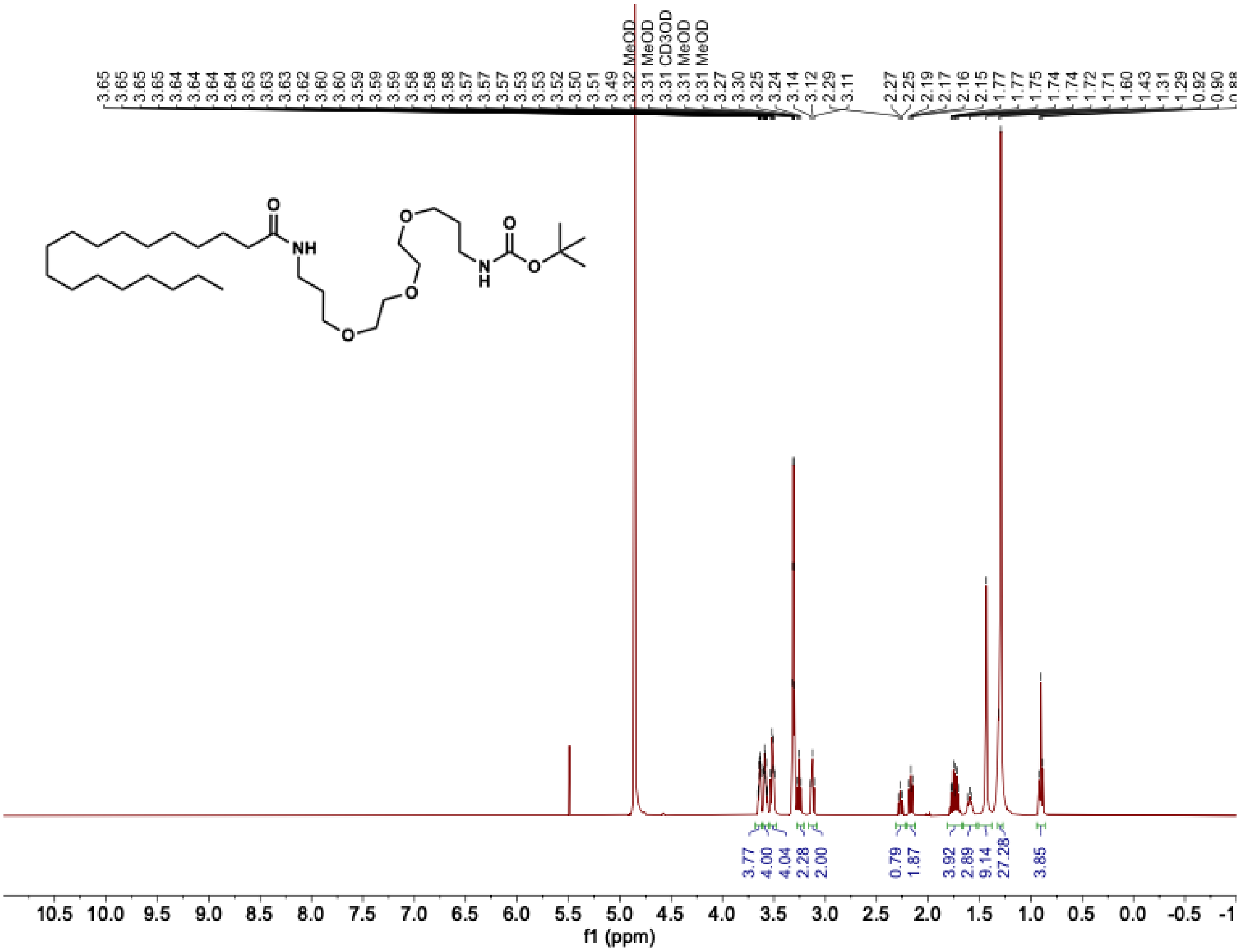
NH
O
O
O
N
H
O
O
MeOD
CD3OD
3.77
4.00
4.04
2.28
2.00
0.79
1.87
3.92
2.89
9.14
27.28
3.85
f1 (ppm)

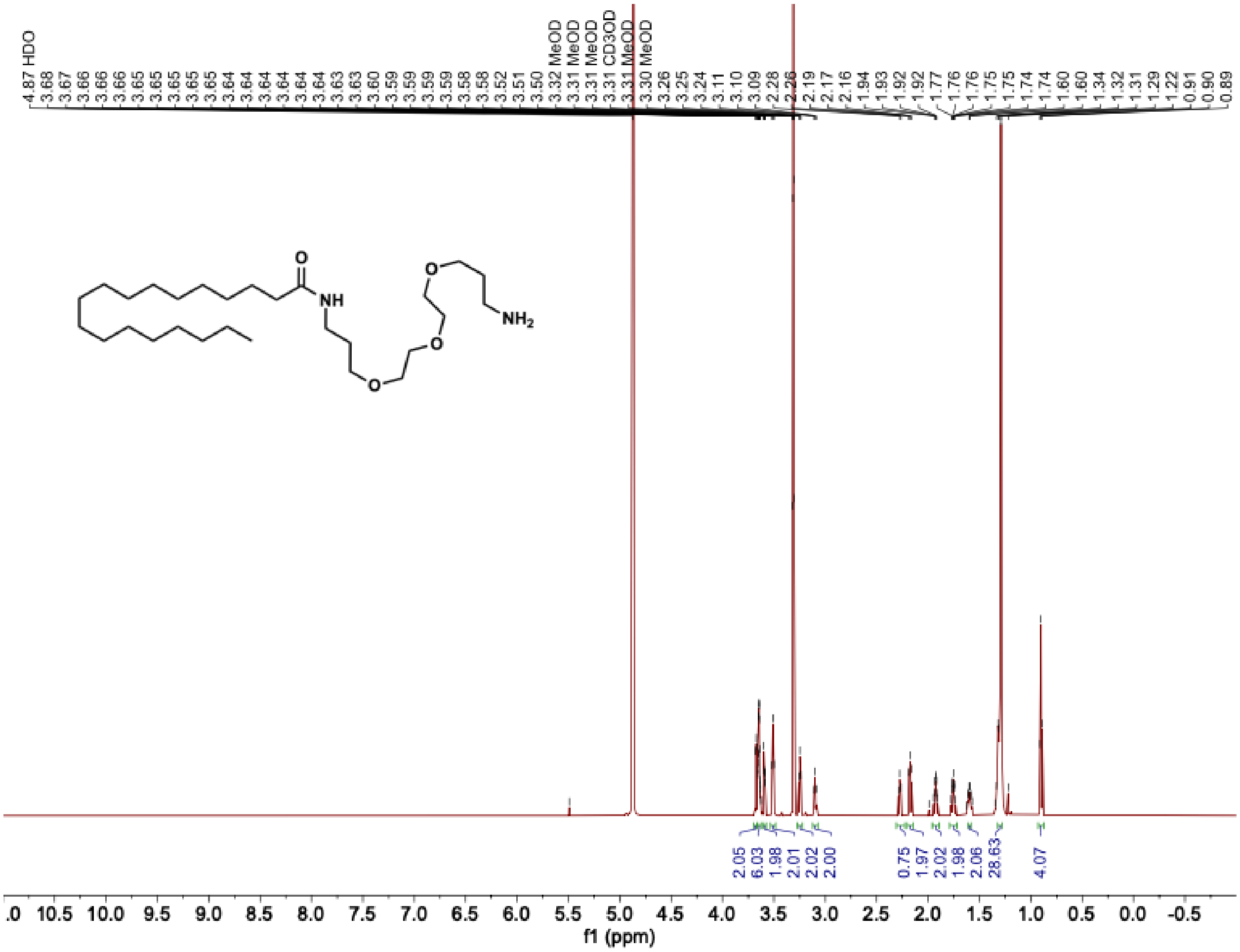
4.87 HDO
3.68
3.67
3.66
3.66
3.65
3.65
3.65
3.65
3.64
3.64
3.64
3.64
3.64
3.63
3.63
3.60
3.59
3.59
3.59
3.59
3.58
3.58
3.52
3.51
3.50
3.32 MeOD
3.31 MeOD
3.31 MeOD
3.31 CD3OD
3.31 MeOD
3.30 MeOD
3.26
3.25
3.24
3.11
3.10
3.09
2.28
2.26
2.19
2.17
2.16
1.94
1.93
1.92
1.92
1.77
1.76
1.76
1.75
1.74
1.74
1.60
1.60
1.34
1.32
1.31
1.29
1.22
0.91
0.90
0.89
O
NH
O
O
O
NH2
2.05
6.03
1.98
2.01
2.02
2.00
0.75
1.97
2.02
1.98
2.06
28.63
4.07
.0 10.5 10.0 9.5 9.0 8.5 8.0 7.5 7.0 6.5 6.0 5.5 5.0 4.5 4.0 3.5 3.0 2.5 2.0 1.5 1.0 0.5 0.0 -0.5
f1 (ppm)

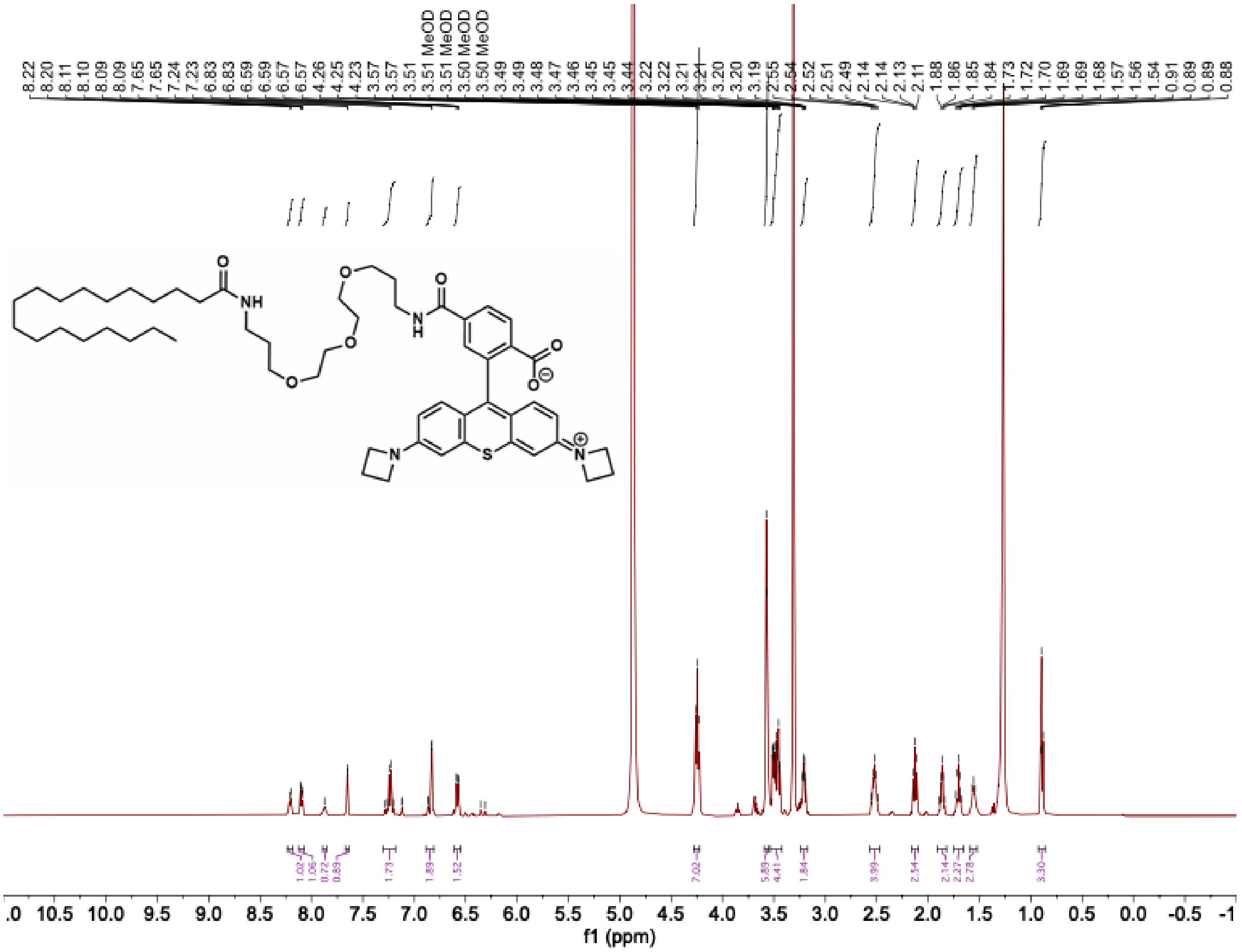

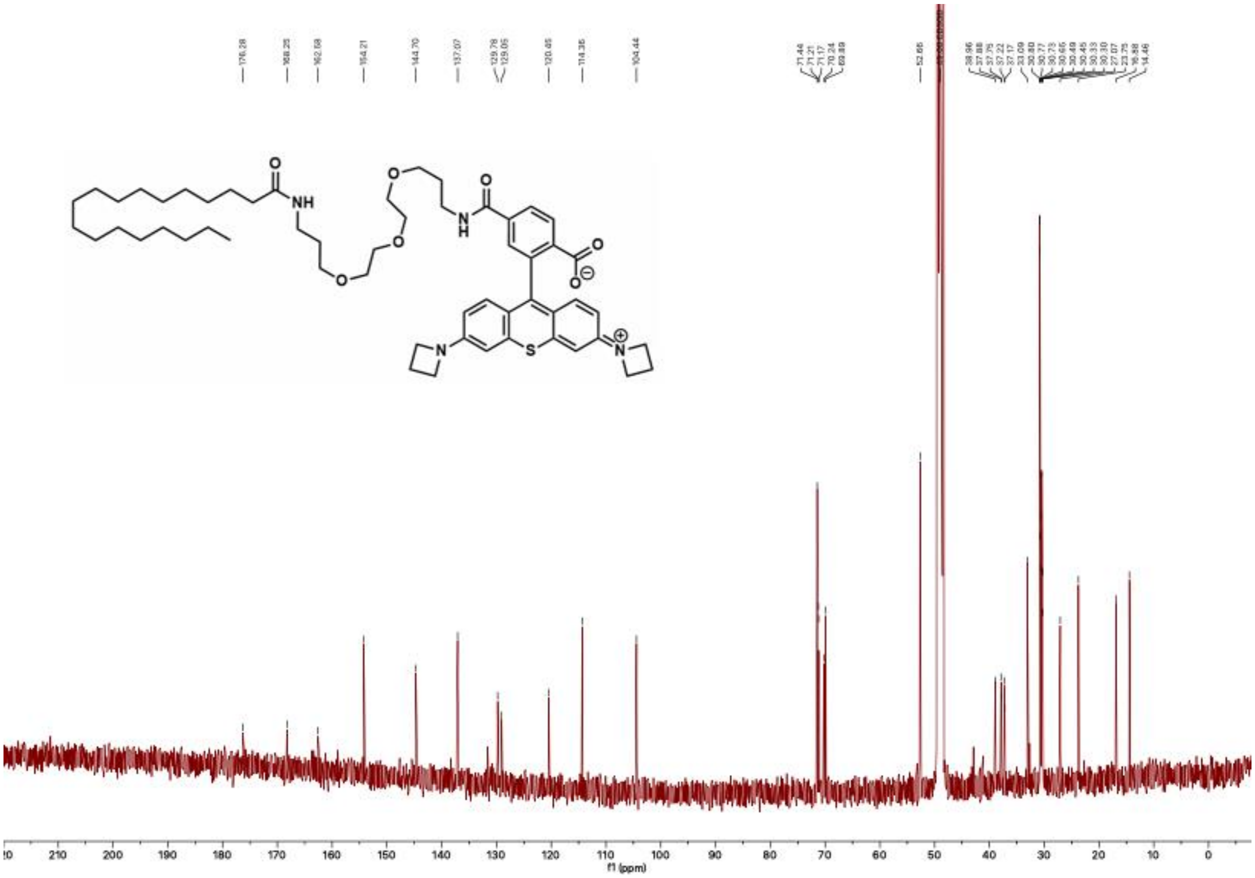
176.28
168.25
162.68
154.21
144.70
137.07
129.78
129.05
120.45
114.35
104.44
71.44
71.21
71.17
70.24
69.89
52.65
38.96
37.88
37.75
37.22
37.17
33.09
30.80
30.77
30.73
30.65
30.49
30.45
30.33
30.30
27.07
23.75
16.88
14.46
NH
O
N
H
S
N
f1 (ppm)

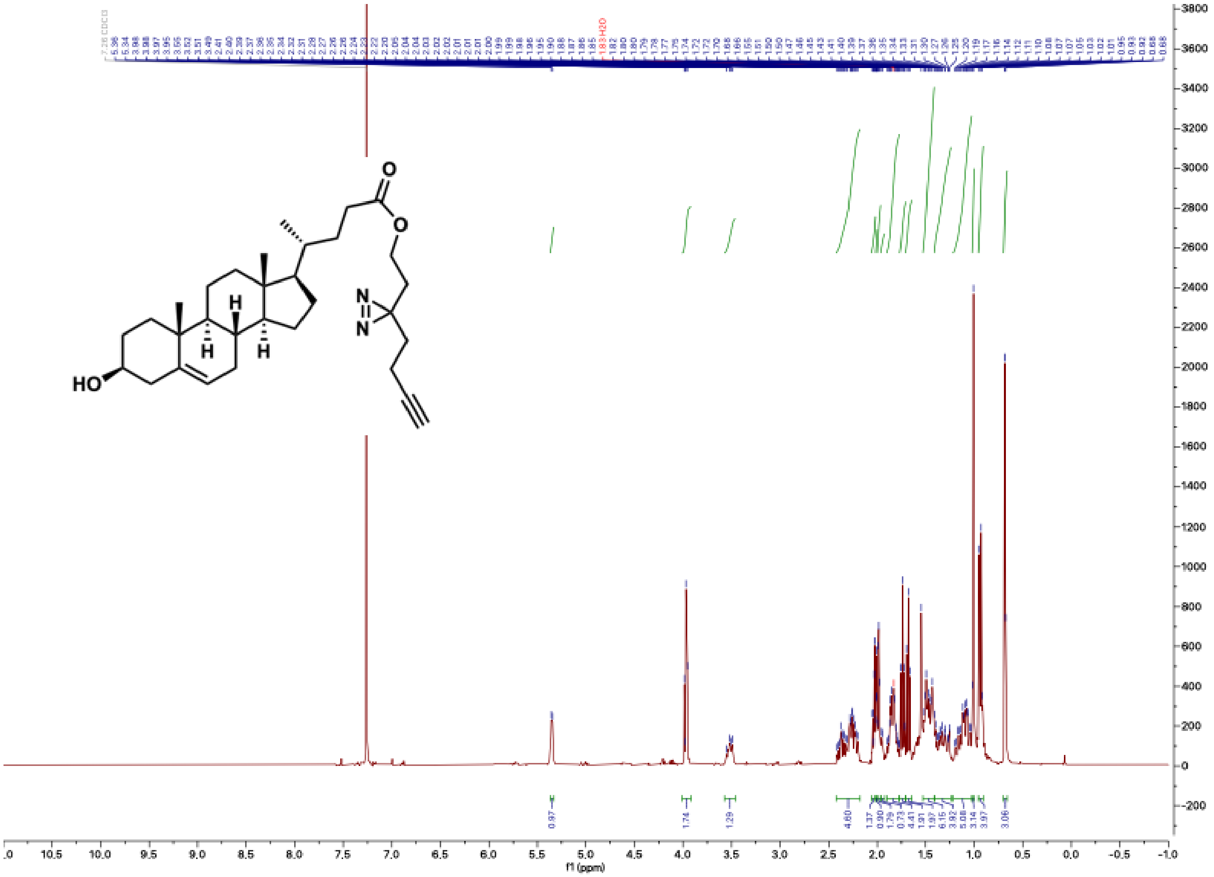

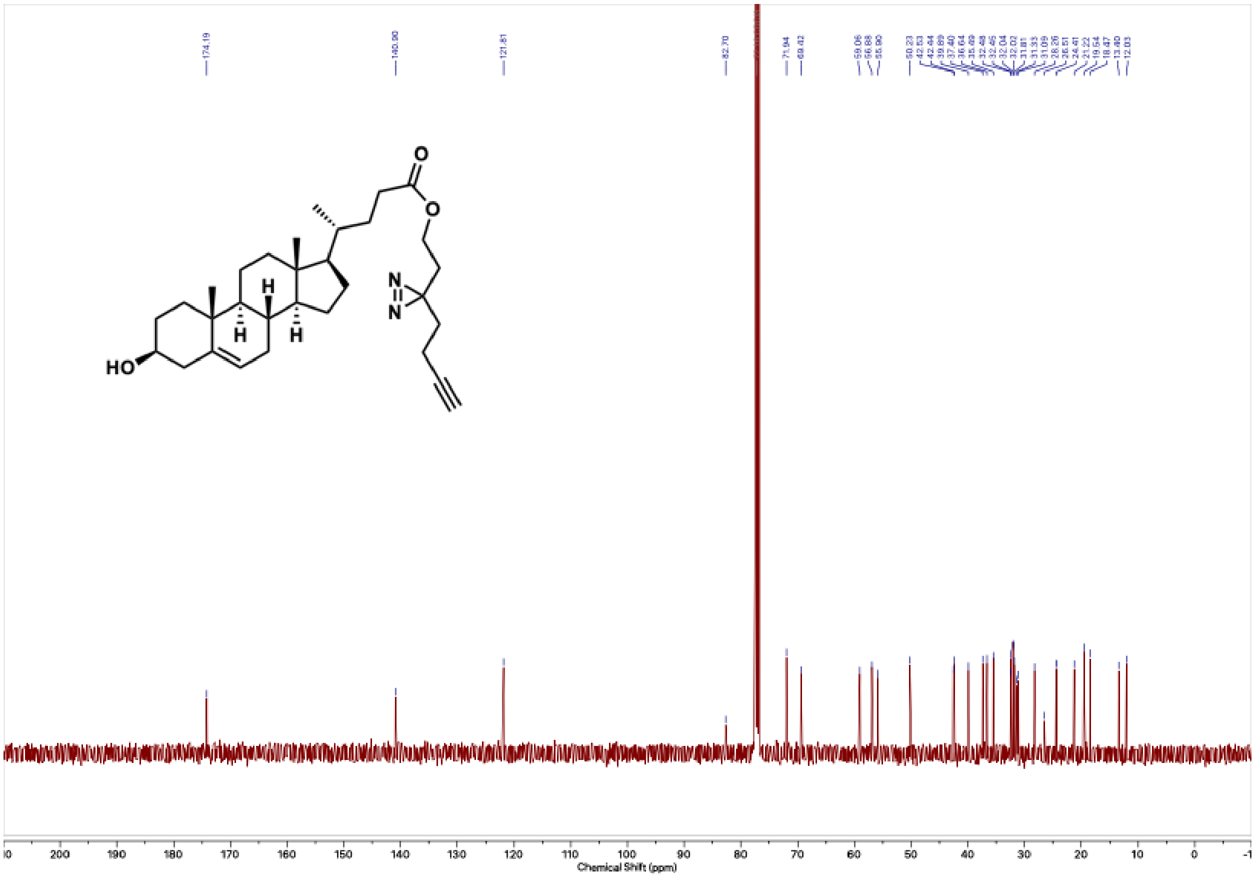
174.19
140.90
121.81
82.70
71.94
69.42
59.06
56.88
55.90
50.23
42.53
42.44
39.89
37.40
36.64
35.69
32.48
32.45
32.04
32.02
31.81
31.33
31.09
28.26
26.51
24.41
23.22
19.54
18.47
13.40
12.03
O
O
N
N
H
H
H
HO
200
190
180
170
160
150
140
130
120
110
100
90
80
70
60
50
40
30
20
10
0
Chemical Shift (ppm)
